# Single cell proteomics of tumor compartments identifies differential kinase activities defining sensitivity to mTOR-PI3-kinase inhibition

**DOI:** 10.1101/2021.01.06.425147

**Authors:** Nezihi Murat Karabacak, Yu Zheng, Taronish D. Dubash, Risa Burr, Douglas S. Micalizzi, Ben S. Wittner, Devon Wiley, Valentine Comaills, Erin Emmons, Kira Niederhoffer, Uyen Ho, Linda Nieman, Wilhelm Haas, Shannon L. Stott, David T. Ting, David T. Miyamoto, Daniel A. Haber, Mehmet Toner, Shyamala Maheswaran

**Author notes:** These authors contributed equally to this work (Nezihi Murat Karabacak, Yu Zheng).

## Abstract

Cancer therapy often results in heterogeneous responses in different metastatic lesions in the same patient. Inter- and intra-tumor heterogeneity in proteomic signaling within the various tumor compartments and its impact on therapy are not well characterized due to the limited sensitivity of single cell proteomic approaches. To overcome this barrier, we applied single cell mass cytometry with a customized 29-antibody panel [against cell states, receptor tyrosine kinases (RTK) and phosphoinositide 3-kinase/mammalian target of rapamycin (PI3K/mTOR)-, mitogen-activated protein kinase (MAPK)-, and cytokine-signaling] to PTEN-deleted orthotopic prostate cancer xenograft models to measure the evolution of kinase activities in different tumor compartments during metastasis and upon drug treatment. Compared with primary tumors and circulating tumor cells (CTCs), bone metastases but not lung and liver metastases exhibited elevated PI3K/mTOR signaling and RTKs including c-Met protein, which, when suppressed, impaired tumor growth in the bone. Intra-tumoral heterogeneity within tumor compartments also arises from highly proliferative EpCAM^high^ epithelial cells with increased PI3K and mTOR kinase activities co-existing with poorly proliferating EpCAM^low^ mesenchymal populations with reduced kinase activities, findings recapitulated in epithelial and mesenchymal CTC populations in metastatic prostate and breast cancer patients. Increased kinase activity in EpCAM^high^ cells rendered them more sensitive to PI3K/mTOR inhibition and drug resistant EpCAM^low^ populations with reduced kinase activity emerged over time. Taken together, single cell proteomics identified microenvironment- and cell state-dependent activation of kinase networks creating heterogeneity and differential drug sensitivity among and within tumor populations across different sites, defining a new paradigm of drug responses to kinase inhibitors.

## MAIN TEXT

Cancer cells initially confined to the primary site eventually disseminate through the blood and lymphatics to distant sites including the bone, lung, liver and brain. The spatially separated cancer cells in each metastatic site continue to evolve within the different microenvironments. This together with selection pressures imposed by drug treatment, leads to cellular and molecular heterogeneity among tumor cells^1–3^. Single cell RNA-sequencing of tumor cells and molecular analyses of circulating tumor DNA and circulating tumor cell (CTC) have been extensively used to interrogate these dynamic adaptations and to develop intervention and diagnostic tools^4, 5^. However, the poor sensitivity of current proteomic approaches have limited the mapping of signal transduction networks and post-transcriptional/translational events in single cells, creating a gap in our understanding of how the proteome evolves during cancer progression and therapeutic responses. Heterogeneity of protein expression in tumors is often evaluated using antibody-based staining of a few targets, and is confounded by the overlapping emission spectra of fluorophores^6^. Mass cytometry, which can simultaneously evaluate up to of 40 different epitopes^7^, is one of the few methodologies currently available to query and quantify larger proteomic signaling networks in single cells and in limiting amounts of tissue samples. Specially designed lanthanide-conjugated antibody panels against targets of interest have been used to collect quantitative multi-dimensional protein expression profiles to identify cellular lineages, proliferation and signaling programs, but most of these studies are primarily focused on the hematopoietic compartment^8–10^ as well as the intestine^11^, pancreas^12^, muscle^13^, brain^14^ where the input material is less limiting.

Inter- and intra-tumor heterogeneity in proteomic signaling networks across primary tumor, CTCs and metastatic tumors residing at various distal sites, and the influence of these site-dependent differences in signaling nodes (often activated and maintained through specific post-translational modifications) on tumor cell survival and drug responses are poorly defined. To address these questions, we applied single cell mass cytometry to two orthotopic PTEN-deleted mouse prostate cancer xenograft models and compared the activity of signaling nodes in the primary tumor, CTCs enriched from blood and in metastatic tumors residing in the lung, liver and bone, and during response to molecularly targeted therapy. These models are derived from orthotopic prostate injections of the androgen receptor negative PTEN-deleted human prostate cancer cell line, PC3^15^, and the PTEN-deleted mouse prostate tumor line, CE1-4, which recapitulates androgenic and epithelial features of primary human prostate cancer^16^. Both models display broad metastatic potential following orthotopic prostate injections. Single cells were interrogated by mass cytometry to measure the signaling nodes active within the different tumor compartments, and signaling changes following treatment with BEZ235, a dual inhibitor of PI3K and mTOR signaling^17^, were evaluated in the PC3 tumor model. The PTEN-deleted tumor xenograft models are particularly relevant since PTEN deletion and PI3K mutations occur in many cancers including prostate- and breast cancer, and inhibitors targeting the PI3K/mTOR signaling pathways present promising therapeutic targets against these cancers^18–21^.

A customized antibody panel against 29 proteins including specific phospho-proteins was used to simultaneously quantify cell lineage/cell state, signaling pathways, proliferation and apoptosis to identify signaling nodes active within tumor cells (**Fig. 1a**, **Extended Data Table 1**). Many of these antibodies have been extensively used in previously reported mass cytometry studies^8, 9, 22–28^, including those specifically marking epithelial tumor cells (EpCAM, and cytokeratin) and leukocytes (CD45), antibodies detecting signaling via PI3K/mTOR [phospho-S6 (pS6), phospho-4EBP1 (p4EBP1), phospho-Akt (pAkt; both p-Thr308 and p-Ser473), and phospho-GSK3β (pGSK3β)], MAPK [phospho-p38 (p-p38), phospho-ERK (pERK), phospho-JNK (pJNK), and phospho-p90RSK (p-p90RSK)], receptor tyrosine kinase (RTK; EGFR, HER2, HER3, IGFR and c-Met) and cytokine [phospho-Stat1 (pStat1), phospho-Stat3 (pStat3), and phospho-Stat5 (pStat5)] signaling pathways. We also included antibodies against critical transcription factors (activated ß-catenin and c-Myc), apoptosis (activated Caspase3) and proliferation (Ki67).

**Figure 1:**
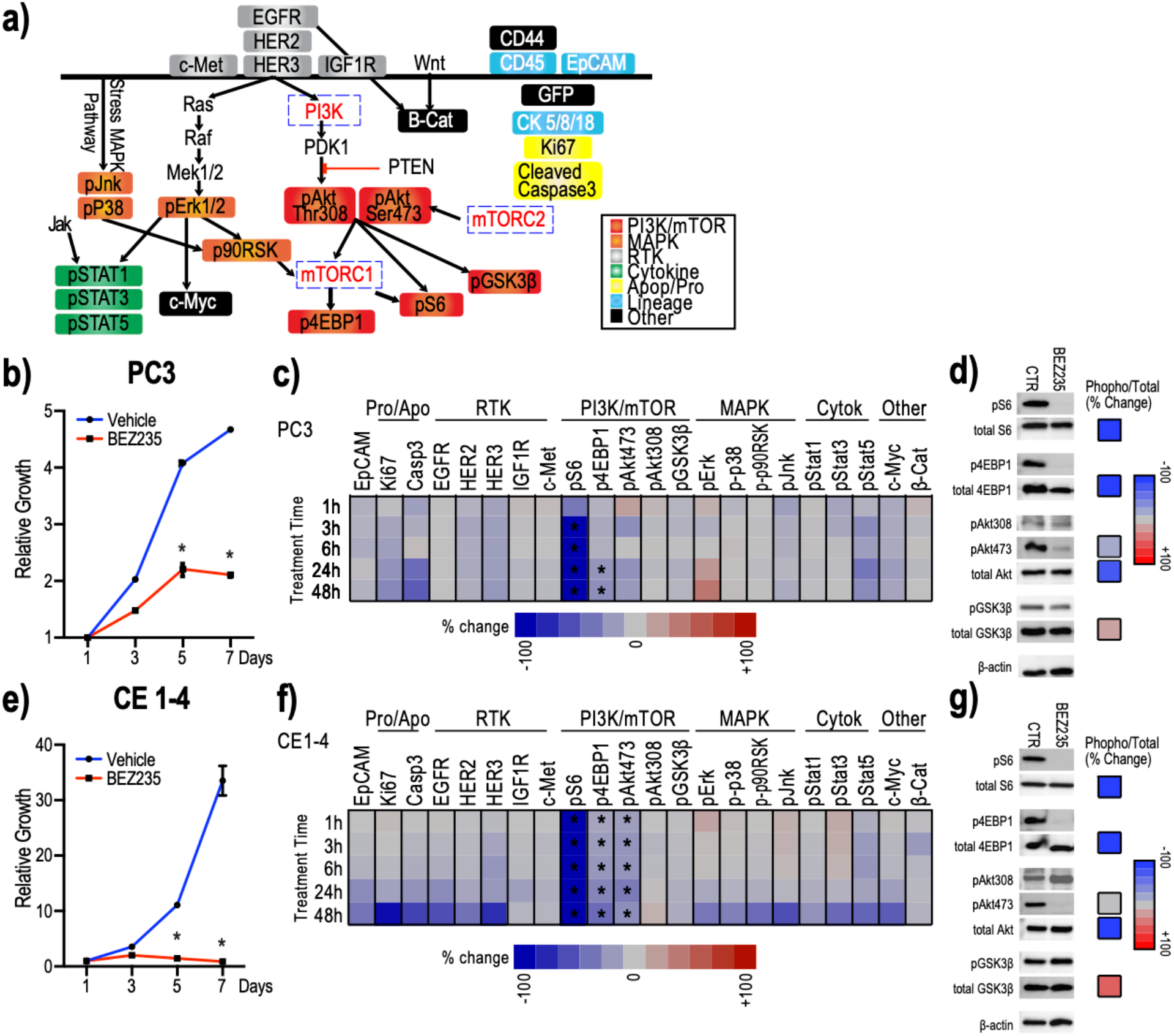
Antibody panel detects suppression of PI3K/mTOR activity in individual PC3 and CE1-4 cells treated with BEZ235 *in vitro*. 1a: The customized 29-antibody panel used to detect and quantify protein and phospho-protein levels in single cells by mass cytometry include PI3K/mTOR-, MAPK-, and cytokine-signaling, RTKs as well as apoptosis, proliferation, lineage and other (c-Myc, β-catenin, CD44 and GFP) markers. 1b: BEZ235 treatment suppresses the growth of PC3 cells *in vitro*. Untreated cells are shown as control (*q<0.1). 1c: Heat map showing mean change in the signaling proteins within each of the indicated pathways in PC3 cells treated with BEZ235 *in vitro*, compared with untreated control cells (n=4 experimental replicates). Significant changes are marked with asterisks (q<0.1 by multiple t-tests FDR correction). 1d: Western blots confirm changes observed by mass cytometry of single PC3 cells. Western blots from BEZ235 or DMSO-treated PC3 cells (left) and percent change in each signaling epitope (phospho-protein/total protein) (right) are shown. 1e: BEZ235 treatment suppresses the growth of CE1-4 cells *in vitro*. Untreated cells are shown as control (*q<0.1). 1f: Heat map showing mean change in the signaling proteins within each of the indicated pathways in CE1-4 cells treated with BEZ235 *in vitro*, compared with untreated control cells (n=4 experimental replicates). Significant changes are marked with asterisks (q<0.1 by multiple t-tests FDR correction). 1g: Western blots confirmed changes observed in single CE1-4 cells by mass cytometry. Western blots from BEZ235 or DMSO-treated CE1-4 cells (left) and percent change in each signaling epitope (phospho-protein/total protein) (right) are shown.

We first tested this 29-antibody panel *in vitro* using PC3 cells treated with BEZ235^17^. BEZ235 significantly reduced the growth of PC3 cells *in vitro* (**Fig. 1b**; q<0.1). Following drug exposure for 1, 3, 6, 24 and 48 hours, a total of more than 650,000 individual PC3 cells were analyzed across four independent experimental replicates. Consistent with BEZ235 inhibiting the PI3K/mTOR signaling axis^17^, both pS6 and p4EBP1 (compared with untreated control) were significantly inhibited (**Fig. 1c**; q<0.1). BEZ235 also reduced pAkt473 but not pAkt308 levels. Western blot analysis validated these findings showing that BEZ235 treatment of PC3 cells specifically suppressed pS6 and p4EBP1 *in vitro*. The drug did not suppress total Akt levels but, consistent with the mass cytometry data, suppressed pAkt473 and not pAkt308 (**Fig. 1d**). Single cell proteomic analysis and western blots of CE1-4 treated with BEZ235 *in vitro* further confirmed these findings showing drug induced suppression of growth as well as pS6, p4EBP1 and pAkt473 (**Fig 1e-g**; q<0.1). Single cell mass cytometry analysis of CE1-4 cells showed that the PI3K inhibitor, GDC0941^29^ suppressed the downstream targets pS6, pAkt308/473, and pGSK3β (**Extended Data Fig. 1a**; p<0.01).

To further confirm the reliability and specificity of this antibody panel, we analyzed the BEZ235-treated breast cancer patient-derived CTC culture, BRX142, harboring mutant PI3K; BEZ235 robustly suppressed the PI3K/mTOR pathway intermediates, pS6, p4EBP1and pAkt473 in BRX142 cells *in vitro* (**Extended Data Fig. 1b**; p<0.01). Taken together, these results confirm and validate the specificity of the customized 29 antibody panel to detect changes in kinases mediating PI3K/mTOR signaling in cells.

Having validated the specificity of the antibody panel using multiple *in vitro* cell culture models, we applied it to interrogate and quantify the differences in signaling pathways in tumor cells growing at different sites in mouse models. Single tumor cells, collected following collagenase dissociation of primary and metastatic tumors growing in the lung, liver and bone, were identified based on GFP, cytokeratin, and nuclear label (Ir-intercalator) positivity and CD45 negativity (**Extended Data Fig. 2a**). For CTC isolation, mouse blood was collected into tubes containing preservatives to stabilize the phospho-epitopes on the proteins. After incubating with immunomagnetic bead-conjugated antibodies against white blood cells, the blood was processed through the CTC-iChip, a microfluidic device that uses negative selection to enrich CTCs^30, 31^. Over 85% of the cells enriched from the various compartments following enzymatic digestion were intact/live as measured by ^103^Rh-intercalator labeling (**Extended Data Fig. 2a**). Osmium labeling intensities showed no significant size differences in the tumor cells isolated from the various tissue compartments (**Extended Data Fig. 2b**).

Dimensional reduction of all epitopes of interest (excluding CD44, CD45, CK, and GFP) using the viSNE method^32^, showed that the primary PC3 tumor cells are highly heterogeneous (**Fig. 2a, Supplementary Data 1**) with some cells overlapping with the CTCs and the metastatic tumor cells enriched from the lung and liver. Bone metastatic tumor cells in the PC3 tumor model clustered separately on the viSNE plot implying unique molecular properties (**Fig. 2a**). Quantification of the mean expression level of each of the epitopes within all the single cells collected from the different tissue compartments (per mouse) (**Fig. 2b**) showed that the bone metastatic cells, compared with primary tumor, CTCs and lung and liver metastasis, expressed significantly higher levels of the proliferation marker, Ki67, PI3K/mTOR - (pS6, p4EBP1, pAkt473, pAkt308, pGSK3β), and MAPK- (pErk and p-p90RSK) signaling intermediates as well as the RTKs, EGFR and c- Met (**Fig. 2b**; q- and p-values for each epitope provided in the figure legend). The PI3K/mTOR signaling index, measured using the sum of standardized pS6, p4EBP1, pAkt473, pAkt308 and pGSK3β channels, was significantly higher in the single cells collected from bone metastasis compared with tumor cells enriched from all the other sites (**Fig. 2c**; q<0.0001). These findings were further confirmed by immunohistochemical staining of primary tumor and bone metastases with antibodies against c-Met, pS6 and p4EBP1, all of which show significantly increased levels in bone metastases compared with matched primary tumors (**Extended Data Fig. 3**; c-Met: p=0.0053; p4EBP1: p=0.0019; pS6 p=0.0386). Increased c-Met and EGFR expression in bone metastases compared with primary prostate cancer, identified using the single cell mass cytometry, is consistent with previously reported findings^33, 34^. This serves as additional confirmation of the specificity of the antibody panel used for single cell proteomic analysis of prostate cancer cells residing within the various tissue compartments.

**Figure 2:**
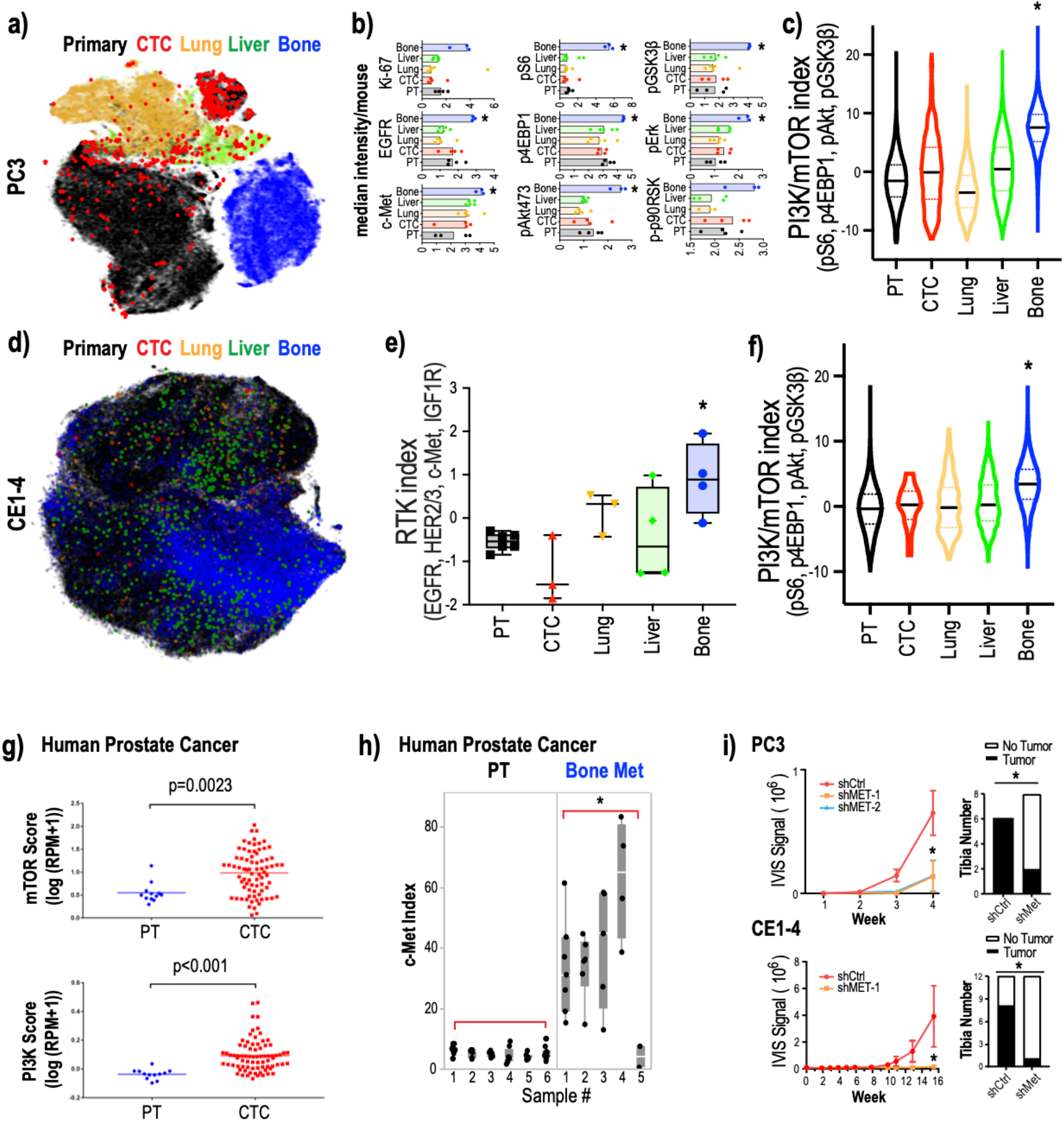
Bone metastases growth requires c-Met activity 2a: Dimensional reduction of the expression of all analyzed epitopes in single tumor cells in the PC3 mouse tumor metastasis model using viSNE shows a large fractured island consisting of dispersed primary tumor cells (black) and CTCs (red), and liver (green), lung (yellow) and bone (blue) metastatic cells. 2b: Signaling intermediates of the PI3K/mTOR- and MAPK-pathways and RTKs are elevated in bone metastases. Median epitope levels across single tumor cells isolated from the primary tumor and lung, liver and bone metastases as well as CTCs per mouse are shown in box plots. Each dot represents the median intensity of the epitope quantified (across single cells from each of the tumor compartment) for a single mouse. Ki67 q=0.3088/p=0.0702, pS6, pGSK3 q=0.002/p=0.0007, EGFR q=0.0098/p=0.0047, p4EBP1 q<0.0001/p<0.0001, pErk q<0.0001 p<0.0001, c-Met q=0.0294/p=0.0.0094 by one-way ANOVA, corrected for multiple comparisons by FDR, Benjamini, Krieger and Yekutieli. 2c: PI3K/mTOR index (measured as the sum of standardized pS6, p4EBP1, pAkt and pGSK3β levels) was significantly elevated in PC3 bone metastatic cells compared to primary tumor cells, CTCs, and metastatic cells in other sites. q<0.0001 by one-way ANOVA, corrected for multiple comparisons by FDR, Benjamini, Krieger and Yekutieli, whereas MAPK- and cytokine-signaling pathway signatures were not significantly different (**Extended Data Fig. 13**). 2d: viSNE of all analyzed epitopes in single tumor cells from the CE1-4 mouse prostate cancer model showing primary tumor cells (black), CTCs (red), and liver (green), lung (yellow) and bone (blue) metastatic cells. 2e: RTK index (measured as the sum of standardized EGFR, HER2, HER3, c-Met and IGF1R levels) is significantly elevated in CE1-4 bone metastatic cells compared to primary tumor cells, CTCs, and metastatic cells in other sites. q<0.0001 by one-way ANOVA, corrected for multiple comparisons by FDR, Benjamini, Krieger and Yekutieli. 2f: PI3K/mTOR index (measured as the sum of standardized pS6, p4EBP1, pAkt and pGSK3β levels) is significantly elevated in CE1-4 bone metastatic cells compared to primary tumor cells, CTCs, and metastatic cells in other sites. q<0.0001 by one-way ANOVA, corrected for multiple comparisons by FDR, Benjamini, Krieger and Yekutieli. 2g: mTOR and PI3K activities are elevated in CTCs isolated from the blood of mCRPC patients^35^. Gene expression signatures associated with mTOR and PI3K activation are significantly higher in single cell RNA-sequencing data derived from 77 individual CTCs collected from 13 mCRPC patients (mean 6 CTCs per patient), compared with RNA-sequencing of 12 primary prostate cancers at single cell resolution. p<0.0023 and p<0.001 respectively, by Mann Whitney test. 2h: c-Met protein expression is significantly elevated prostate cancer bone metastases compared with primary tumors. Quantification of c-Met protein expression by IHC of primary human prostate cancer (n=6) and prostate cancer bone metastases (n=5). p=0.0066 by t-test. 2i: Suppression of c-Met impairs prostate tumor growth in the bone. c-Met depleted (using shRNAs against c-Met) and control PC3 (1000 cells each) and CE1-4 (250 cells each) were injected into the tibia of mice and their growth was monitored over 4 weeks and 16 weeks, respectively. Left panel showing bioluminescence signal over time and right panel showing number of tibiae with and without tumor growth observed at the final time point. p<0.01 by t-test. c-Met knockdown impaired tumor growth in the bone in both models. p=0.0188 for PC3, p=0.0114 for CE1-4 by t-test.

To further validate these findings, we evaluated a second orthotopic mouse prostate cancer model developed from the PTEN-deleted mouse prostate tumor cells, CE1-4^16^. We applied single cell proteomic analysis following the same procedures as described for the PC3 model. The antibodies used to detect EpCAM, c-Met and CK, however, were switched to specifically detect mouse-specific epitopes. We analyzed individual tumor cells enriched from the CE1-4 primary tumor, CTCs, and metastatic tumors collected from the lung, liver and bone (n=4 mice). Consistent with the PC3 tumor model, metastatic CE1-4 cells growing in the bone but not in the lung and the liver showed significantly increased RTKs (the sum of standardized values for EGFR, HER2/3, c-Met and IGF1R) and PI3K/mTOR signaling index (the sum of standardized values for pS6, p4EBP1, pAkt473, pAkt308, pGSK3β) compared with the primary tumor and CTCs (**Fig. 2d-2f**; q<0.0001).

We then wanted to determine whether c-Met and PI3K/mTOR signaling is also elevated in bone metastases in human prostate cancer patients. Prostate cancer predominantly metastasizes to the bone, which cannot be readily biopsied. As such, CTCs in the blood of metastatic castration resistant prostate cancer (mCRPC) patients, in whom the primary tumor has been controlled, serve as proxy for cancer cells residing within the bone. Analysis of single cell RNA-sequencing data from 77 CTCs collected from 13 mCRPC patients (a mean of six CTCs per patient and these CTCs harbor androgen receptor mutations and splice variants as previously described in^35^; PTEN/PI3K mutation status not available) showed that the PI3K and mTOR gene expression signatures are indeed significantly elevated in the CTCs compared with the primary tumors (**Fig. 2g**; primary tumors n=12 patients; PI3K: p<0.001; mTOR: p=0.0023). c-Met mRNA was not elevated in prostate CTCs compared with the primary tumor cells (**Extended Data Fig. 4a**); analysis of multiple publicly available datasets also showed that c-Met RNA is not upregulated in metastatic prostate cancer compared with primary tumors (**Extended Data Fig. 4b**). However, c-Met protein expression, evaluated using immunostaining, was higher in four out of five human bone metastatic prostate cancers compared with primary tumors (n=6) (**Fig. 2h**; p=0.0066; **Extended Data Fig. 5**) suggesting that c-Met protein expression is elevated in prostate cancer bone metastases most likely through post-transcriptional mechanisms.

To determine whether elevated c-Met protein is essential for the survival and growth of prostate cancer cells in the bone, we knocked down c-Met expression in both PC3 and CE1-4 cells using two different shRNAs against c-Met. Knockdown of c-Met expression does not change the proliferation rate of the PC3 and CE1-4 cells *in vitro* compared to control cells (**Extended Data Fig. 6a and 6b**). However, depletion of c-Met significantly impairs both PC3 and CE1-4 tumor growth in the bone microenvironment when inoculated into the mouse tibia (**Fig. 2i**; PC3: p=0.0188; CE1-4: p=0.0114) suggesting the c-Met is essential for prostate tumor cell growth in the bone. Taken together, these findings show that tumors residing within different compartments, despite originating from the same primary tumor, exhibit differential kinase activities and growth dependencies, which may provide a mechanistic explanation for heterogeneous responses to therapies that can occur across different metastatic sites.

We then characterized drug-induced changes in tumor burden and in signaling pathways across tumors growing within the different tissue compartments. Mice bearing two week-old GFP and luciferase-labeled, orthotopic PC3 tumors were treated with BEZ235^17^ for 9 to 10 weeks (**Fig. 3a**; n=4 mice for untreated control, n=5 mice for BEZ235 treatment). Growth was monitored by IVIS live imaging. BEZ235 significantly suppressed PC3 tumor burden in mice (**Fig. 3b**; p-values shown in figure legends). To determine signaling changes within the different tumor deposits, single cells were collected from primary tumors and lung, liver and bone metastases of untreated and drug treated mice at the end of the experiment. CTCs were enriched from mouse blood. While 75% of mice in the untreated group presented with bone metastases, none of the BEZ235-treated mice (0%) had bone metastases (**Fig. 3c**, p=0.047). Consistent with this observation, no tumor cells were evident in the bones of five BEZ235-treated mice (0%) whereas we were able to collect a total of >20K bone metastatic cells from untreated animals.

**Figure 3:**
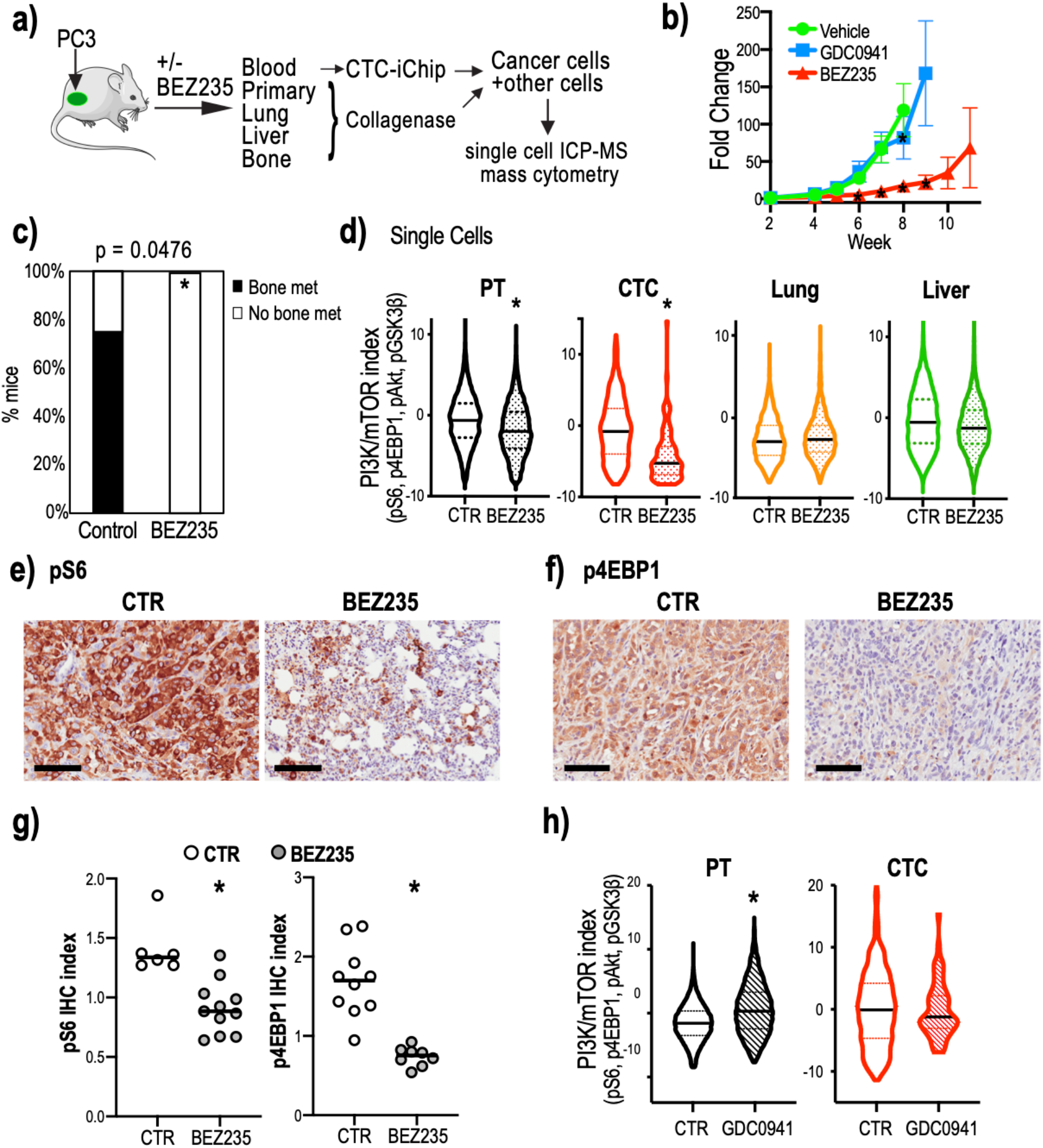
Dual PI3K/mTOR inhibitor treatment shows heterogeneous drug responsiveness of tumor deposits. 3a: Mouse model outlining the experiment to collect single cells from the primary and metastatic tumors and CTCs from untreated and BEZ235-treated mice harboring orthotopic PC3 tumor xenografts. CTCs were enriched from the mouse blood processed through the CTC-iChip. 3b: GFP and luciferase labeled PC3 cells were injected into the prostates of immune-compromised mice and 2-week old tumors were treated with BEZ235 or GDC0941 (n=6 mice for BEZ235; n=4 mice for GDC0941). BEZ235-treatment significantly suppressed tumor growth (week 5 p=0.3571, week 6 p=0.0032, week 7 p<0.0001, week 8 p<0.0001, two-way ANOVA, Dunnett correction). GDC0941 had no effect on tumor growth (week 5 p=0.9579, week 6 p=0.4257, week 7 p=0.9294, week 8<0.001, two-way ANOVA, Dunnett correction). Untreated tumor growth is shown as control (n=7 mice). 3c: BEZ235-treatment eliminates tumor growth in the bone. The bar graph shows that the percentage of mice with bone metastases in the untreated group is significantly higher than in the BEZ235-treated group (p=0.047 by Two-tailed Fisher’s exact test). 3d: PI3K/mTOR indices (measured as the sum of standardized pS6, p4EBP1, pAkt and pGSK3β levels) of single tumor cells enriched from the primary tumors, CTCs and lung and liver metastases collected from mice untreated or treated with BEZ235. PI3K/mTOR indices in primary tumor cells and CTCs were significantly decreased upon drug treatment (p<0.0001 by Mann-Whitney test); lung and liver metastases showed no significant change upon BEZ235 treatment. 3e-3g: pS6 and p4EBP1 staining of primary tumors resected from untreated control (CTR) and BEZ235-treated animals. Representative images are shown in **Fig. 3e and 3f**. Scale bar represents 100 µm. Quantification of staining indices in **Fig. 3g** shows that BEZ235 treatment significantly reduced pS6 and p4EBP1 levels in primary tumors (n=3 mice and n=17 tumor areas for pS6, n=2 mice and n=18 tumor areas for p4EBP1). p<0.001 for p4EBP1, p=0.0011 for pS6 by Mann-Whitney test. 3h. PI3K/mTOR indices primary tumor-derived cells and CTCs treated with the PI3K inhibitor, GDC0941. Untreated controls are shown (CTR). GDC0941 significantly increased PI3K/mTOR levels in the primary tumor cells (p<0.001 by Mann-Whitney test), whereas the levels in CTCs did not change.

Single cell analysis showed that the PI3K/mTOR index in primary tumor cells and CTCs was significantly decreased following BEZ235 treatment (**Fig. 3d**; p<0.0001). Suppression of pS6 and p4EBP1 levels in the primary tumor following BEZ235 treatment was further confirmed through quantification of immunohistochemical staining of these tissues (**Fig. 3e-3g**; p4EBP1: p<0.001; pS6: p=0.0011). Interestingly, despite the PI3K/mTOR index in the lung and liver metastases being comparable to the primary tumor (**Fig. 2c**), these metastatic tumor cells, unlike the primary tumor, did not show any significant differences in PI3K/mTOR activity upon drug treatment (**Fig. 3d**).

To determine whether drug induced signaling changes in the PI3K/mTOR pathway coincide with growth responses, we evaluated PC3 tumor-bearing mice treated with the PI3K inhibitor GDC0941, which did not affect tumor growth in this model (**Fig. 3b**). Single cells collected from PC3 tumor bearing mice untreated or treated with GDC0941 for 8 weeks showed that the PI3K/mTOR signaling axis was not significantly suppressed in the primary tumor or CTCs (**Fig. 3h**) suggesting that suppression of the PI3K/mTOR is indeed associated with growth responses to these drugs. Taken together, these results indicate that that PI3K/mTOR signaling activity and drug responsiveness of different tumor compartments and distal sites may be modulated by the tumor microenvironment.

Since epithelial to mesenchymal transition (EMT) is a critical process implicated in promoting resistance to therapeutic intervention with kinase inhibitors, we analyzed the extent to which EMT impacts BEZ235 responsiveness in the PC3 mouse tumor model. We first stained the untreated and BEZ235-treated primary tumors with antibodies against EpCAM and GFP, which distinguishes the human tumor cells from mouse stromal cells. EpCAM expression was heterogeneous throughout the GFP^+^ tumor tissue (**Fig. 4a**). The percentage of EpCAM positive cells was drastically reduced in the drug-resistant primary tumors remaining after 11 weeks of treatment (**Fig. 4b**; p<0.001; **Fig. 4c**; Primary tumors: p=0.037; CTC: p=0.0014). Islands of low-EpCAM (EpCAM^low^) expressing tumor cells in untreated primary tumors suggest the preexistence of these resistant cells in untreated tumors. The presence of resistant tumor cells prior to drug exposure has been shown in several models^36–38^. To determine whether high and low EpCAM, an epithelial marker, accurately identified epithelial and mesenchymal states, respectively, we interrogated the expression of the mesenchymal marker, N-Cadherin in EpCAM^high^ and EpCAM^low^ PC3 cells; EpCAM^high^ cells expressed lower N-Cadherin compared with EpCAM^low^ cells demonstrating that EpCAM levels accurately differentiate epithelial and mesenchymal PC3 populations (**Extended Data Fig. 7**).

**Figure 4:**
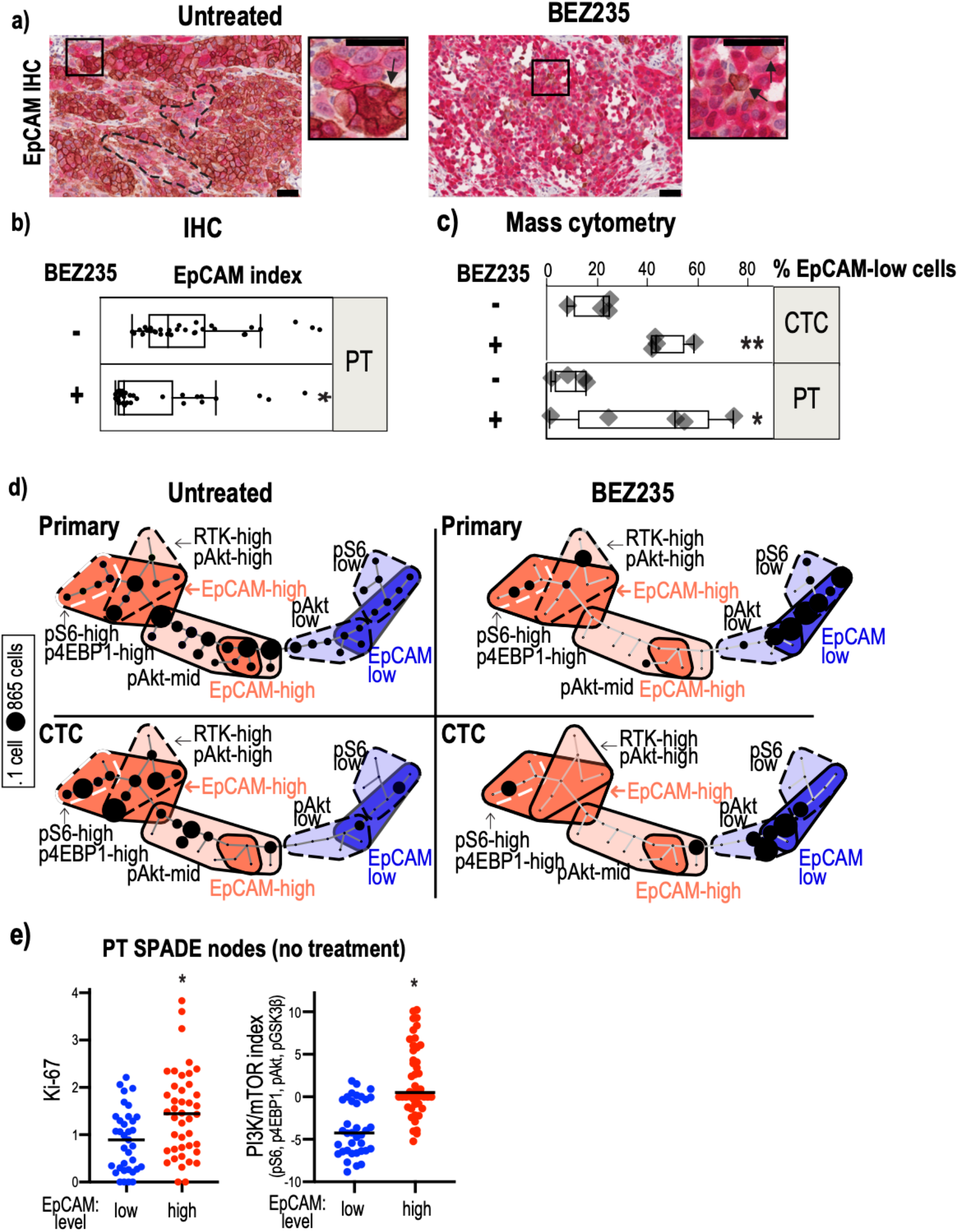
Epithelial and mesenchymal tumor cell populations exhibit differential PI3K/mTOR activity. 4a: EpCAM (brown-DAB) and GFP (red-alkaline phosphatase) staining of primary tumors resected from untreated and BEZ235-treated mice. Sections were counterstained with hematoxylin. Clusters of EpCAM^low^ cells (marked with hatched lines) pre-exist within EpCAM^high^ cell populations in the untreated tumors. A EpCAM^high^ cell cluster is marked with an arrow in the high magnification image of the inset. BEZ235-treated tumors have a prevalence of EpCAM^low^ cells with very few EpCAM positive cells (marked by arrows in the higher magnification inset). Scale bar represents 50µm. 4b: Quantification of EpCAM intensity in the untreated and drug-treated primary tumors following immunohistochemical staining with an EpCAM antibody. p<0.001 by Mann-Whitney test. 4c: The bar graph shows the percentage of EpCAM^low^ cells in untreated and BEZ235-treated primary tumor (PT) cells (p=0.037 by t-test) and in CTCs (p=0.0014 by t-test). EpCAM^low^ is defined by the median level of EpCAM in all primary tumor cells. 4d: SPADE map of primary tumor cells and CTCs from untreated and BEZ235-treated mice shows two major tumor cell populations defined by EpCAM expression: EpCAM^high^ (red) and EpCAM^low^ (blue). The EpCAM^high^ cells are further classified into three subclasses based on the expression levels: EpCAM^high^-RTK^high^-pAkt^high^, EpCAM^hghi^-pS6^high^-p4EBP1^high^, EpCAM^hi^-pAkt^mid^) and mesenchymal subpopulations into four subclasses (EpCAM^very-low^-pS6^low^, EpCAM^very-low^-pAkt^low^, EpCAM^low^pAkt^mid^, EpCAM^low^pS6^low^). The number of tumor cells within each compartment is illustrated using circles, the size of which corresponds to the number of cells. 4e: Median Ki67 and PI3K/mTOR levels were compared in SPADE nodes of untreated primary tumors that had low or high median EpCAM levels. EpCAM^high^ nodes harbored significantly higher Ki67 (p=0.0003 by Mann-Whitney test) and PI3K/mTOR index (p<0.0001 by Mann-Whitney test).

Quantification of EpCAM by single cell proteomic analysis of untreated primary tumor cells identified the co-existence of EpCAM^low^ and EpCAM^high^ populations with differential kinase activities (**Fig. 4d**). The EpCAM^high^ epithelial populations had elevated PI3K/mTOR activity marked by increased pS6, p4EBP1 and pAkt, as well as RTKs whereas the EpCAM^low^ expressing mesenchymal cells harbored lower kinase activities.

Quantification of Ki67 showed that the EpCAM^high^ cells, are more proliferative compared with the EpCAM^low^ cells, which harbor a lower PI3K/mTOR index (**Fig. 4e**; Ki67: p=0.0003; PI3K/mTOR: p<0.0001). The CTCs isolated from untreated mice also show the coexistence of PI3K/mTOR activity high and low epithelial and mesenchymal populations, respectively (**Fig. 4d**). Consistent with their elevated PI3K/mTOR activity, the epithelial populations are more sensitive to BEZ235 and their decline upon treatment coincided with an increase in mesenchymal cells with low pAkt and pS6 activity (**Fig. 4d**).

To determine if the differential PI3K/mTOR kinase activation in epithelial and mesenchymal tumor cell populations observed in the mouse models could be extrapolated to human patient-derived tumor cells, we interrogated single cell RNA-sequencing data from CTCs isolated from mCRPC (n=77 CTCs from 13 patients, described above^35^) and breast cancer patients (n=241 CTCs from a total 78 patients across two different datasets^39^). Expression of the EMT gene signature^40^ identified 42 epithelial and 35 mesenchymal CTCs in mCRPC patients (**Fig. 5a, 5b** (left); p=0.0098). The gene expression signature associated with PI3K/mTOR activity^41, 42^ was significantly elevated in epithelial CTCs compared with mesenchymal CTCs in these prostate cancer patients (**Fig. 5b** (middle); p=1.1e^-07^); CTCs in 5 patients who exhibited radiographic and/or PSA progression during enzalutamide therapy harbored higher PI3K/mTOR activity compared with 8 patients who had not received enzalutamide (**Fig. 5b** (right); p=0.041, **Extended Data Fig. 8**). We then extended this study to breast cancer patients: single cell RNA-sequencing analysis of breast CTCs showed that the gene expression signature associated with PI3K/mTOR activity^41, 42^ was significantly elevated in epithelial CTCs compared with mesenchymal CTCs in both dataset 1 (EMT: p=5.8e^-34^; PI3K/mTOR: p=6.2e^-19^) and in dataset 2 (EMT: p=1.5e^-14^; PI3K/mTOR: p=1.3e^-14^) (**Fig. 5c, Extended Data Fig. 9, 10**). Since clinical outcome data was available for breast cancer patient dataset 2 (GSE144494, **Supplementary Table 2**), we investigated whether elevated PI3K/mTOR activity in CTCs correlated with either overall survival (OS) and/or time to progression (TTP) on previous therapy. Using a Cox proportional hazards model, we identified that increased PI3K/mTOR activity in epithelial CTCs is associated with poor OS [Log Rank p=0.0023; hazard ratio (HR)=4.4 (1.6, 12)] and TTP [Log Rank p=0.012; HR=2.7 (1.2, 5.9)] (**Fig. 5d**, **Extended Data Fig. 11**).

**Figure 5:**
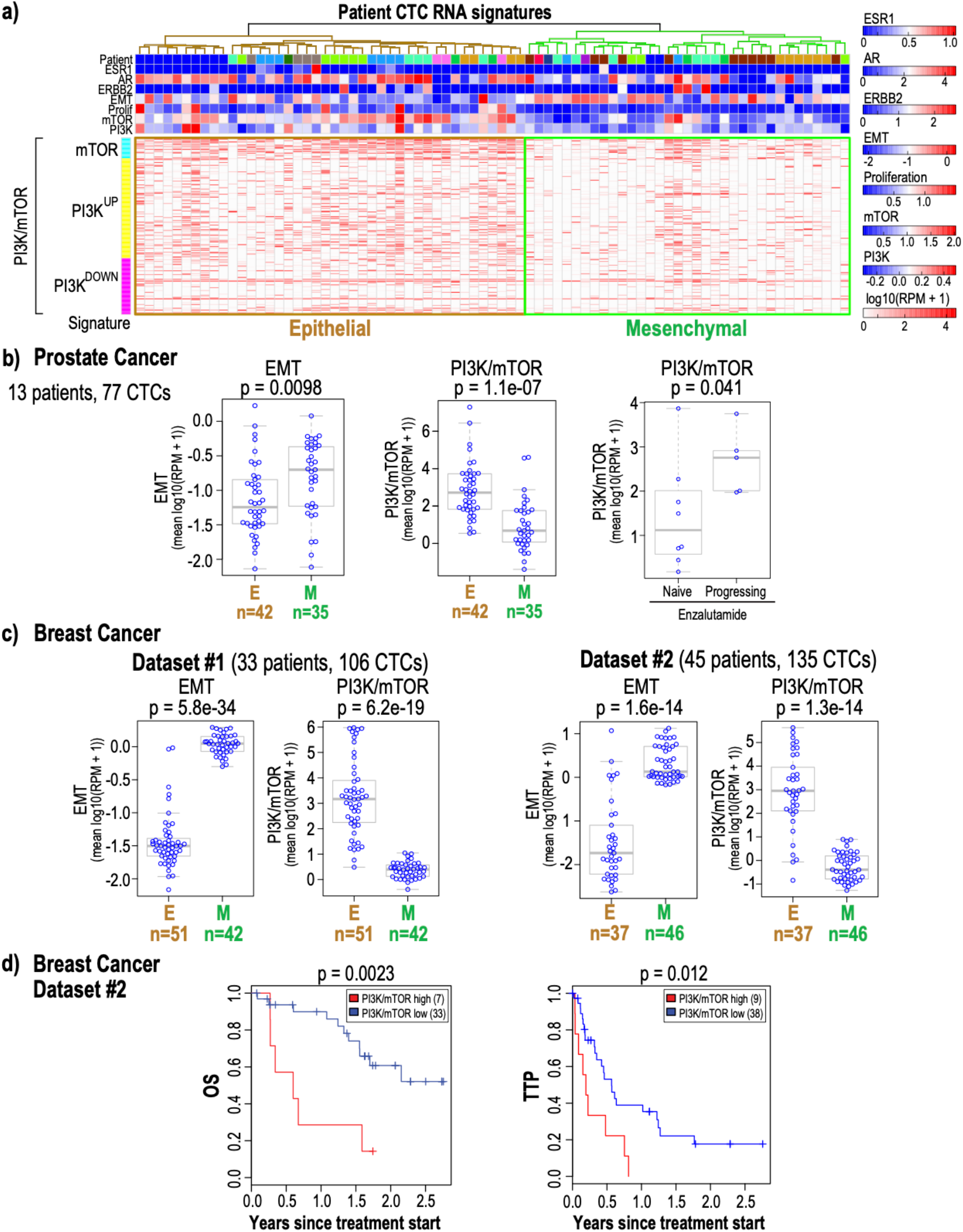
Epithelial and mesenchymal CTC populations enriched from metastatic breast and mCRPC patients exhibit differential PI3K/mTOR activity. 5a: Single cell RNA-sequencing data derived from 77 CTCs enriched from 13 mCRPC patients (mean 6 CTCs per patient) was interrogated for the expression of estrogen and androgen receptor and ERBB2 expression as well as for proliferation^54^, EMT^40^, PI3K and mTOR^41, 42^ signatures. Expression values represent log10 (RPM+1). The epithelial and mesenchymal CTC populations identified based on EMT marker expression are highlighted. 5b: The plots show the level of EMT (left) and combined PI3K/mTOR (middle) gene signatures in each individual epithelial and mesenchymal CTCs enriched from mCRPC patients. The number of CTCs in each category is shown below. The right panel shows that patients with increased PI3K/mTOR signaling progress on enzalutamide compared with treatment naïve patients who harbor lower levels of PI3K/mTOR activity. The p-values shown were calculated using the two-sided Welch t-test. 5c: The plots show the level of the EMT (left) and combined PI3K/mTOR (middle) gene signatures in each individual epithelial and mesenchymal CTCs enriched from two sets of breast cancer patients (dataset #1 and dataset#2). The number of CTCs in each category is shown below. The p-values shown were calculated using the two-sided Welch t-test. 5d: Kaplan-Meier analysis of overall survival (OS) and time to progression (TTP) on previous therapy for metastatic breast cancer patients (dataset #2) with high average PI3K/mTOR activity versus low average PI3K/mTOR activity. The high and low PI3K/mTOR subgroups were determined based on average PI3K/mTOR activity for each patient blood draw using Otsu’s method^55^. P-value was calculated by log rank test.

In summary, single cell proteomic analysis of PTEN-deleted prostate cancer cells residing within different tissue compartments in mouse prostate cancer xenograft models identified tumor-microenvironment and cell-state specific activation of kinase networks generating inter- and intra-tumoral heterogeneity that modulate growth and drug responses. The increased PI3K/mTOR activity in metastatic prostate tumors growing in the bone microenvironment, compared with isogenic tumor cells residing within other tissue compartments, suggests that it is independent of the mutational status of the tumor; instead, tumor microenvironmental factors, which remain to be identified, may be significantly contributing to this differential kinase activity. Although the 29-antibody panel identifies cell state-specific activation of the PI3K–Akt–mTOR kinase axis specifically in PTEN-deleted prostate cancer xenograft models, the analysis of CTCs from breast cancer patient data set 2 (for which PI3K mutational data was available) shows that increased of PI3K/mTOR activity in CTCs is independent of the patient’s PI3K mutational status (p=0.0021; **Extended Data Fig. 12**). This suggests that the differential kinase activity identified in the epithelial and mesenchymal tumor compartments may be dictated by the lineage differences of tumor cells.

The high and low kinase activity in the epithelial and mesenchymal tumor cell populations, respectively, provides a mechanism underlying the differential sensitivity of these cell states to kinase inhibitors and identifies a new molecular paradigm for the kinase inhibitor-resistant phenotypes associated with EMT, as observed by the accumulation of PI3K/mTOR^low^ mesenchymal cells in PC3 tumors treated with BEZ235 (**Fig. 4a-d**). The PI3K/mTOR signaling pathway is a master regulator of the RNA translation machinery. Inhibiting this pathway suppresses the translation of mRNAs encoding ribosomal proteins^43^. We recently showed that a subset of proliferative CTCs from breast cancer patients harbor strong ribosome and translational gene signatures, which coincided with the expression of epithelial lineage markers^39^. It is possible that increased PI3K/mTOR activity in the epithelial CTCs contributes to increased translation in this subset of CTCs and it may sensitize them to inhibitors of translation as well.

PI3K/mTOR activation, typically by genetic aberrations in PTEN and PI3K, is common in advanced castration resistant prostate cancer^21^ as well as in other cancers^19^ including breast cancer^18, 20^. Deregulation of the PI3K-Akt-mTOR signaling axis, an important intracellular pathway in regulating cell cycle, quiescence, and proliferation presents an attractive molecularly targetable therapeutic node. The first PI3K inhibitor, alpelisib, was recently approved for clinical management of breast cancer patients^44^; alpelisib plus fulvestrant led to a complete or partial response in heavily pretreated metastatic breast cancer patients with PI3K-mutated tumors^45^. Although the first-generation mTOR inhibitors have limited efficacy due to partial inhibitory effects, the second-generation mTOR inhibitors, currently under clinical investigation^46^, might be more competent in blocking mTOR activity through complete inhibition of 4EBP1 phosphorylation^47–49^. A third-generation of mTOR inhibitors being generated by connecting rapamycin-like domain to an ATP-site may further enhance their activity in the clinic^50^. Complementing current therapeutic interventions in cancers including prostate and breast cancer with inhibitors targeting PI3K/mTOR may prove useful to clinically exploit the PI3K/mTOR-driven molecular heterogeneity exhibited by the different tumor cell populations.

Recently, simultaneously suppressing androgen-signaling with abiraterone and the PI3K-Akt-mTOR-signaling axis with the Akt inhibitor, ipatasertib, (compared with abiraterone alone) prolonged radiographic progression free survival in mCRPC patients with PTEN-loss^51^. As such, combining inhibitors of both PI3K/mTOR and androgen receptor activities may result in superior anti-tumor activity compared with either agent alone. Our data shows that CTCs enriched from mCRPC patients progressing on enzalutamide indeed harbor increased PI3K/mTOR activity (**Fig. 5b**; right panel) which may render them sensitive to inhibitors of PI3K/mTOR signaling axis.

Finally, the striking difference in c-MET activity in bone metastases compared to other metastatic sites may provide an intriguing explanation for the puzzling results from the pivotal phase 3 clinical trial of the c-MET/VEGFR2 inhibitor cabozantanib, which demonstrated profound bone scan responses in metastatic castration-resistant prostate cancer patients and yet failed to improve overall survival^52, 53^. Based on our data, one potential mechanistic explanation for this result is that the drug preferentially targets bone metastases which have increased c-MET activity, but does not induce sufficient tumor cell killing in other sites of disease where c-MET activity may be lower due to differences in the tumor microenvironment. Such heterogeneity in kinase activity across multiple metastatic sites may necessitate the use of combination therapies to ensure simultaneous targeting of all therapeutic vulnerabilities of cancer cells in different tissue compartments.

## Acknowledgements

We thank Dr. Nick Dyson for critical reading of the manuscript. We thank Michael Koulopoulos, Dr. Narges Rashidi, MGH Pathology Core Facility, MGH Flow and Mass Cytometry Core Facility for experimental assistance. This work was supported by funding from the ESSCO Breast Cancer Research Fund (S.M), BCRF Drug Research Collaborative (S.M.) NIH/NCI (U01CA214297; S.M., D. A. H., M.T.) NIH P41 BioMEMS Resource Center (EB002503; M.T.), NIH/NIBIB (EB012493; M.T.), the Howard Hughes Medical Institute (D.A.H.), NIH/NCI (2R01CA129933; D.A.H.), National Foundation for Cancer Research (D.A.H.), Shriners Hospital for Children Mass Spectrometry Special Shared Facility (N.M.K.), Harvard Medical School Eleanor and Miles Shore Fellowship (N.M.K) and Tosteson & Fund for Medical Discovery Fellowship (N.M.K.).

## Author Contributions

N.M.K, Y.Z., T.D.D. and S.M. conceived and designed the study. N.M.K. designed and performed mass cytometry experiments and analyzed the data. N.M.K. and D.W. performed microfluidic isolation of CTCs. N.M.K, Y.Z., T.D.D., V.C., U.H. performed cell culture experiments. Y.Z., T.D.D., R.B., V.C., K.N. performed mouse experiments and imaging. T.D.D. and E.E. performed western blotting. D.S.M., B.W. performed analysis of patient RNA sequencing and survival data. M.T. developed CTC-iChip Technology. D.T.T. provided RNA sequencing support. S.L.S. provided blood preservation method. L.N. assisted with microscopy. N.M.K., Y.Z., T.D.D., D.H. and S. M. wrote the manuscript. All authors discussed results and provided input and edits on the manuscript.

## Extended Data Figures

**Extended Data Figure 1:**
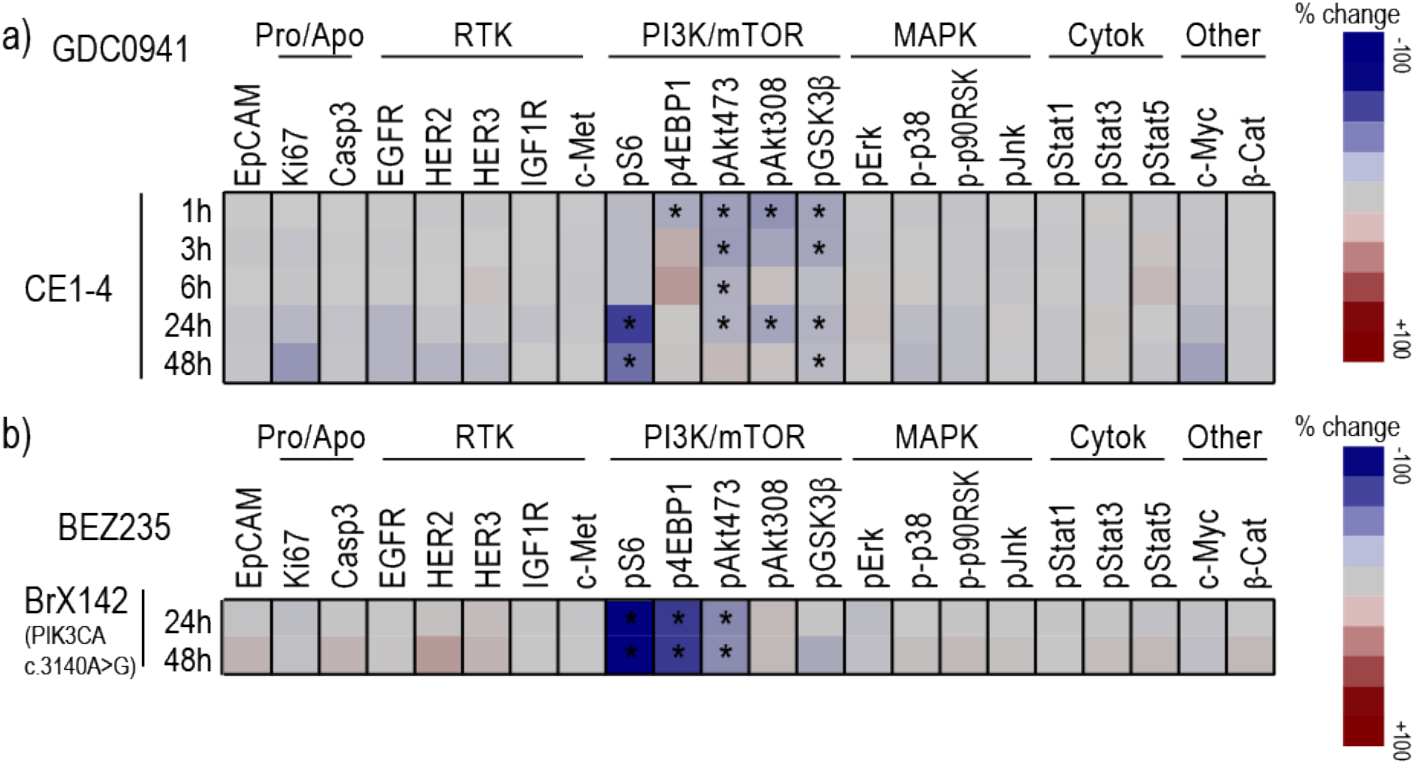
1a: Heatmap showing mean change in the signaling proteins within each of the indicated pathways in CE1-4 cells treated *in vitro* with the PI3K inhibitor, GDC0941, compared with untreated control cells (n=2 experimental replicates). Significant changes are marked with asterisks. p<0.01 corrected for multiple comparisons by Bonferroni-Dunn. 1b: Heatmap showing mean changes in proteins following BEZ235 treatment of the breast cancer patient derived PI3K mutant CTC cell line, BRX142. p<0.01 corrected for multiple comparisons by Bonferroni-Dunn.

**Extended Data Figure 2:**
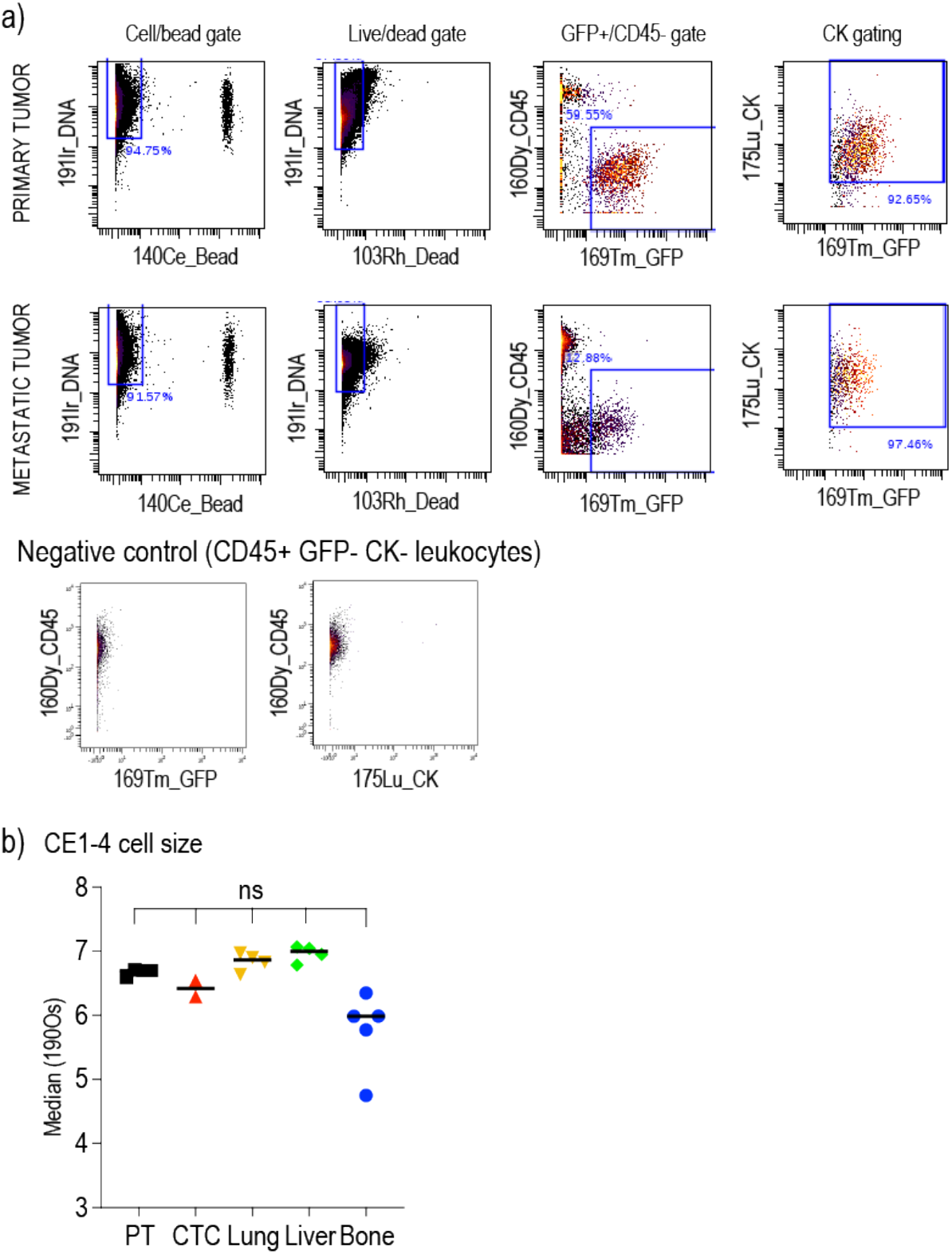
2a: Gating strategy used for selecting tumor cells from mass cytometry data of mouse prostate cancer xenograft models. We performed manual gating of events positive for nucleic acid intercalator, CK, GFP and negative for CD45, Ce140 (control bead). Top and middle series of panels show cells from primary and metastatic PC3 tumors, bottom panel shows leukocytes. 2b: Osmium tetroxide labeling applied to the CE1-4 tumor model to compare cell sizes across the different tumor sites shows no significant differences (PT: Primary tumor, ns: not significant) (p=0.2301 for PT vs Bone, p=0.2441 for PT vs Liver, p=0.4637 for PT vs Lung, p=0.4637 for PT vs CTC).

**Extended Data Figure 3:**
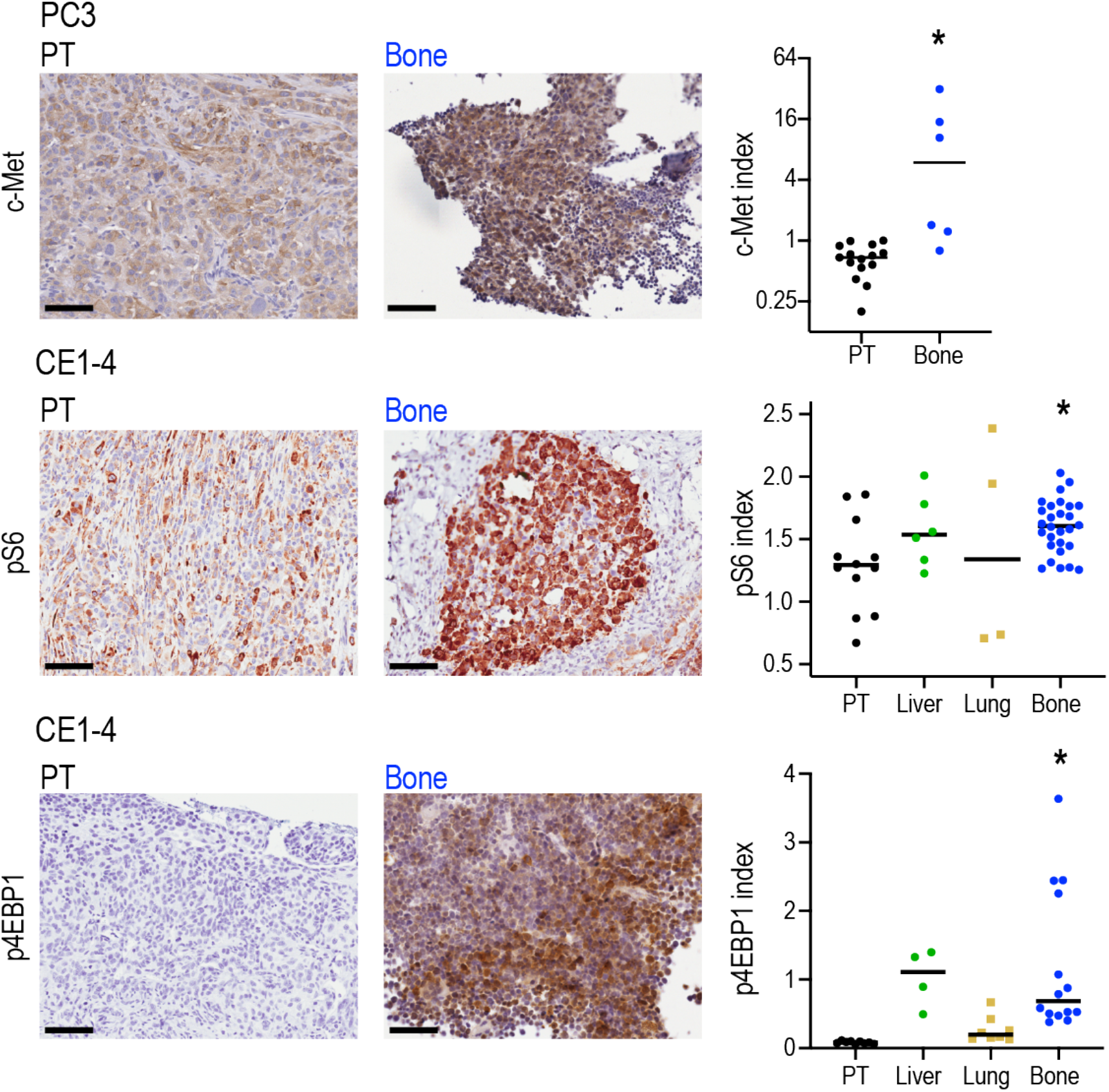
The primary tumors and bone metastases derived from the xenograft models shown were stained for c-Met, pS6 or p4EBP1 (DAB) and counterstained with hematoxylin. Representative images are shown in left. Scale bars represent 50 µm. Quantification of c-Met, pS6 and p4EBP1 levels show significantly increased expression for all three epitopes in bone metastases when compared to primary tumors (right panels). c-Met: p=0.0182 by Mann-Whitney test, pS6: p=0.0386, and p4EBP1 p=0.0019 by one-way ANOVA, Holm-Sidak’s multiple comparisons test.

**Extended Data Figure 4:**
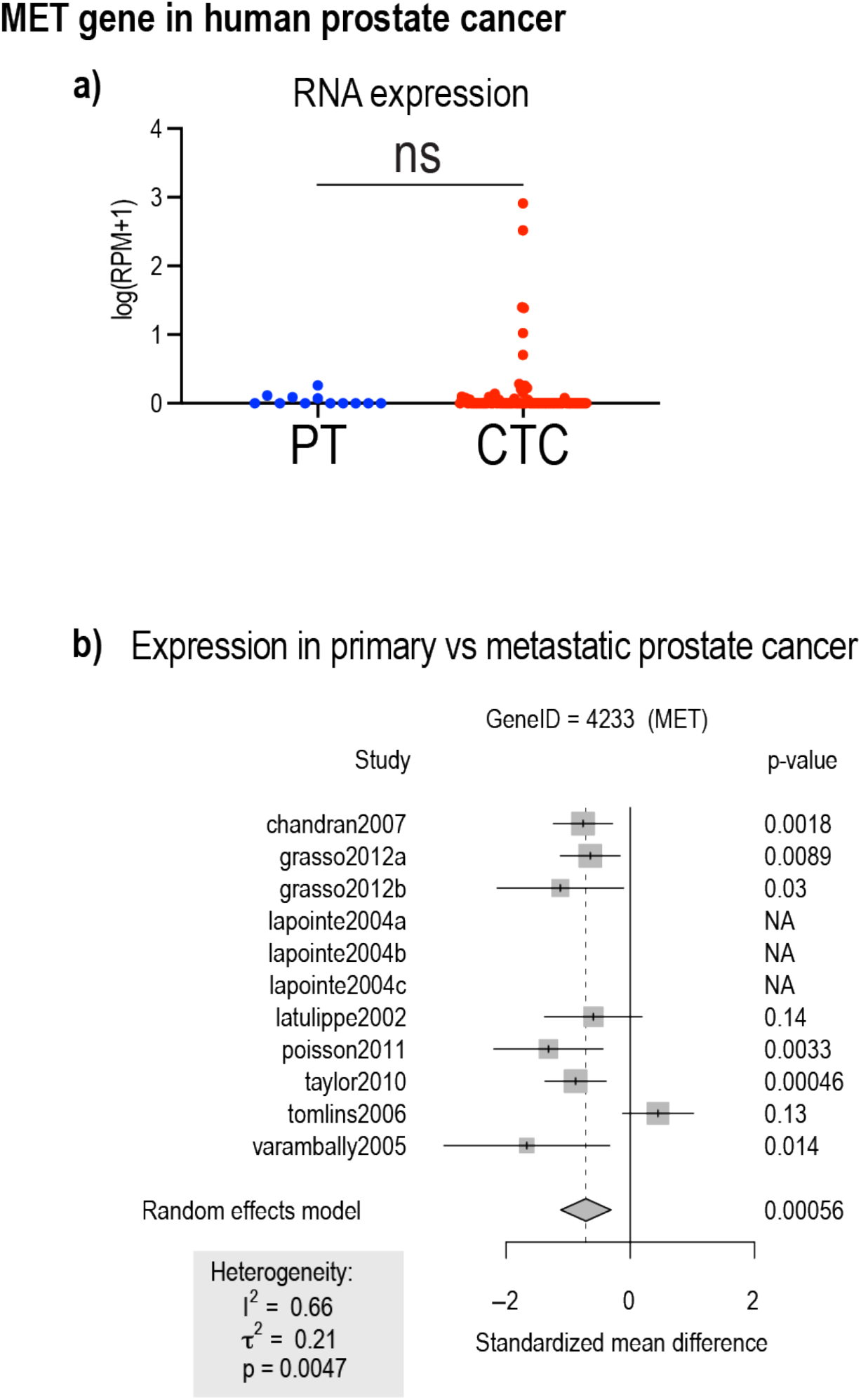
4a: c-Met transcript is not significantly increased in single cell RNA-Sequencing data derived form 77 CTCs collected from 13 metastatic castration resistant prostate cancer patients, compared with 12 primary prostate cancers sequenced at single cell resolution (PT)^35^. 4b: Meta-analysis of c-Met gene expression in prostate cancer datasets that have both primary and metastatic tumors. Data shows significantly reduced c-Met mRNA expression in metastatic tumors.

**Extended Data Figure 5:**
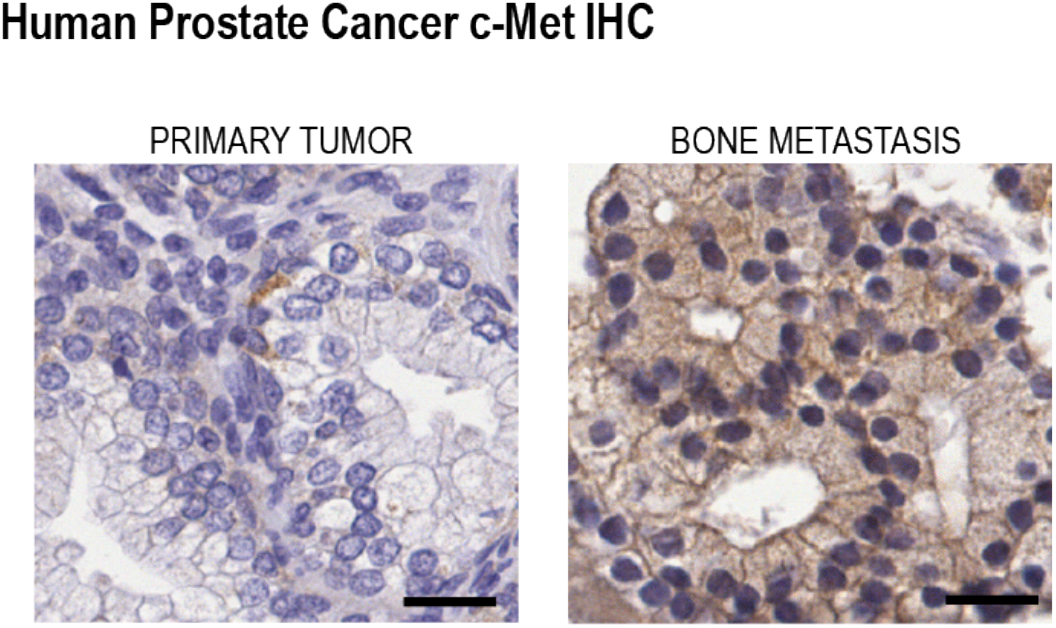
c-Met protein expression in the primary tumor and in bone metastasis derived from human prostate cancer patients was measured using IHC. Representative images of c-Met and hematoxylin counter-stained primary tumors and bone metastases are shown. Scale bar represents 50µm. The quantification of c-Met index showed elevated c-Met protein levels in bone metastases compared with primary tumors (**Fig. 2h**).

**Extended Data Figure 6:**
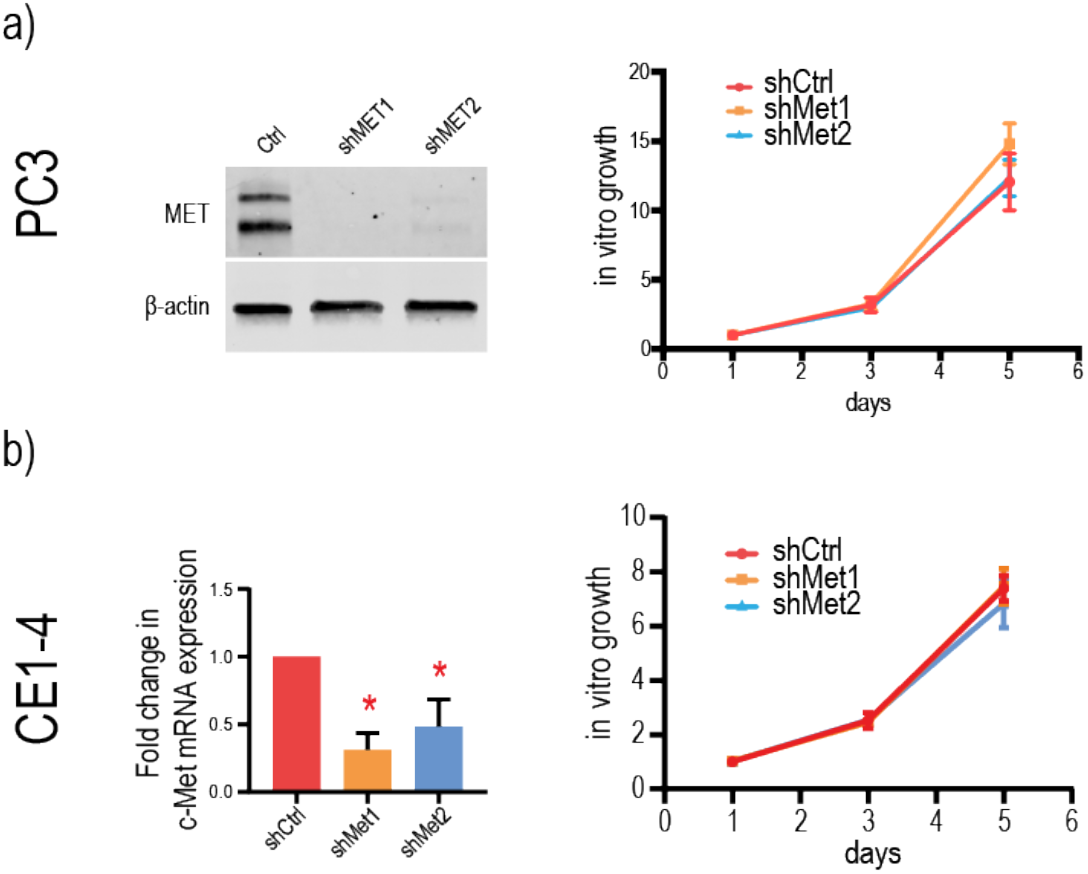
6a: Knockdown of c-Met expression in PC3 cells using two different shRNAs against c-Met. Depletion of c-Met protein in PC3 cells is shown. ß-actin is shown as loading control. The plot on the right shows that c-Met knockdown does not impair cell proliferation *in vitro*. q=0.8654 CTR vs shMET1, q=0.9553 CTR vs shMET2 by two-way ANOVA, FDR correction. 6b: Knockdown of c-Met expression in CE1-4 cells using two different shRNAs against c-Met shows reduced expression c-Met RNA (p<0.01, t-test). c-Met-KD does not interfere with *in vitro* proliferation. q=0.8999 CTR vs shMET1, q=0.5529 CTR vs shMET2, two-way ANOVA, FDR correction.

**Extended Data Figure 7:**
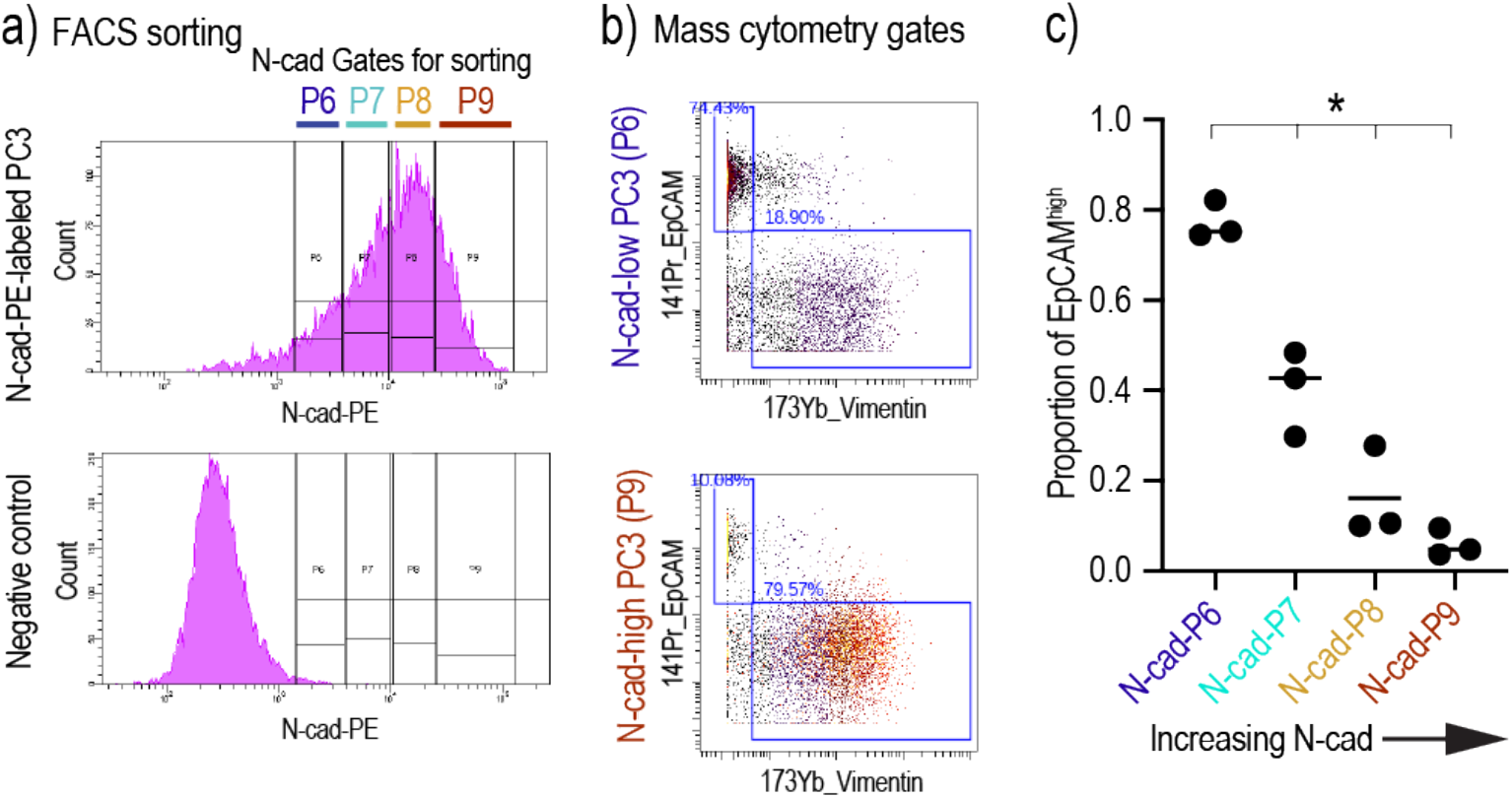
7a: PC3 cells, grown *in vitro*, were sorted into 4 populations based on their level of N-Cadherin (N-Cad) expression, labeled as P6, P7, P8 and P9, from lowest to highest N-Cad, respectively. Top panel shows the level of N-Cad detected in sorted cells via PE fluorescence intensity. Bottom panel; trypsin treated N-Cad cells that lost the N-Cad epitope are shown as negative control. 7b: Gating strategy for analysis of EpCAM^high^ cell proportion in N-Cad sorted PC3 cells, top panel showing P6 (N-Cad^low^) and bottom panel showing P9 (N-Cad^high^). Top left quadrant was used to quantify EpCAM^high^ cells. 7c: Proportion of EpCAM^high^ cells compared within sorted populations. Proportion of EpCAM^high^ PC3 population significantly decreases with increasing N-Cad levels. q=0.0207 N-cad-P6 vs N-cad-P7, q=0.0207 N-cad-P6 vs N-cad-P8, q=0.0087 for N-cad-P6 vs N-cad-P9, one-way ANOVA, FDR correction.

**Extended Data Figure 8:**
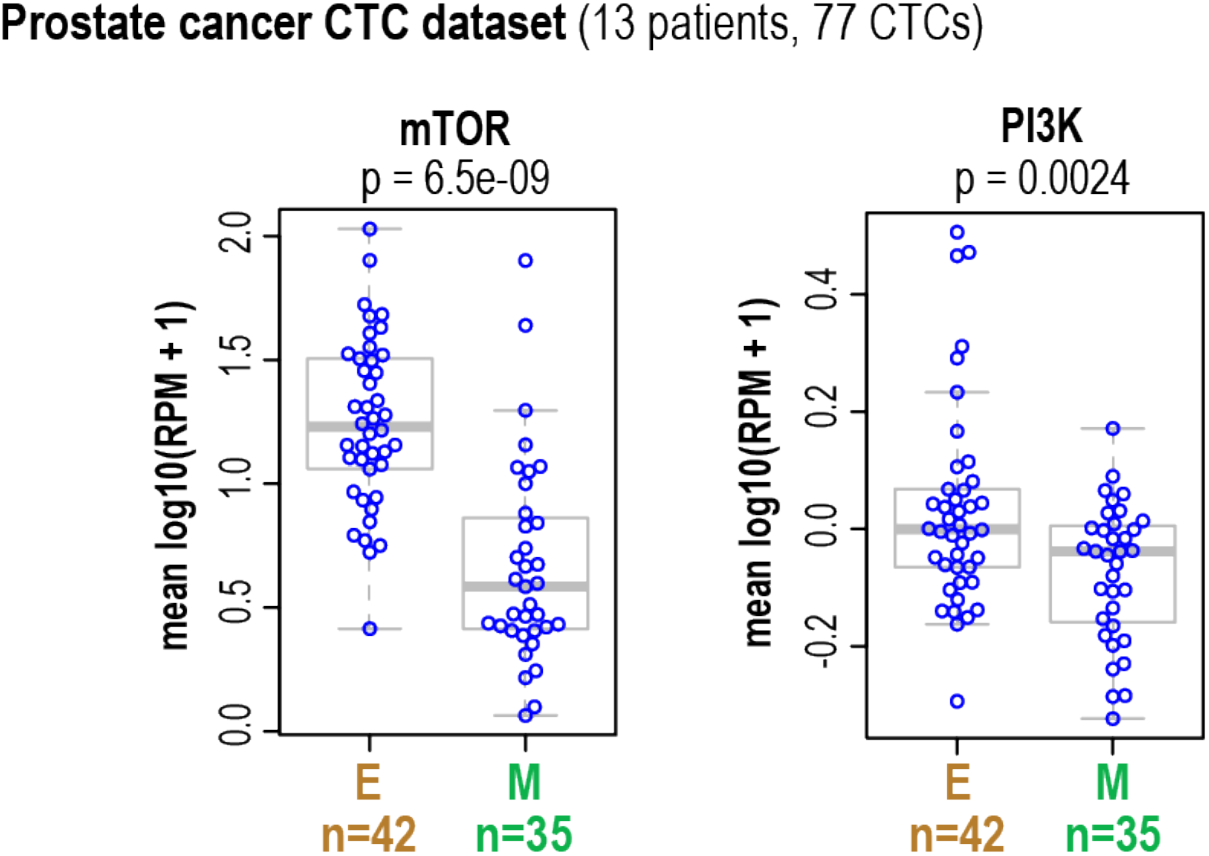
Epithelial and mesenchymal prostate cancer CTC subsets were defined based on single cell RNA sequencing data and analyzed for mTOR and PI3K signatures. The p-values shown were calculated using the two-sided Welch t-test. Dendrogram that represents the entire dataset is reported in **Fig. 5a**.

**Extended Data Figure 9:**
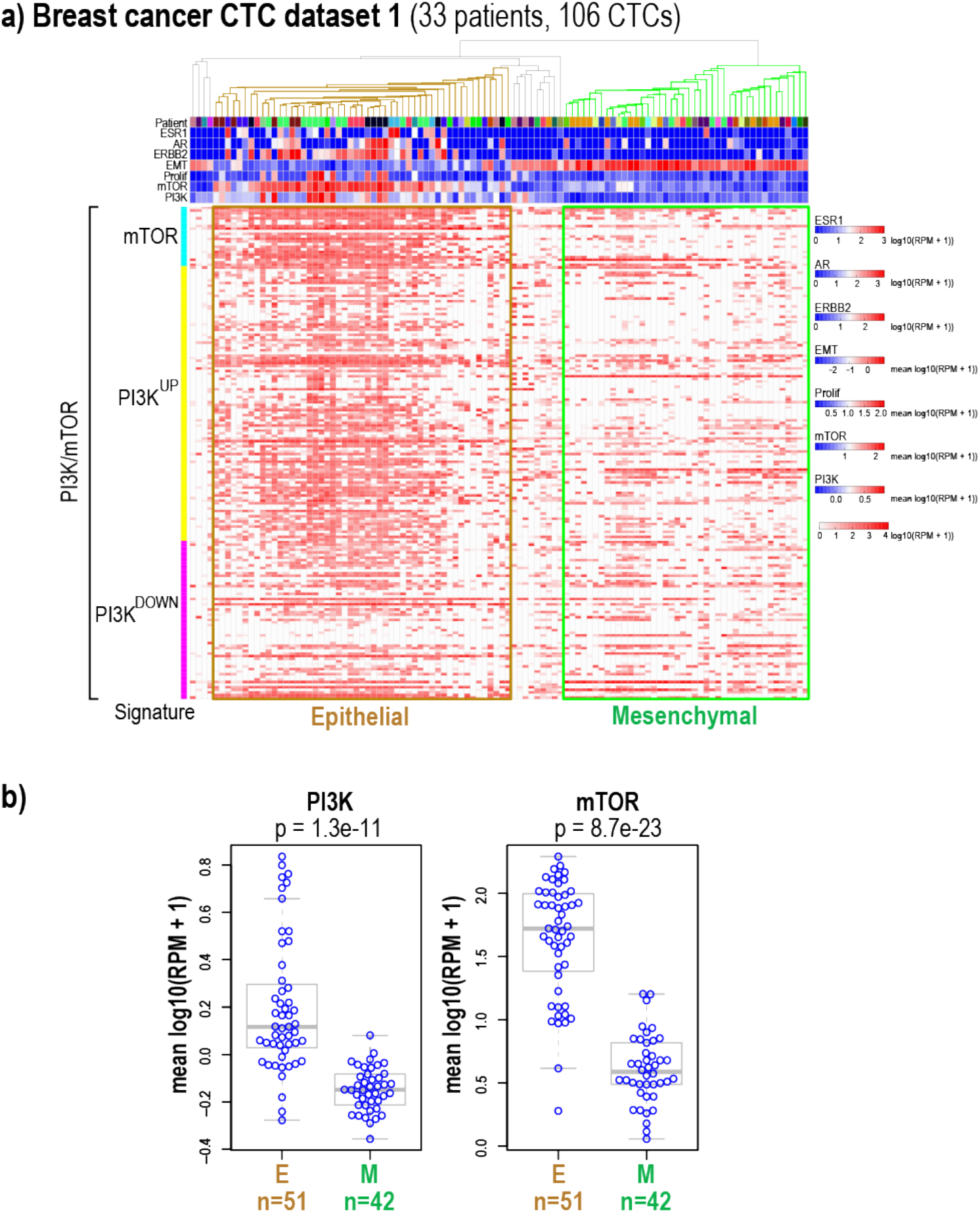
9a: Single cell RNA-Sequencing of 106 CTCs enriched from 33 metastatic breast cancer patients (Breast cancer dataset #1). Top dendrogram represents EMT signature-based hierarchical clustering of single CTCs represented in columns below. Top rows represent patient ID, and gene signature levels. Rows below represent individual transcript levels used for calculating mTOR and PI3K signatures. 9b: Based on the hierarchical clustering, epithelial and mesenchymal subsets of CTCs were selected and analyzed for PI3K and mTOR signatures. The p-values shown were calculated using the two-sided Welch t-test.

**Extended Data Figure 10:**
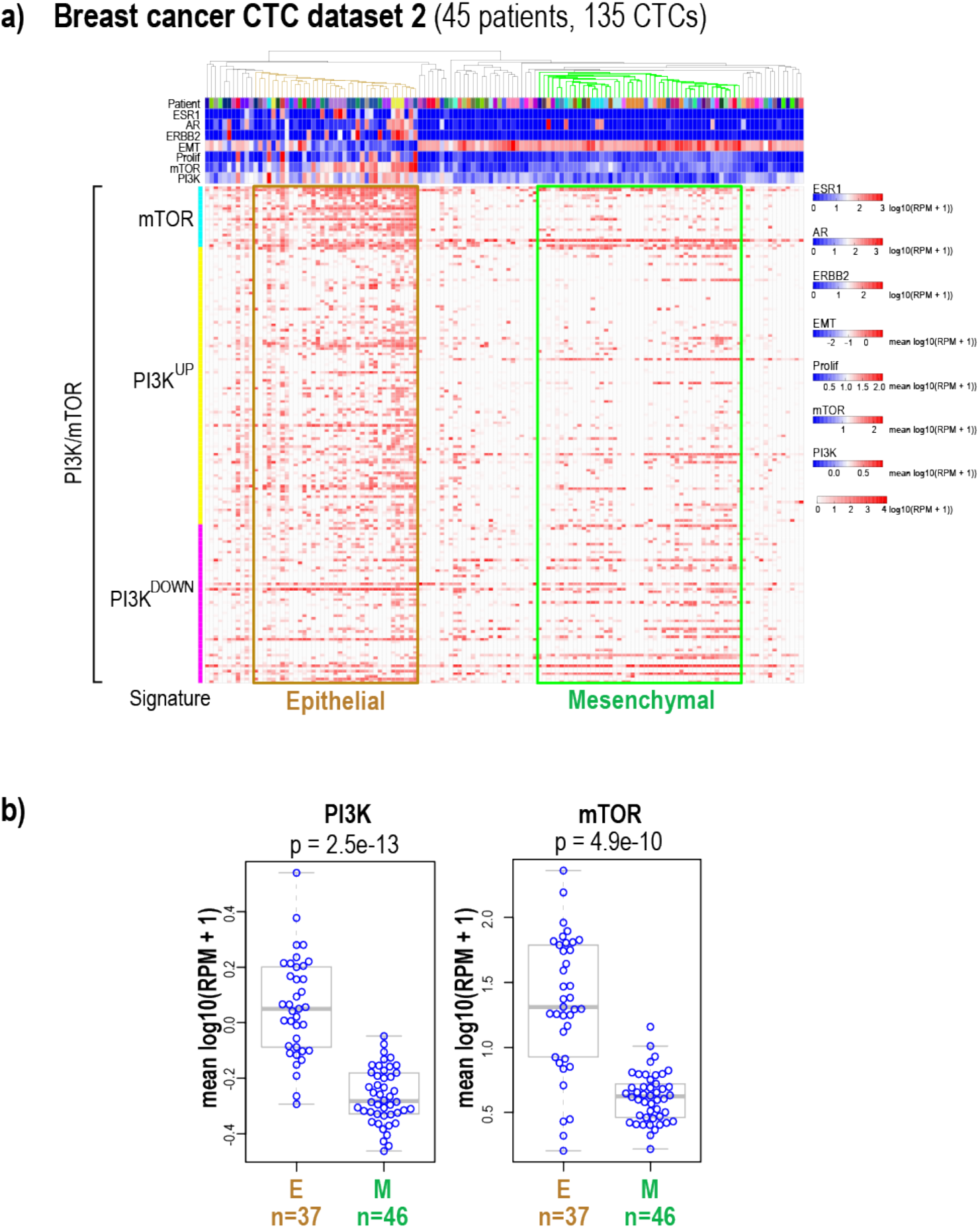
10a: Single cell RNA-Sequencing of 135 CTCs enriched from 45 metastatic breast cancer patients (Breast cancer dataset #2). Top dendrogram represents EMT signature-based hierarchical clustering of single CTCs represented in columns below. Top rows represent patient ID, and gene signature levels. Rows below represent individual transcript levels used for calculating mTOR and PI3K signatures. 10b: Based on the hierarchical clustering, epithelial and mesenchymal subsets of CTCs were selected and analyzed for PI3K and mTOR signatures. The p-values shown were calculated using the two-sided Welch t-test.

**Extended Data Figure 11:**
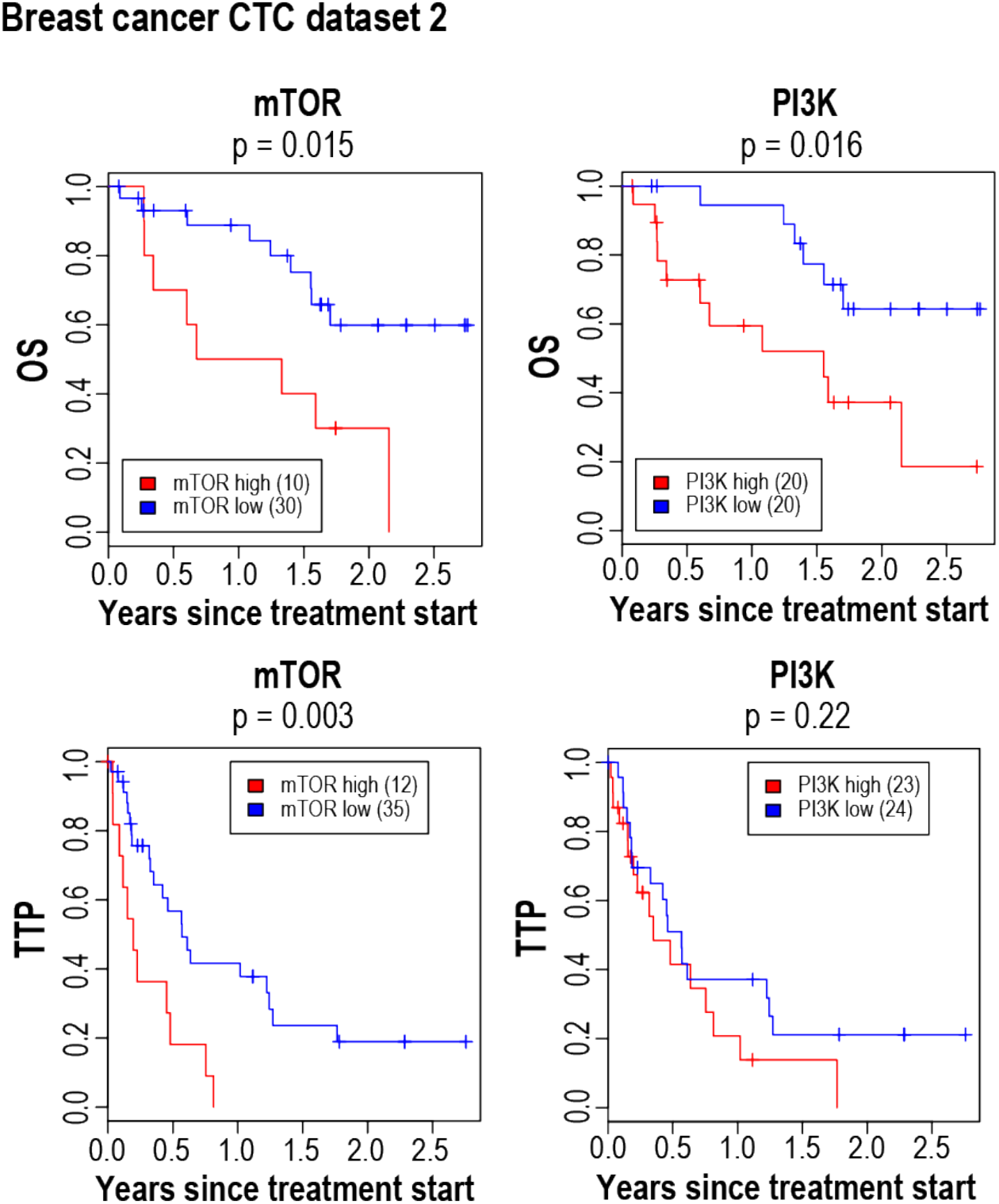
Kaplan-Meier analysis of overall survival (OS) and time to progression (TTP) on previous therapy for breast cancer patients (dataset #2, Supplementary Table 2) with high average PI3K and mTOR activity versus low average PI3K and mTOR activity. The high and low PI3K and mTOR subgroups were determined based on average PI3K and mTOR activity for each patient blood draw. P-value was calculated by log rank test.

**Extended Data Figure 12:**
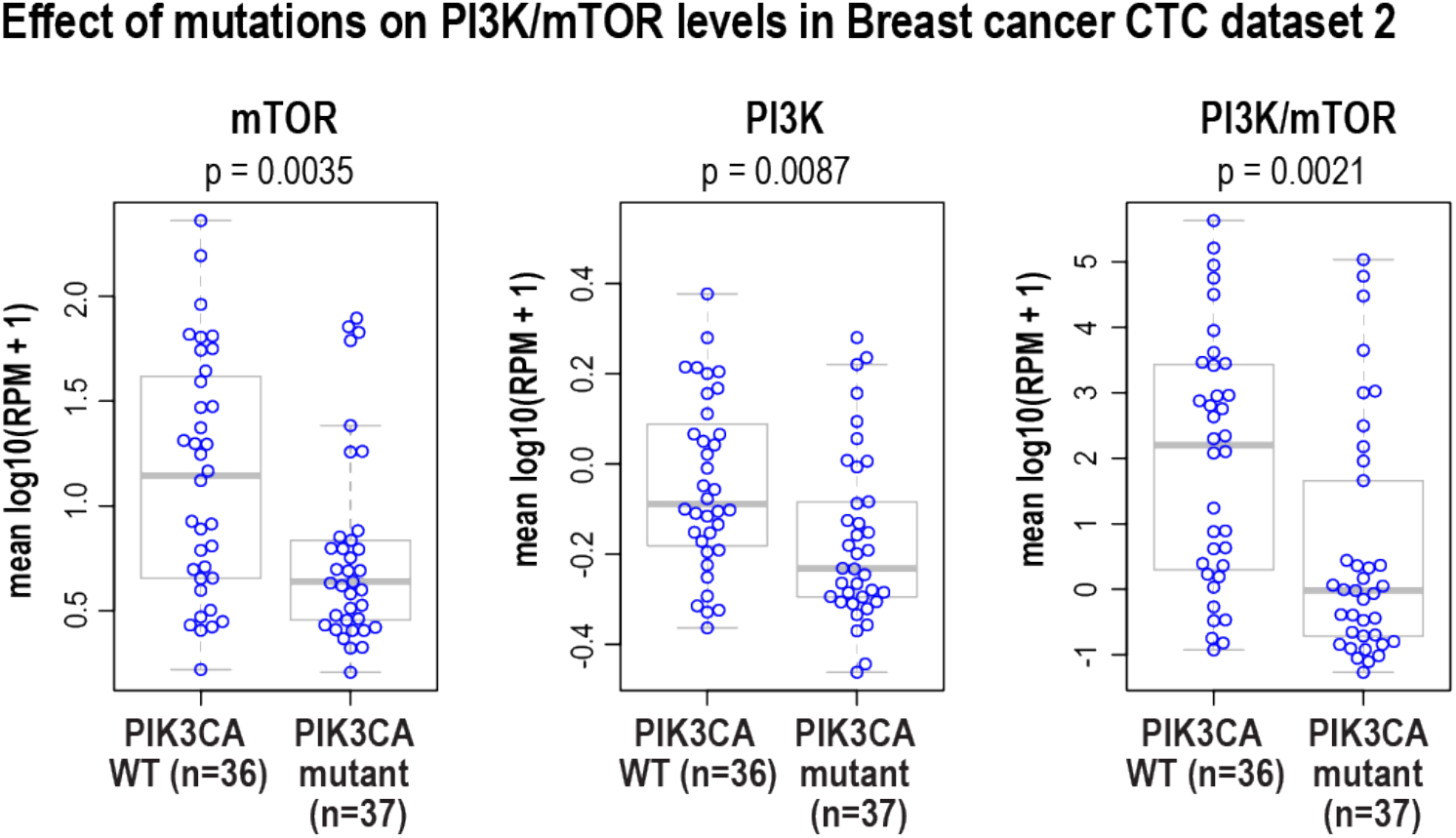
The plots show the level of the individual PI3K and mTOR signaling and combined PI3K/mTOR gene signatures in each individual CTCs enriched from breast cancer patients (dataset #2) harboring wild type or mutant PI3K. The p-values shown were calculated using the two-sided Welch t-test.

**Extended Data Figure 13:**
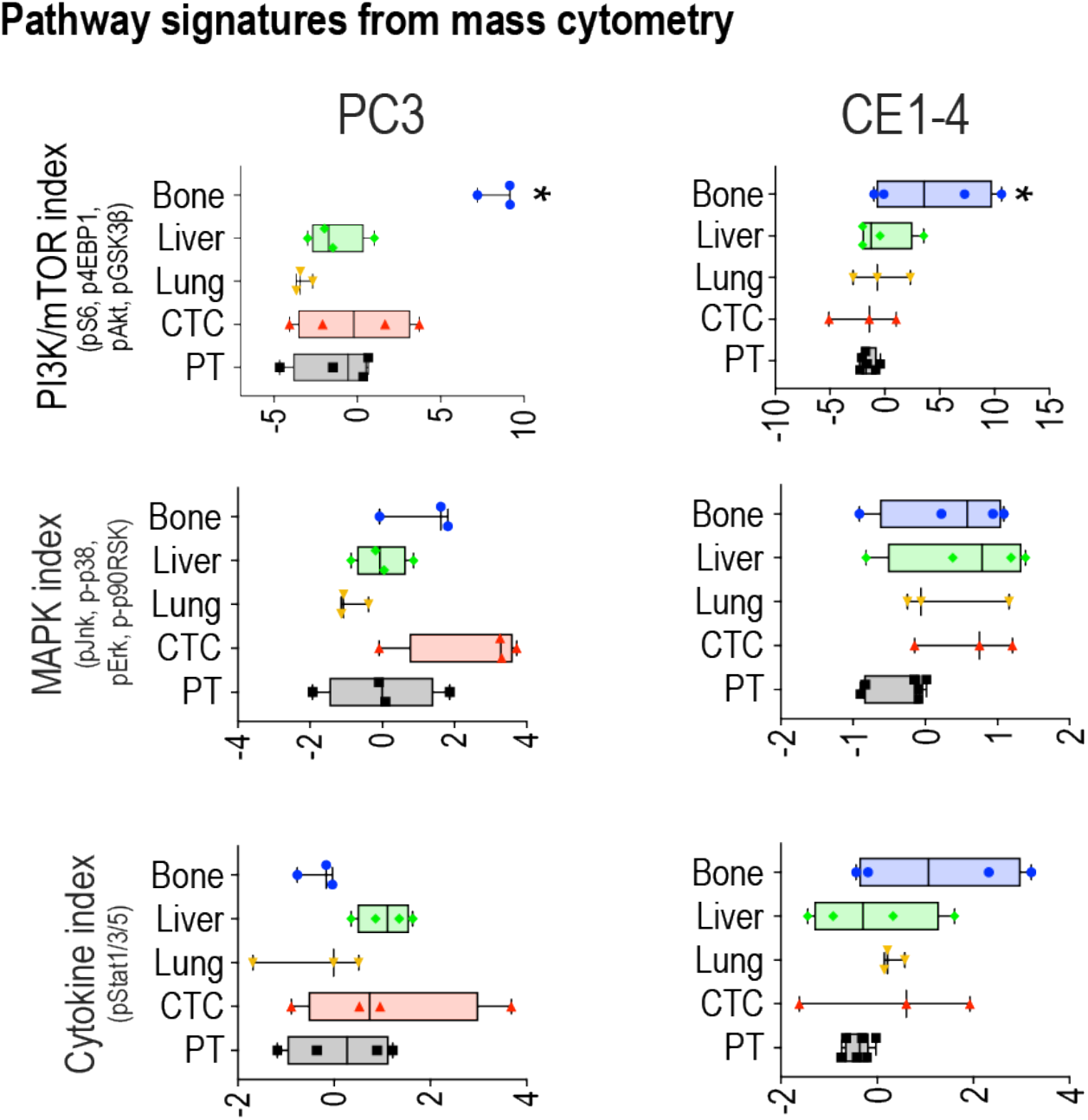
PI3K/mTOR, MAPK and cytokine pathway signatures show median levels of each of the pathways in primary and metastatic tumor cells and in CTCs in PC3 and CE1-4 models. These signatures were calculated by summing standardized levels of epitopes shown in y axes. Pathways significantly elevated are marked with asterisks. q<0.1, one-way ANOVA, FDR correction.

**Extended Data Table 1:**

| <u>Mass</u> | <u>Metal</u> | <u>Epitope</u> | <u>Clone</u> | <u>Concentration</u> |  | <u>Vendor</u> |
| --- | --- | --- | --- | --- | --- | --- |
| 141 | Pr | EpCAM | EBA-1 | 0.5 | µg/ml | BD |
| 142 | Nd | Casp3 (cleaved) | D3E9 | 0.5 | % | Fluidigm |
| 143 | Nd | β-cat (active) | D2U8Y | 0.25 | µg/ml | Cell Signaling |
| 145 | Nd | EGFR | D38B1 | 0.2 | µg/ml | Cell Signaling |
| 146 | Nd | IgG | DA1E | 0.1 | µg/ml | Cell Signaling |
| 147 | Sm | HER3 | D22C5 | 0.25 | µg/ml | Cell Signaling |
| 148 | Nd | HER2 | 29D8 | 0.25 | µg/ml | Cell Signaling |
| 149 | Sm | p4EBP1 (T37/46) | 236B4 | 0.25 | % | Fluidigm |
| 150 | Nd | pStat5 (Y694) | 47 | 1 | % | Fluidigm |
| 151 | Eu | c-Met | D1C2 | 0.2 | µg/ml | Cell Signaling |
| 152 | Sm | pAkt (S473) | D9E | 1 | % | Fluidigm |
| 153 | Eu | pStat1 (Y701) | 58D6 | 0.25 | µg/ml | Cell Signaling |
| 154 | Sm | pAkt (T308) | D25E6 | 0.25 | µg/ml | Cell Signaling |
| 156 | Gd | p-p38 | D3F9 | 0.5 | % | Fluidigm |
| 158 | Gd | pStat3 (Y705) | Y705 | 1 | % | Fluidigm |
| 159 | Tb | c-Myc | D3N8F | 0.25 | µg/ml | Cell Signaling |
| 160 | Dy | m-CD45 | 30-F11 | 0.05 | µg/ml | Biolegend |
| 162 | Dy | pJnk (T183/Y185) | G9 | 0.3 | µg/ml | Cell Signaling |
| 164 | Dy | p-p90RSK (S380) | D5D8 | 0.25 | µg/ml | Cell Signaling |
| 165 | Ho | pGSK3β (S9) | D85E12 | 0.1 | µg/ml | Cell Signaling |
| 166 | Er | CD44 | IM7 | 0.05 | µg/ml | Biolegend |
| 168 | Er | Ki-67 | B56 | 0.5 | % | Fluidigm |
| 169 | Tm | GFP | 5F12.4 | 0.5 | % | Fluidigm |
| 170 | Er | IGF1R-B | D23H3 | 0.3 | µg/ml | Cell Signaling |
| 171 | Yb | pErk (T202/Y204) | D13.14.4E | 0.5 | µg/ml | Cell Signaling |
| 172 | Yb | pS6 (S235/236) | N7-548 | 0.25 | % | Fluidigm |
| 175 | Lu | CK 5, 8, 18 | RCK102, C-04 | 0.025 | µg/ml | Abcam |

## SUPPLEMENTARY INFORMATION

**Supplementary Table 1:** Cell number statistics for PC3 and CE1-4 engrafted mouse models.

**Supplementary Table 2:** Patient survival data for breast cancer dataset 2.

**Supplementary Table 3:** Antibody signal spillover table.

**Supplementary Data:** PC3 ViSNE dot plot colored by antigen level (related to **Fig. 2a**)

## METHODS

### Reagents

Details of antibodies used in mass cytometry experiments are listed in **Extended data table 1**. For experiments involving CE1-4 tumor cells, clones of EpCAM, c-Met and CK antibodies were exchanged with the following clones to detect mouse epitopes: EpCAM: G8.8 (Biolegend), c-Met: 1G7NB (Novus Biologicals), CK: AE1+AE3 (Novus Biologicals). Metal labeling kits were purchased from Fluidigm and used according to manufacturer’s instructions. Mass-tag barcoding reagents were purchased from Fluidigm and used with a modified protocol detailed below. BEZ235 and GDC0941 were purchased from Selleck Chemical, and stock solutions were prepared fresh by dissolving in DMSO at 1 µM concentration. Osmium tetroxide (>99%) was purchased from ACROS Organics. AbC™ Total Antibody Compensation Bead Kit (catalog # A10513) was purchased from Thermo Fisher Scientific and used for validating all the antibodies and generating signal spillover table (**Supplementary Table 3**).

### Cell Culture and lentiviral expression

The human prostate cancer cell line PC3 was obtained from ATCC, authenticated by STR profiling and maintained as recommended. No mycoplasma contamination was detected. The mouse prostate cancer cell line CE1-4 has been described^16^. shRNA constructs against c-Met and non-targeting shRNA control were acquired from the molecular profiling laboratory (MPL) at MGH. Transfection of cells was performed using lipofectamine together with lentiviral packaging plasmids. Lentivirus was collected 48 and 72 hours after transfection. PC3 and CE1-4 cells were infected with lentivirus in the presence of 8μg/ml polybrene and selected in growth medium containing 2 µg/mL puromycin for 3 days.

### Primers and quantitative real-time PCR

Total RNA was extracted using RNeasy Mini Kit (Qiagen). 1µg of RNA was used to generate cDNA using superscript III First Strand synthesis system (Life Technologies). Reactions were amplified and analyzed in triplicate using the ABI 7500 Real-Time PCR System. Primers used to detect c-Met were 5’-TCCCCAATGACCTGCTGAAA-3’ (Forward) and 5’-CTTTTCCAAGGACGGTTGAAGAA-3’ (Reverse).

### Cell Proliferation Assay

Proliferation was measured using CellTiter-Glo (Promega). Briefly, 2,000 cells were plated in each well of a 96-well plate in a volume of 100 μl growth medium at day 0. At indicated time points, 100 μl of CellTiter-Glo reagent was added into each well and measurements were done using a luminescence plate reader.

### Western blot analysis

Cells were lysed in RIPA Buffer (cat# BP-115X) supplemented with Halt^TM^ Protease inhibitor cocktail (Thermo Scientific 78425) and passed through a needle to enhance lysis. Protein concentration was determined using Pierce^TM^ BCA Protein Assay Kit (cat# 23227 Thermo Scientific). Protein lysates, 15 µg per sample, were separated on SDS/4-15% polyacrylamide gels (Bio-Rad) and transferred onto nitrocellulose membranes (Invitrogen). After blocking with 5% BSA buffer for 1 hour at room temperature, membranes were incubated with primary antibodies overnight at 4^0^C and followed with the relevant HRP-conjugated secondary antibodies (Anti-rabbit IgG, HRP-linked Antibody cell signaling-7074 and Anti-mouse IgG, HRP-linked Antibody Cell signaling-7076) and visualized using Clarity Western ECL Substrate (BIORAD) and G box (Syngene). ß-actin was used as a loading control. The following primary antibodies were used for western blotting: c-Met (Cell Signaling; 1:1000), pS6 (Fluidigm; 1:1000), Total S6 (Cell Signaling; 1:1000), p4EBP1 (Fluidigm; 1:500), Total 4EBP1 (Cell Signaling; 1:1000), pAKT308 (Cell Signaling; 1:1000); pAKT473 (Fluidigm; 1:1000), Total Akt (Cell Signaling; 1:1000), pGSK3ß (Cell Signaling; 1:500), Total GSK3ß (Cell Signaling; 1:1000), ß-actin (Cell Signaling; 1:1000).

### PC3 and CE1-4 xenograft models

PTEN-deleted PC3 and CE1-4 orthotopic prostate cancer xenograft models were used to investigate the signaling changes within CTCs as well as in primary and metastatic tumor cells in untreated and drug-treated mice. Briefly, 1×10^6^ cells stably expressing luciferase and GFP were introduced into the prostates of mice. Tumors were grown for two weeks, after which the PC3-tumor-bearing mice were treated with 40mg/kg of BEZ235 for 9 additional weeks with a 5-days on/2-days off weekly regimen. Tumor growth was monitored weekly using the Xenogen IVIS Spectrum *in vivo* imaging system (Caliper Life Sciences). Animal care was in accordance with institutional guidelines.

For evaluating tumor growth in the bone, control or in c-Met depleted PC3 (1,000 cells) and CE1-4 (250 cells) cells with luciferase and GFP expression were inoculated into mouse tibia, and signals were measured by Xenogen IVIS Spectrum *in vivo* imaging system every week for 4 weeks and 16 weeks, respectively.

### Tumor cell Isolation

Blood was directly collected into preservatives containing Cyto-Chex BCT tubes (Streck) and processed the same day using the microfluidic CTC-iChip as described^31^. Briefly, preserved blood was incubated while rocking after addition of biotinylated anti-mouse CD45 for twenty minutes and twenty more minutes after the addition of magnetic microbeads coated with streptavidin. CTC-iChip was used to remove the CD45 positive leukocytes, plasma, platelets and red blood cells. The CTC enriched sample was fixed with 4% PFA, washed with PBS and transferred to 90% methanol and stored at −80^0^C until the labeling step.

PC3 and CE1-4 tumors in the prostate, bone, liver and lung were excised and removed with the help of luminescence and the fluorescence of GFP positive cells. Portions of the tumors were fixed at this stage for immunohistochemical analyses. The rest of the tumors were quickly minced and incubated with collagenase solution (Stemcell Technologies) in serum-free media for thirty minutes in a 37°C incubator. Following digestion, cells were immediately fixed in 4% PFA, washed with PBS, and transferred to 90% methanol and stored at −80^0^C until the labeling step.

### Tumor Cell Labeling

Methanol permeabilized samples were washed with PBS and incubated for thirty minutes with barcoding reagents (5 µl) that were diluted in PBS (500 µl). Cells were washed with 0.5% BSA in PBS and mixed together before labeling was performed. Samples were resuspended in 0.5% BSA in PBS containing the antibody cocktail and incubated at room temperature while being rocked for 1 hour. To minimize the loss of rare CTCs, samples were washed using the CTC-iChip, by processing the cells through the chip without depletion antibodies and magnetic beads. Samples were then incubated overnight with iridium-labeled DNA intercalator with 0.2% PFA in PBS for labeling DNA. Following a wash with 0.5% BSA in FBS, cells were fixed in 4% PFA, washed with pure water and filtered. Internal control beads (Fluidigm Sciences) were added for normalizing any changes to signal response of the instrument. The samples were analyzed using CyTOF 2 housed at the Ragon Institute Facility or CyTOF Helios housed at the MGH Flow and Mass Cytometry Core Facility.

### Mass Cytometry Data Analysis

Data was normalized, concatenated and debarcoded using the software from the Nolan Lab (http://web.stanford.edu/group/nolan/resources.html). Data was gated and analyzed online using Cytobank and all gated events were exported for further analysis at JMP. Tumor cells were identified by gating for Ir intercalator+, ^140^Ce-(internal control bead signal), CD45-, IgG-(non-specific binding), cytokeratin+, GFP+ (**Extended Data Fig. 2**). We used positive control cells (PC3) and negative hematopoietic cells (CD45+) that were labeled with the same cocktail and analyzed the same day to determine the gates. Intensities were normalized using arcsinh transform (arcsinh(x/5)) and exported to JMP software where illustrations were prepared and to Prism and R, where statistics were computed. In samples without barcoding (PC3 BEZ235 *in vivo* treatment), additional data quality control was performed on data, by comparing background antigen levels on CD45+/CK-/GFP- events across all samples and identifying outliers using Mahalanobis distances (JMP). All data supporting the findings of this study are available from the corresponding author on request.

### Patient CTC RNA-Sequence based signaling score calculations

Primary tumors and CTCs isolated from prostate cancer patients and CTCs enriched from breast cancer patients were analyzed by single cell RNA-Seq as previously described^35, 39^. Given a gene signature and reads-per-million (RPM) values for all genes for a sample, we defined the metagene value as the mean over the genes in the signature of log_10_(RPM + 1). If the signature has both up genes and down genes, the log_10_(RPM + 1) values for the down genes are added into the calculation of the mean only after being multiplied by −1. In addition to using the metagene values for a dataset, we sometimes combined the metagene values for two gene signatures as follows. For each gene signature we normalized the metagene values by dividing them by the sample standard deviation of the metagene values across the dataset. We then added the two normalized metagene values for each sample to form the combination metagene value for that sample.

Expression of genes previously identified in a mTOR gene signature (Rad001 sensitive genes)^41^ were averaged. For the PI3-Kinase signature, expression of genes identified in a PI3-kinase signature^42^ were analyzed. Positively correlating genes were assigned a positive log10(RPM+1) value while negatively correlating genes were assigned a negative log10(RPM+1). The average log(RPM+1) expression across primary tumors and circulating tumor cells was compared using the Mann-Whitney comparison with significance consider to be p<0.05.

Creation of Kaplan-Meier plot and calculation of logrank p-value was done on a per-patient basis for breast CTC dataset 1 and on a per-blood-draw basis for breast CTC dataset 2 as follows. If there were multiple CTCs per patient or per blood-draw, the metagenes for all the CTCs from a given patient or blood-draw were averaged. We classified those averages into high and low values using Otsu’s method^55^

### Immunohistochemistry

Discarded formalin fixed paraffin embedded primary prostate tumors and prostate cancer bone metastases from 6 and 5 patients, respectively, were obtained in accordance with Institutional Review Board approved protocols. Tissues were sectioned, and processed for c-Met immunohistochemistry. Portions of the mouse primary tumor samples were fixed overnight in 4% PFA and stored in 70% ethanol at 4°C. Portions of mouse bone samples were fixed and decalcified (Cal-Ex II, Fisher Scientific). Tissues sections were stained with hematoxylin and the following antibodies: anti-c-Met (DAB), anti-pS6 (DAB), anti-p4EBP1 (DAB) or anti-EpCAM (DAB) plus anti-GFP (alkaline phosphatase) and imaged with the Aperio Scanscope. Quantification was done using ImageJ color deconvolution followed by thresholding, and measuring mean number of positive pixels.

## Supplementary Methods

### FACS sorting

BD FACSAria II Cell Sorter was used to sort PC3 cells labeled with the N-Cadherin antibody (Clone 8C11, PE labeled, Biolegend). PE Mouse IgG1 (Biolegend) was used as isotype control. Gating strategy is shown in **Extended Data Fig. 7a**.

### Clustering technique

SPADE clustering was implemented by Cytobank. We performed SPADE with 40 nodes and 1% downsampling, and clustered using the following epitope levels: EpCAM, EGFR, c-Met, HER3, HER2, CD44, pS6, p4EBP1, pAkt308, p-p38, pStat3, pErk, p-p90RSK.

### Statistical analysis

Median levels of individual epitopes from *in vitro* treatment analysis were compared using multiple t-tests, corrected using Bonferroni-Dunn method (p<0.05, number of tests: 200). Median levels of epitopes in xenograft experiments were compared using multiple t-tests with FDR correction (q<0.1) corrected for multiple comparisons using two-stage Benjamini, Krieger and Yekutieli correction. Medians levels of epitopes in BEZ235 or GDC0941 *in vivo* treatment experiments were compared using multiple t-tests with FDR correction corrected for multiple comparisons using two-stage Benjamini, Krieger and Yekutieli correction (q<0.1). For statistics of single cell PI3K/mTOR indices, samples with more than 5,500 cells were down sampled to avoid cells from a single animal to be overrepresented in the result. The down sampling was performed by select random sample function of JMP. The PI3K/mTOR indices were compared by Mann-Whitney test (p<0.001) and was accepted significant only when medians values of PI3K/mTOR of all cells from each animal showed identical directional change.

