## Supplementary Data 1 for "Single cell proteomics of tumor compartments identifies differential kinase activities defining sensitivity to mTOR-PI3-kinase inhibition"

### EpCAM

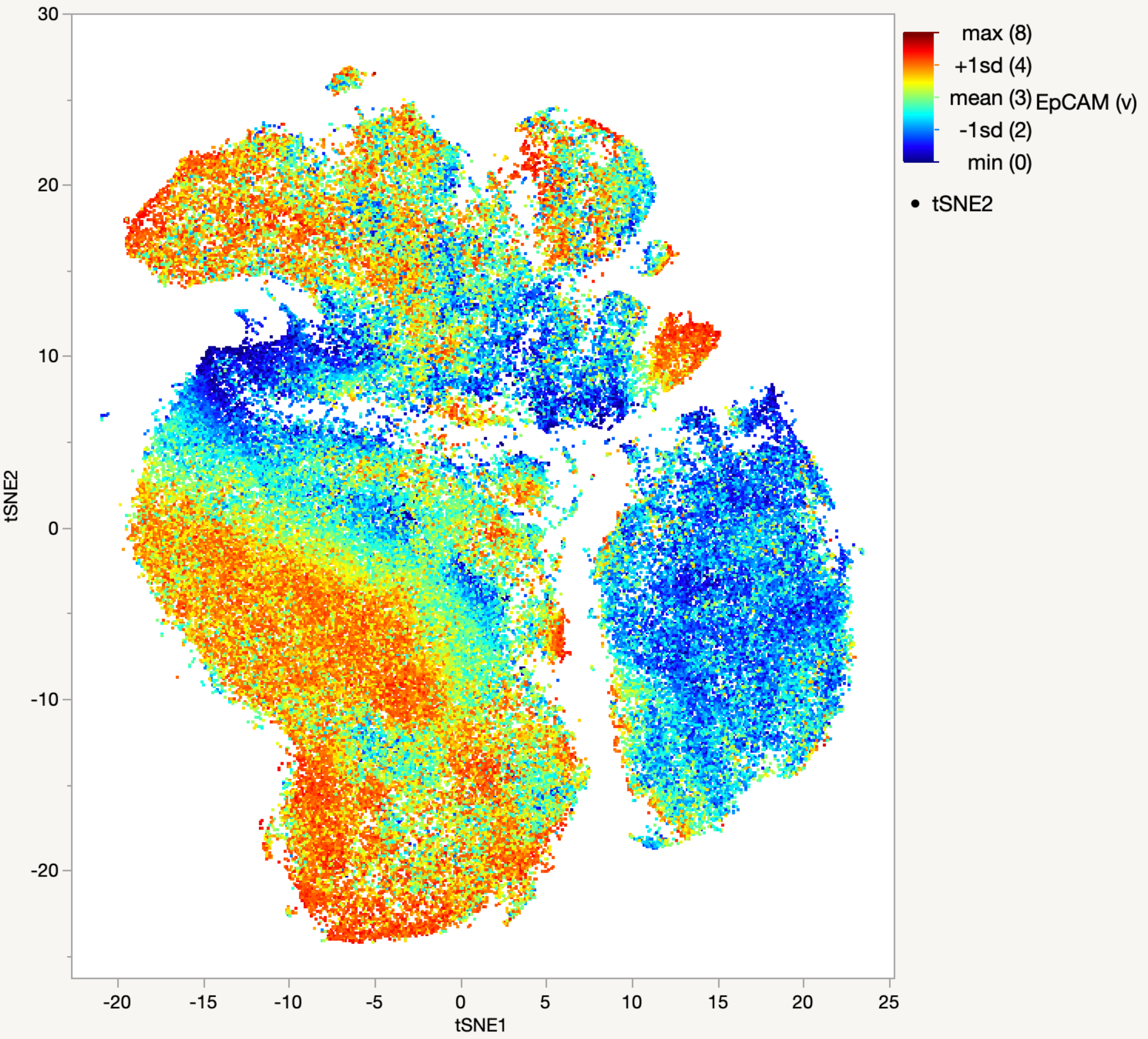

max (8)
+1sd (4)
mean (3)
-1sd (2)
min (0)
EpCAM (v)
tSNE2
tSNE1
30
20
10
0
-10
-20
-20
-15
-10
-5
0
5
10
15
20
25

### c-Met

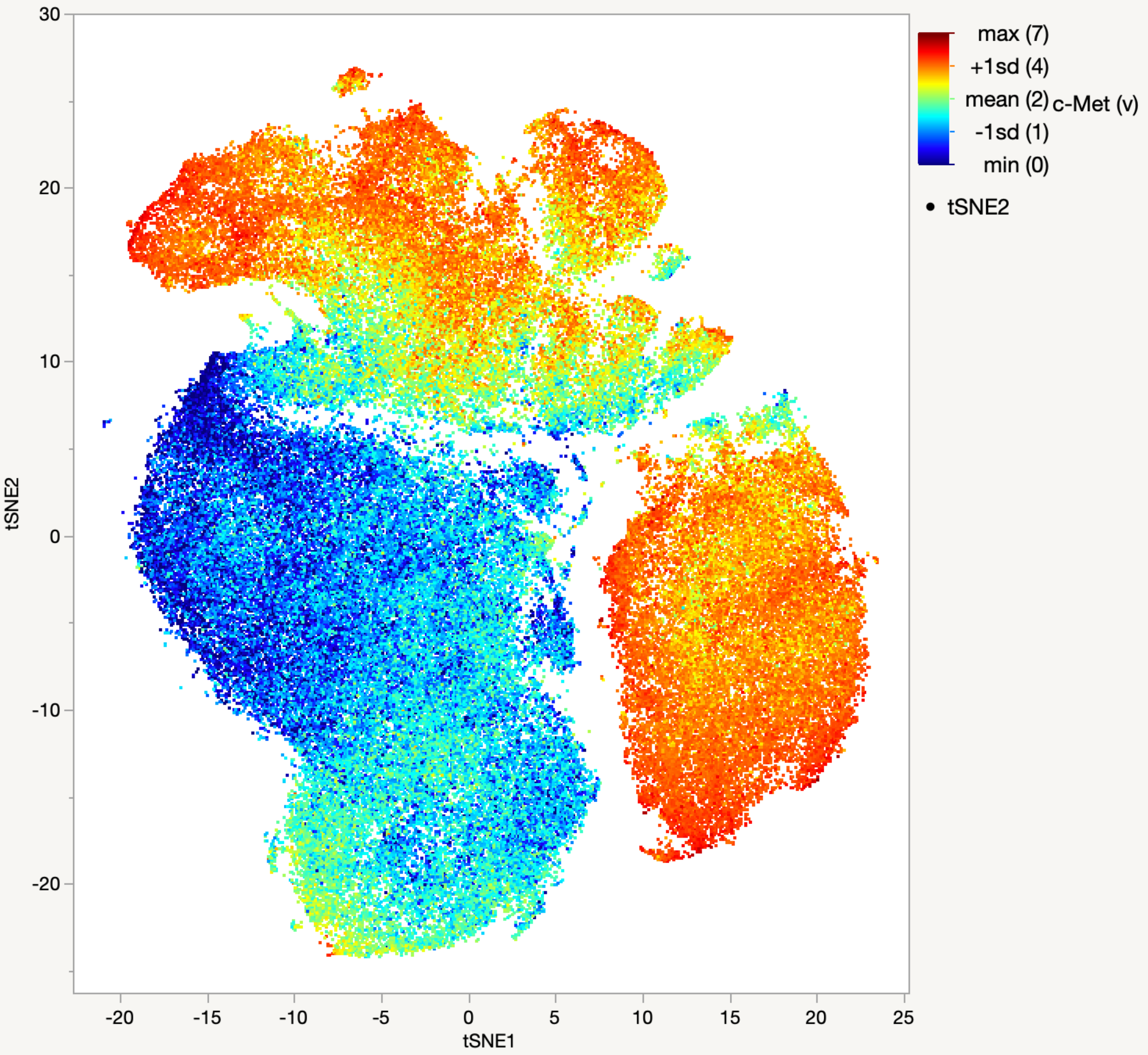

30
20
10
0
-10
-20
tSNE2
-20
-15
-10
-5
0
5
10
15
20
25
tSNE1
max (7)
+1sd (4)
mean (2)
-1sd (1)
min (0)
c-Met (v)
tSNE2

### EGFR

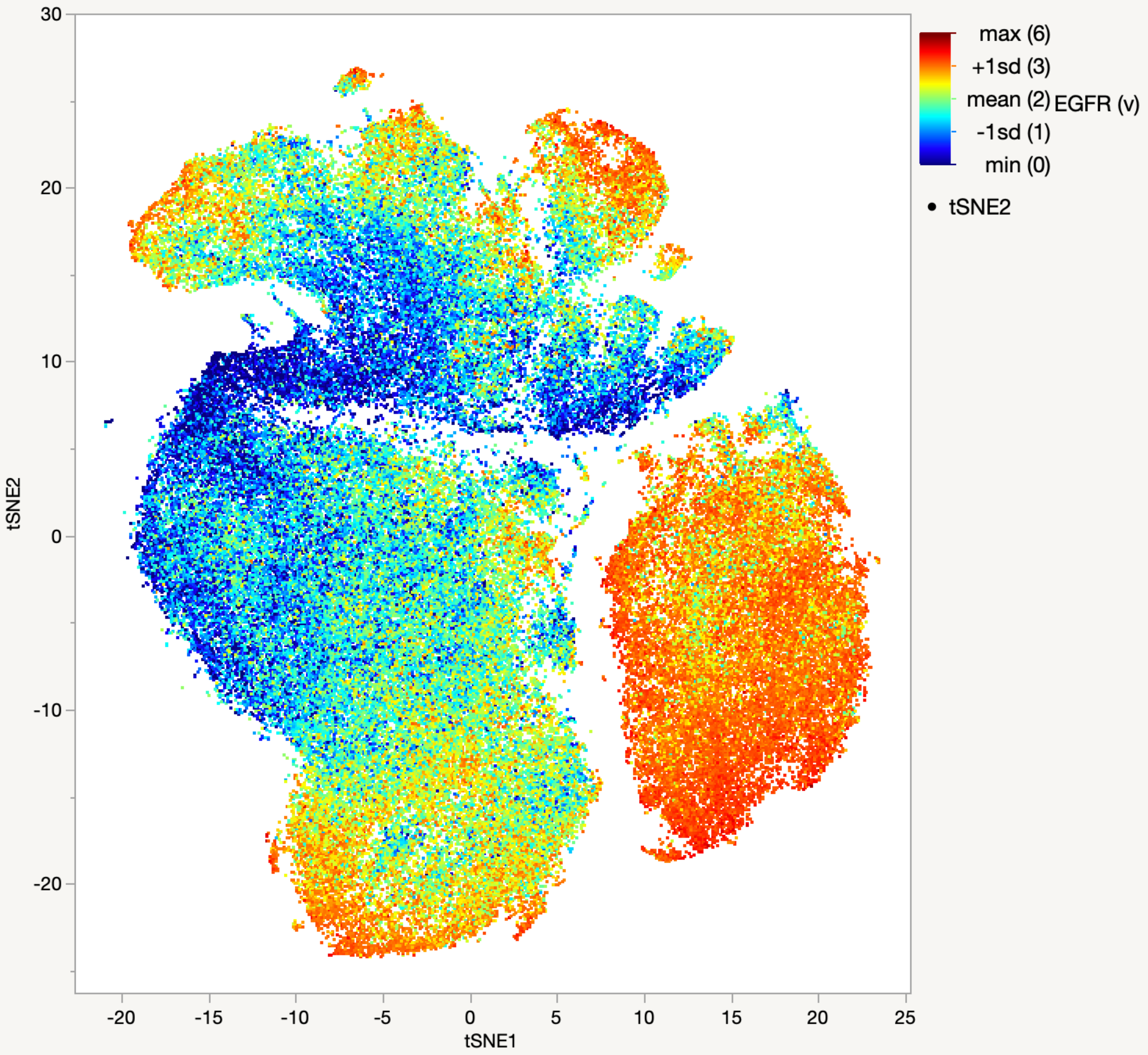

max (6)
+1sd (3)
mean (2)
-1sd (1)
min (0)
EGFR (v)
tSNE2
tSNE1
30
20
10
0
-10
-20
-20
-15
-10
-5
0
5
10
15
20
25

### HER2

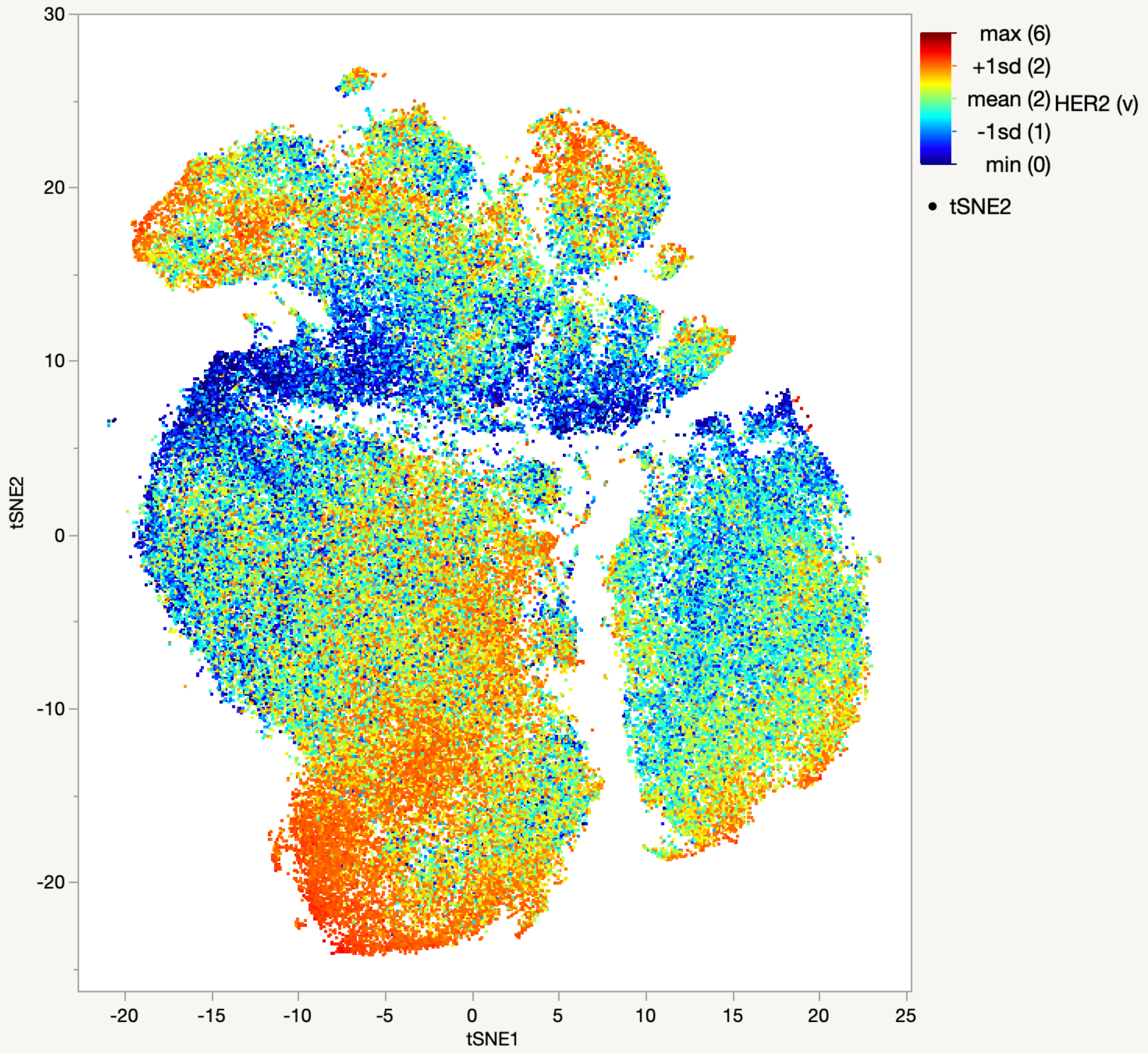

30
20
10
0
-10
-20
tSNE2
-20
-15
-10
-5
0
5
10
15
20
25
tSNE1
max (6)
+1sd (2)
mean (2)
-1sd (1)
min (0)
HER2 (v)
tSNE2

HER3

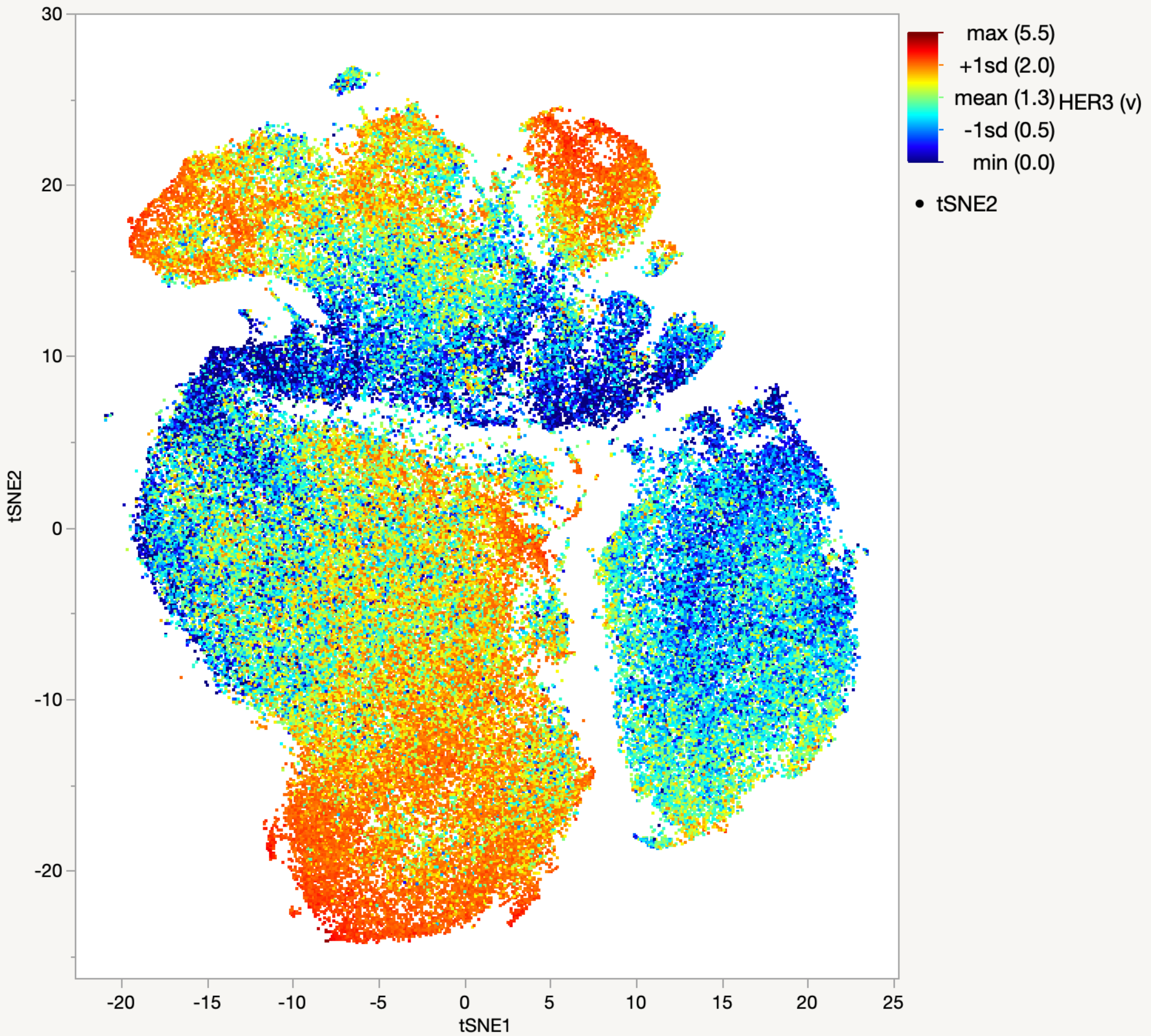

max (5.5)
+1sd (2.0)
mean (1.3)
HER3 (v)
-1sd (0.5)
min (0.0)
tSNE2
tSNE2
tSNE1
30
20
10
0
-10
-20
-20
-15
-10
-5
0
5
10
15
20
25

### IGF1R

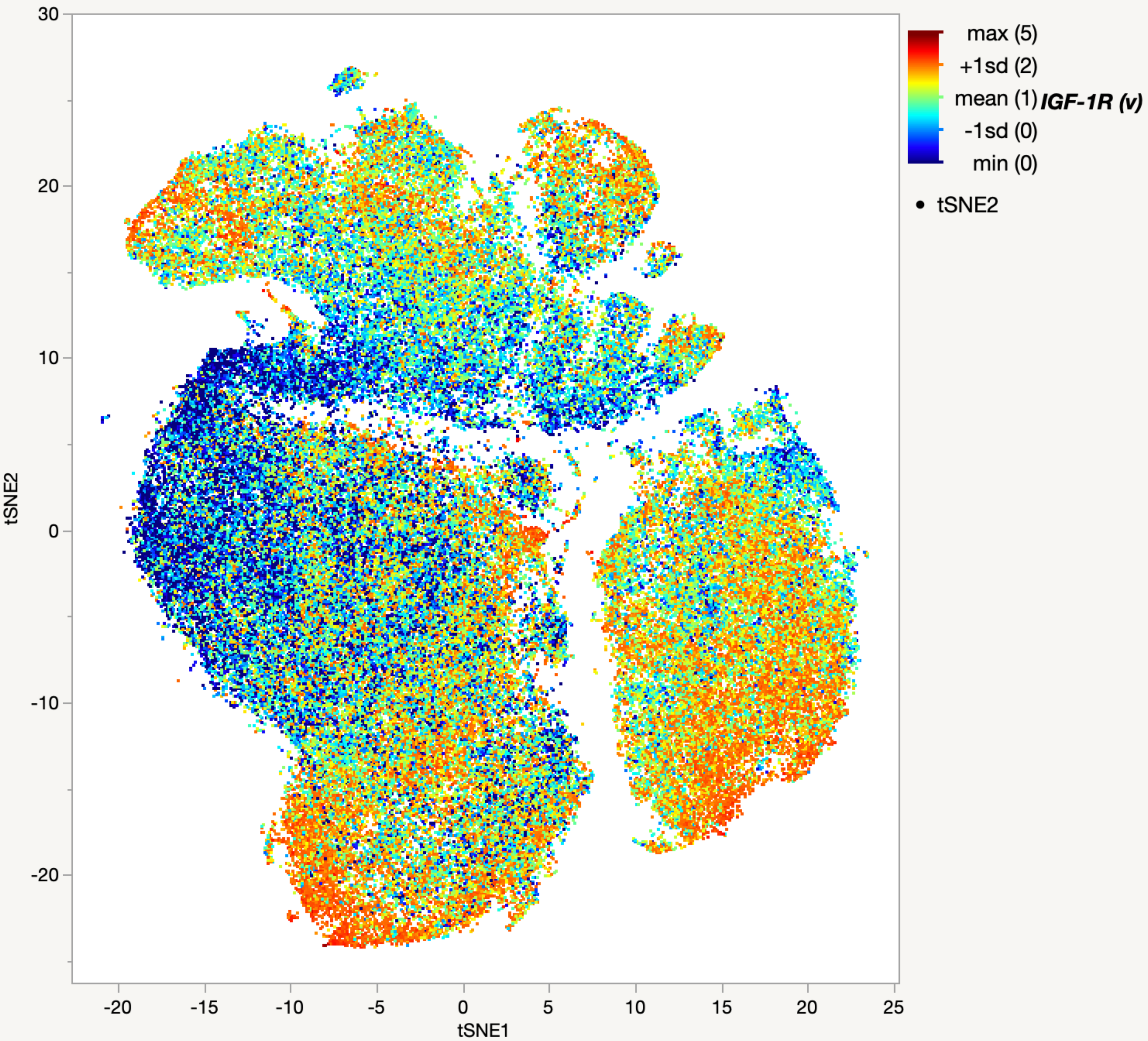

max (5)
+1sd (2)
mean (1)
IGF-1R (v)
-1sd (0)
min (0)
tSNE2
tSNE2
tSNE1
30
20
10
0
-10
-20
-20
-15
-10
-5
0
5
10
15
20
25

# Ki-67

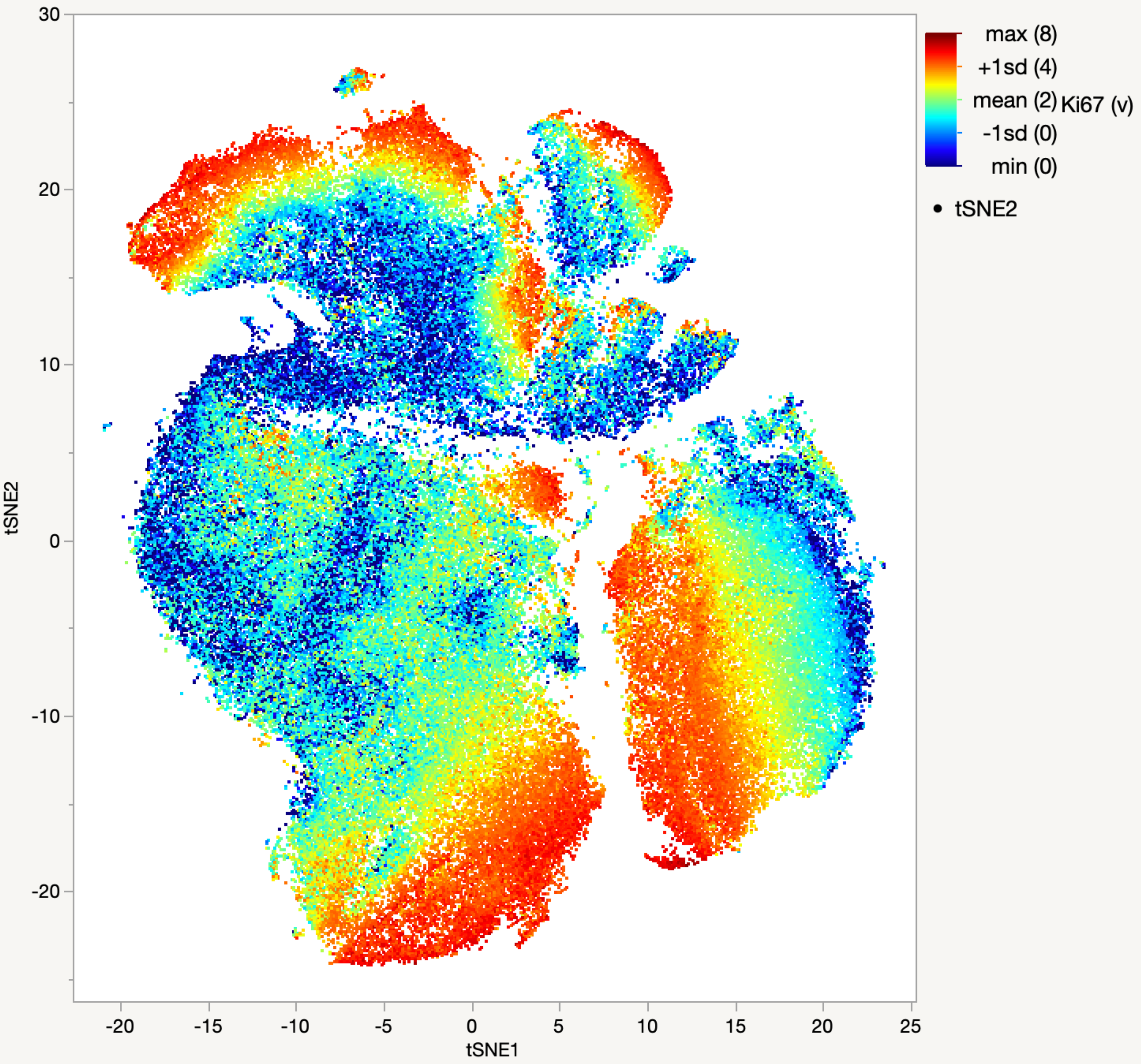

30
20
10
0
-10
-20
tSNE2
-20
-15
-10
-5
0
5
10
15
20
25
tSNE1
max (8)
+1sd (4)
mean (2)
-1sd (0)
min (0)
Ki67 (v)
tSNE2

p4EBP1

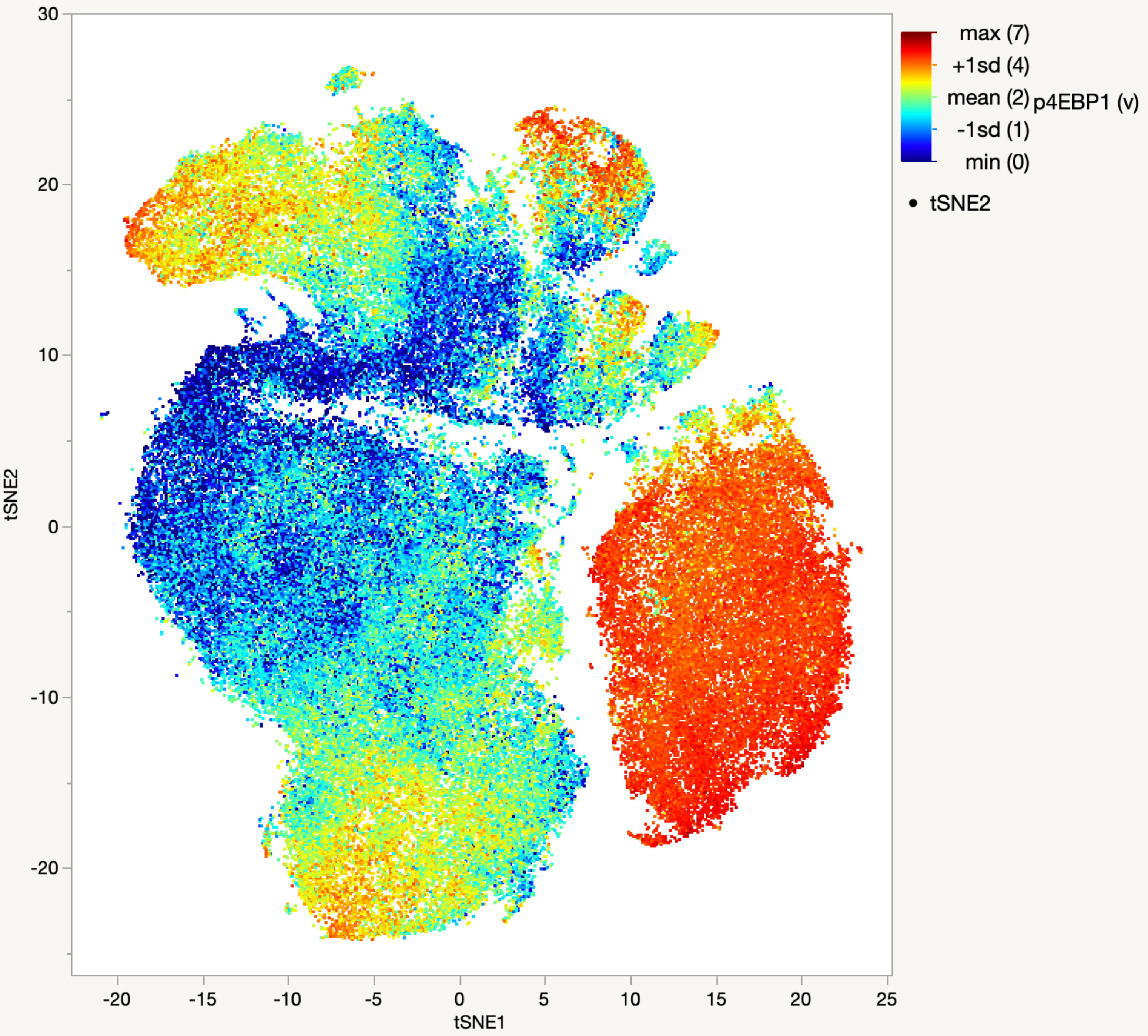

max (7)
+1sd (4)
mean (2)
-1sd (1)
min (0)
p4EBP1 (v)
tSNE2
tSNE1
30
20
10
0
-10
-20
-20
-15
-10
-5
0
5
10
15
20
25

### pAkt308

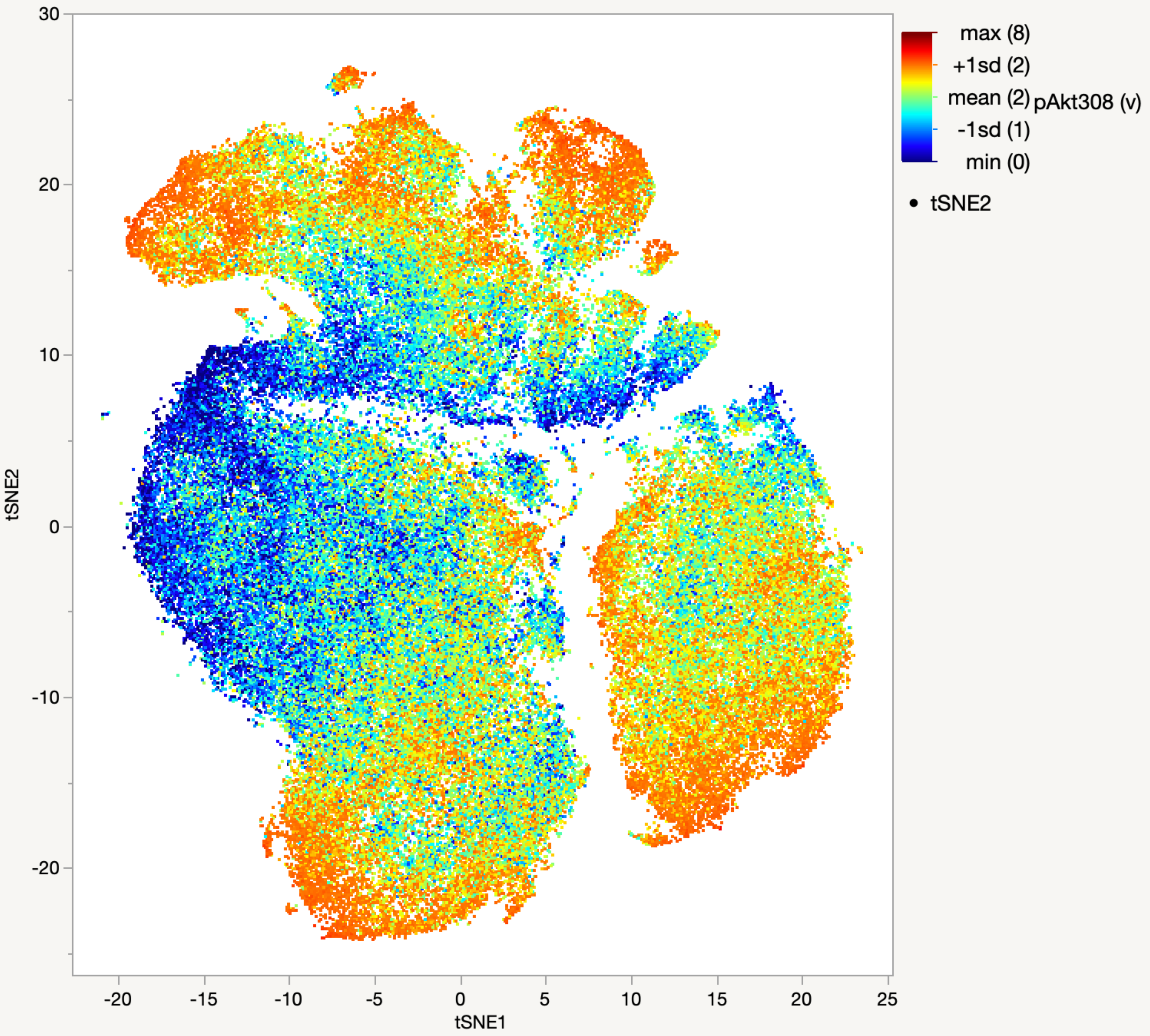

max (8)
+1sd (2)
mean (2)
-1sd (1)
min (0)
pAkt308 (v)
tSNE2
tSNE1
30
20
10
0
-10
-20
-20
-15
-10
-5
0
5
10
15
20
25

### pAkt473

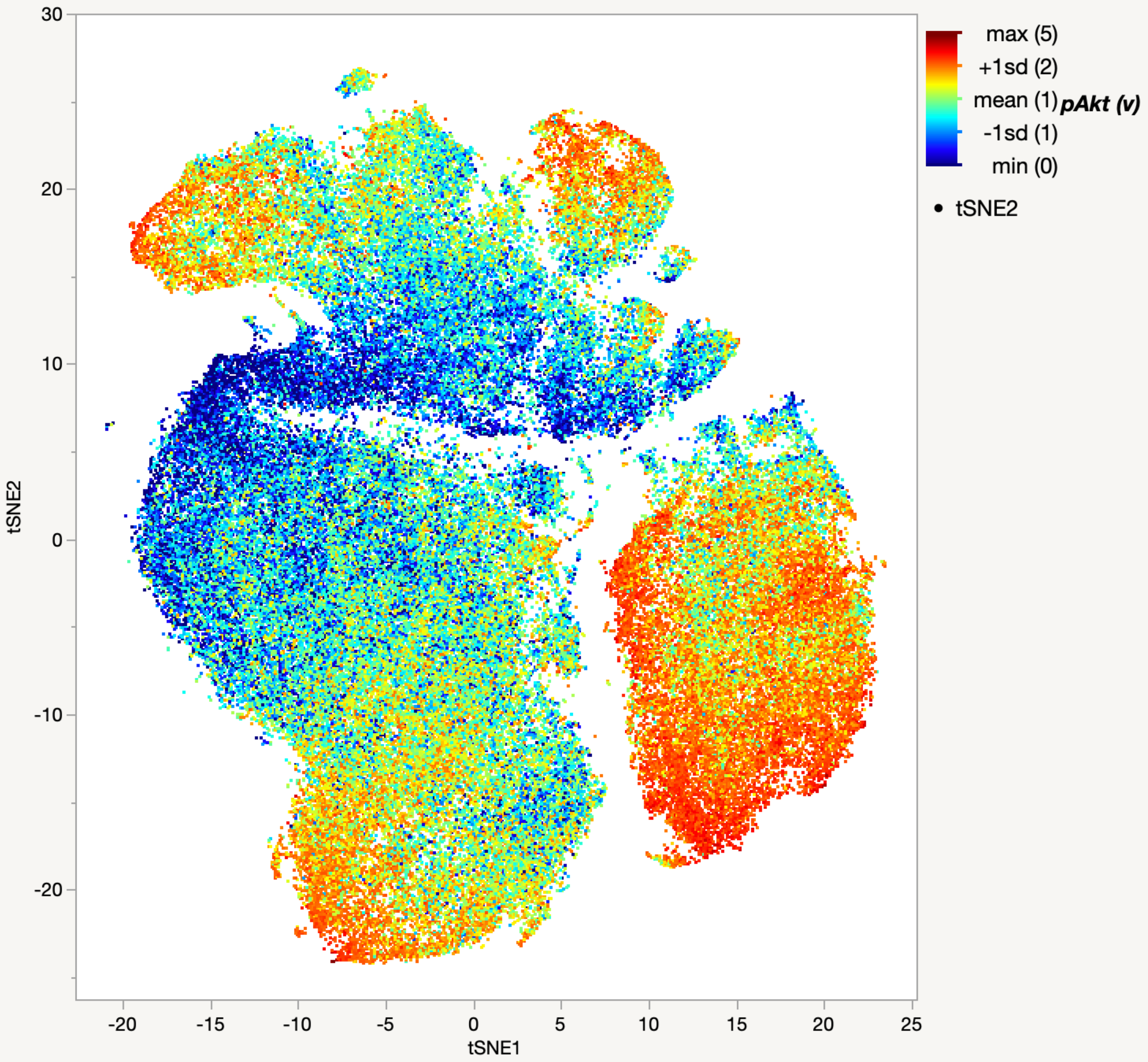

max (5)
+1sd (2)
mean (1)
-1sd (1)
min (0)
pAkt (v)
tSNE2
30
20
10
0
-10
-20
tSNE2
-20
-15
-10
-5
0
5
10
15
20
25
tSNE1

pGSK3β

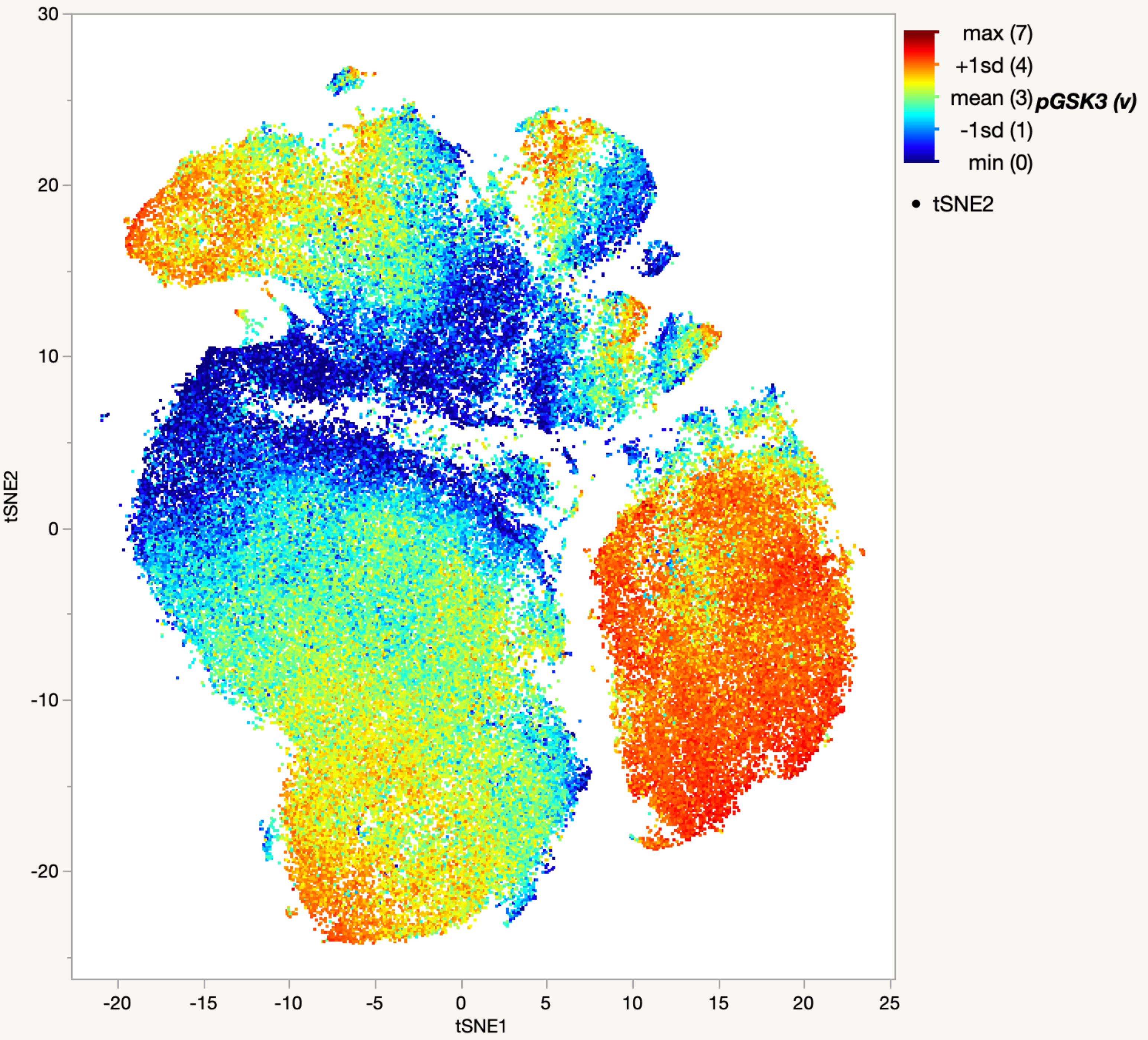

30
20
10
0
-10
-20
tSNE2
-20
-15
-10
-5
0
5
10
15
20
25
tSNE1
max (7)
+1sd (4)
mean (3)
-1sd (1)
min (0)
pGSK3 (v)
tSNE2

pS6

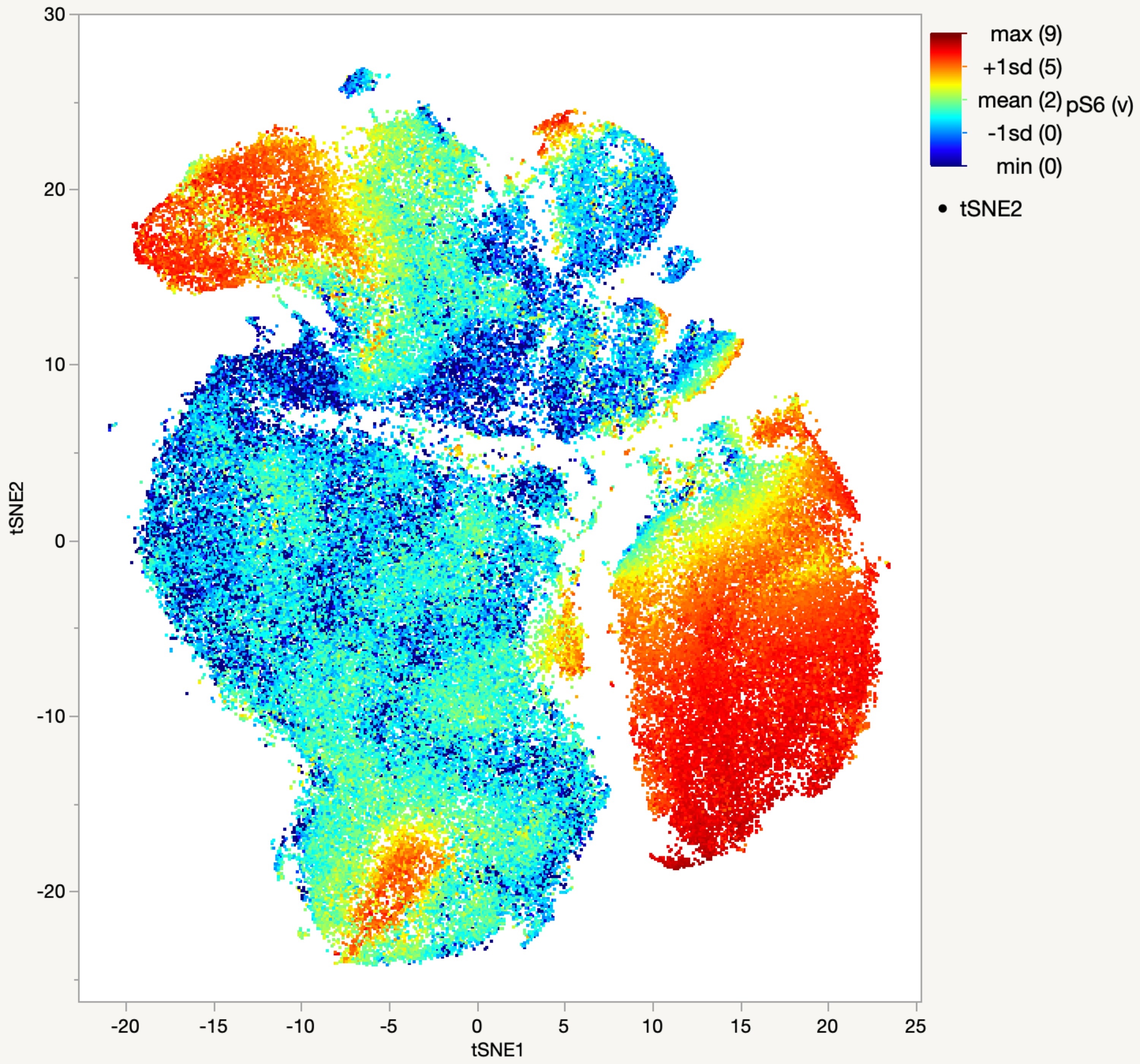

30
20
10
0
-10
-20
tSNE2
-20
-15
-10
-5
0
5
10
15
20
25
tSNE1
max (9)
+1sd (5)
mean (2)
-1sd (0)
min (0)
pS6 (v)
tSNE2

p-p90RSK

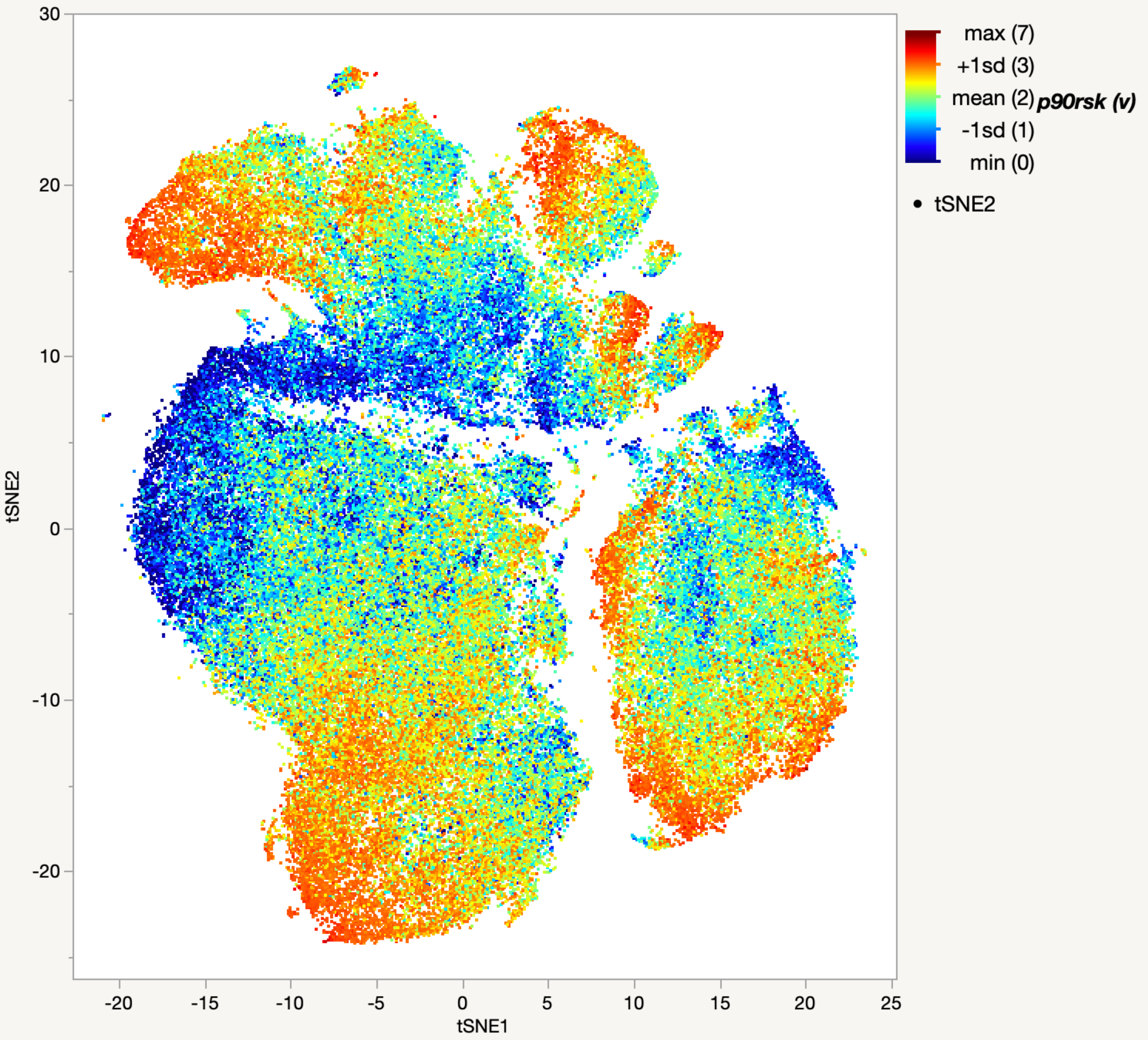

30
20
10
0
-10
-20
tSNE2
-20
-15
-10
-5
0
5
10
15
20
25
tSNE1
max (7)
+1sd (3)
mean (2)
-1sd (1)
min (0)
p90rsk (v)
tSNE2

p-p38

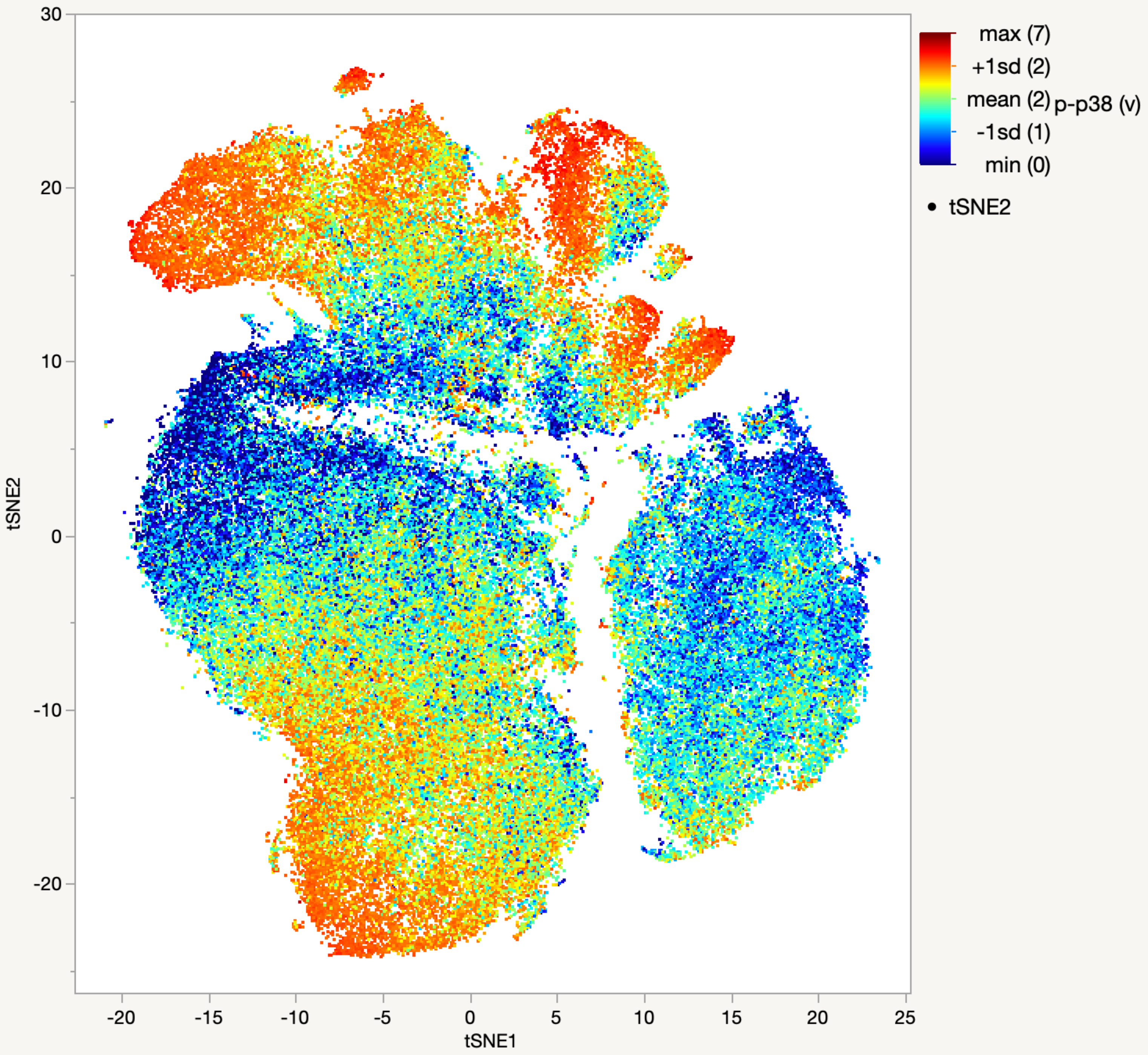

max (7)
+1sd (2)
mean (2)
-1sd (1)
min (0)
p-p38 (v)
• tSNE2
tSNE2
tSNE1
30
20
10
0
-10
-20
-20
-15
-10
-5
0
5
10
15
20
25

pErk

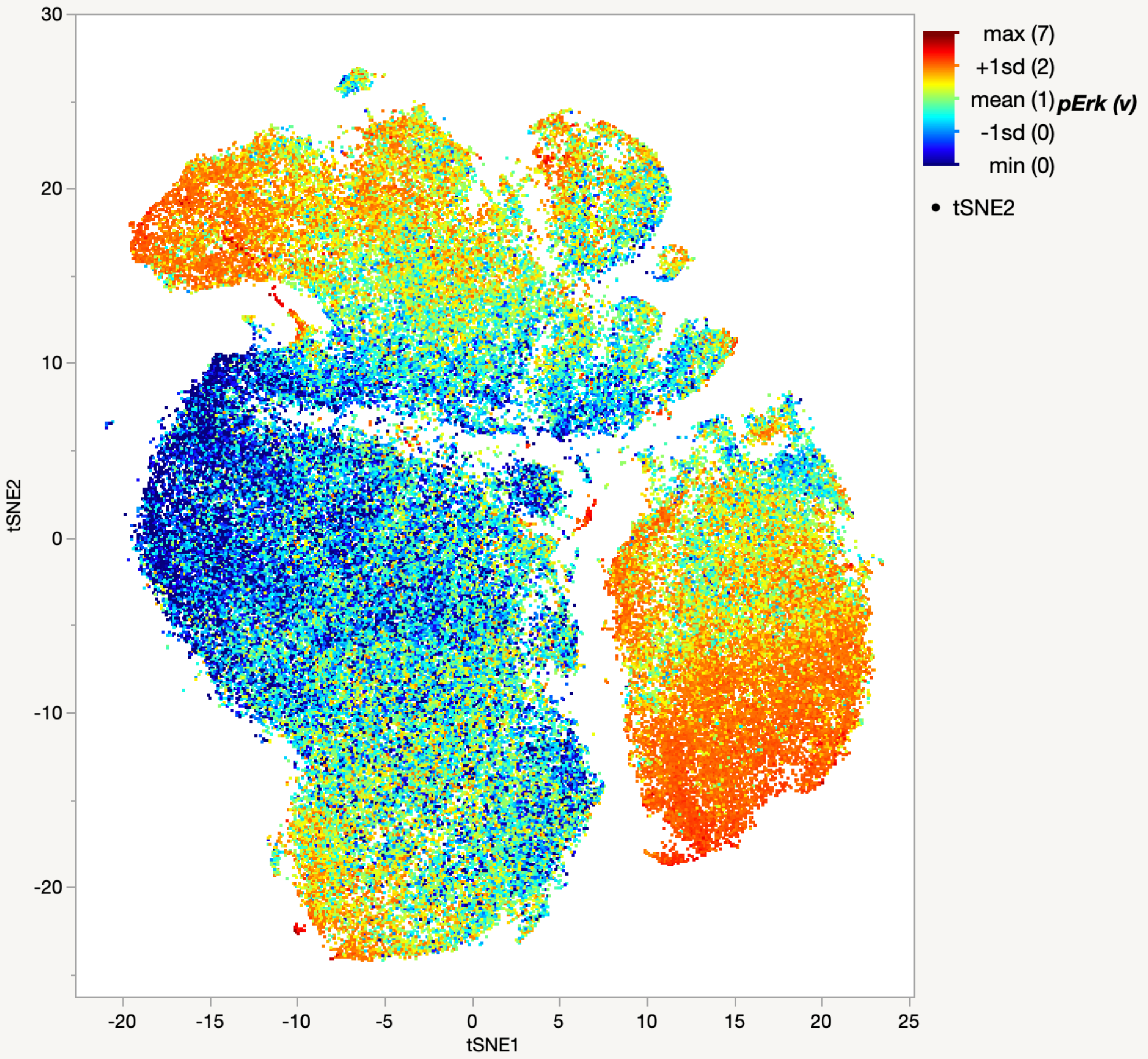

30
20
10
0
-10
-20
tSNE2
-20
-15
-10
-5
0
5
10
15
20
25
tSNE1
max (7)
+1sd (2)
mean (1)
-1sd (0)
min (0)
pErk (v)
tSNE2

pJnk

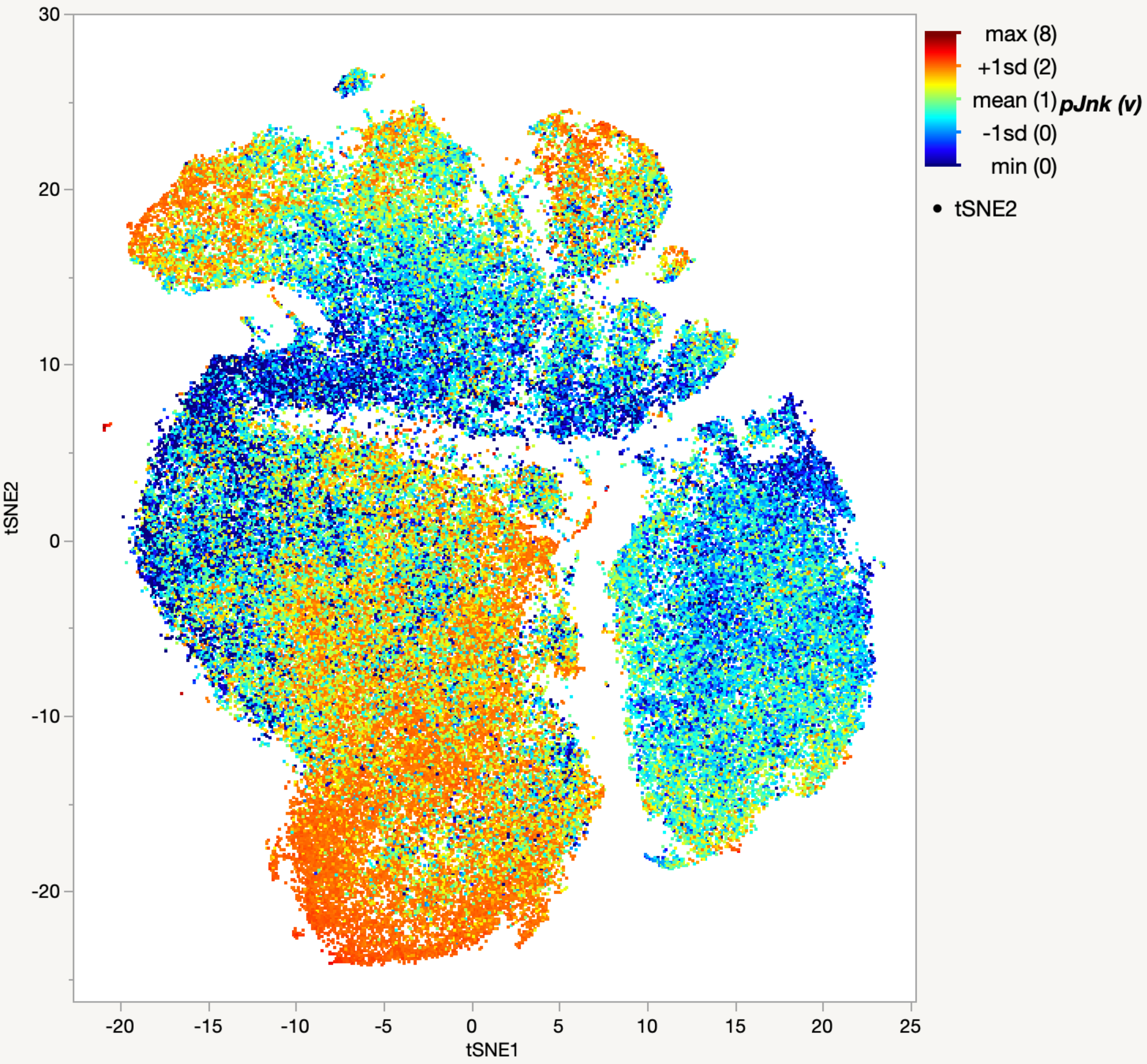

max (8)
+1sd (2)
mean (1)
-1sd (0)
min (0)
pJnk (v)
tSNE2
30
20
10
0
-10
-20
-20
-15
-10
-5
0
5
10
15
20
25
tSNE1

pStat1

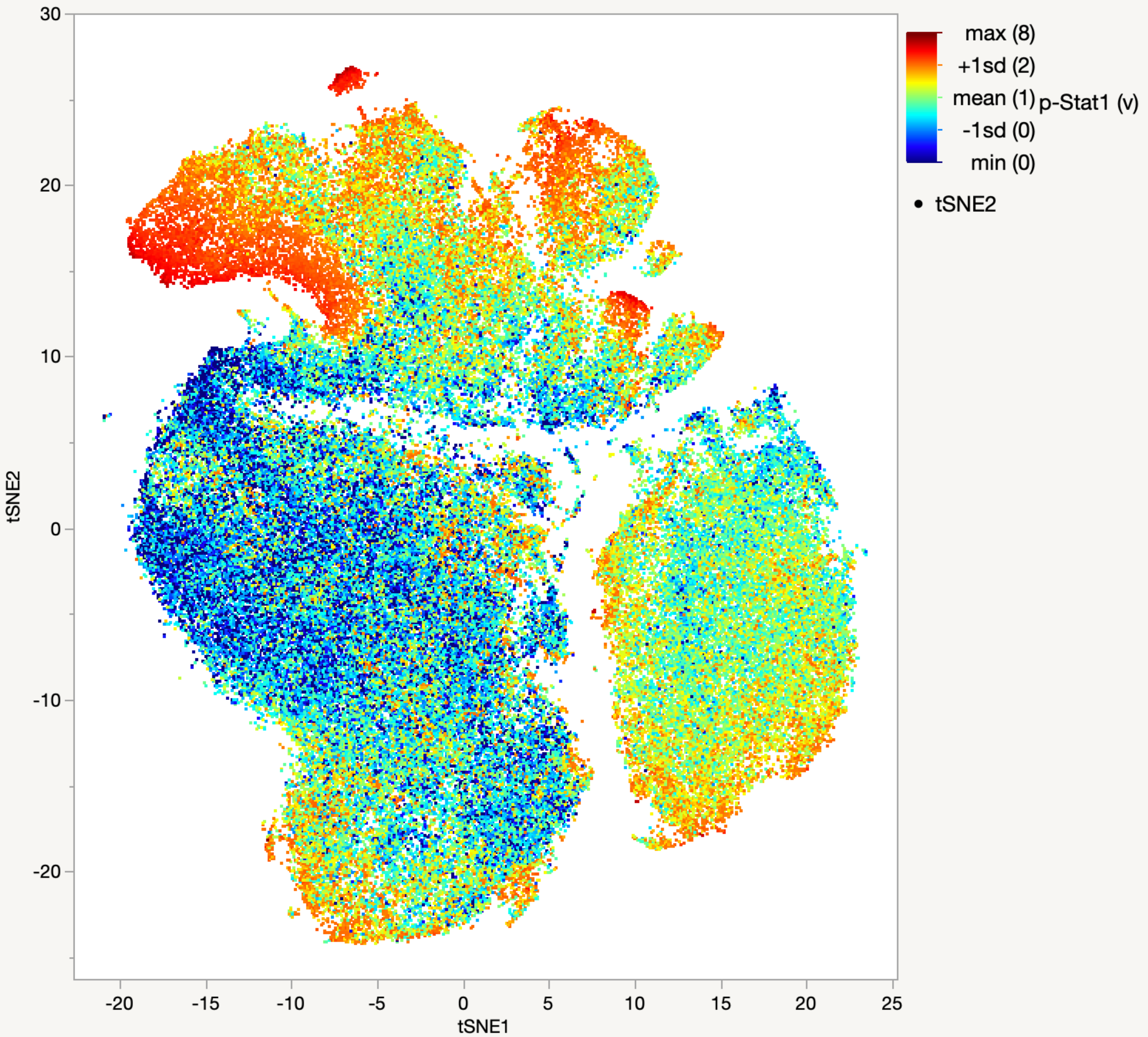

max (8)
+1sd (2)
mean (1)
-1sd (0)
min (0)
p-Stat1 (v)
tSNE2
30
20
10
0
-10
-20
-20
-15
-10
-5
0
5
10
15
20
25
tSNE1

### pStat3

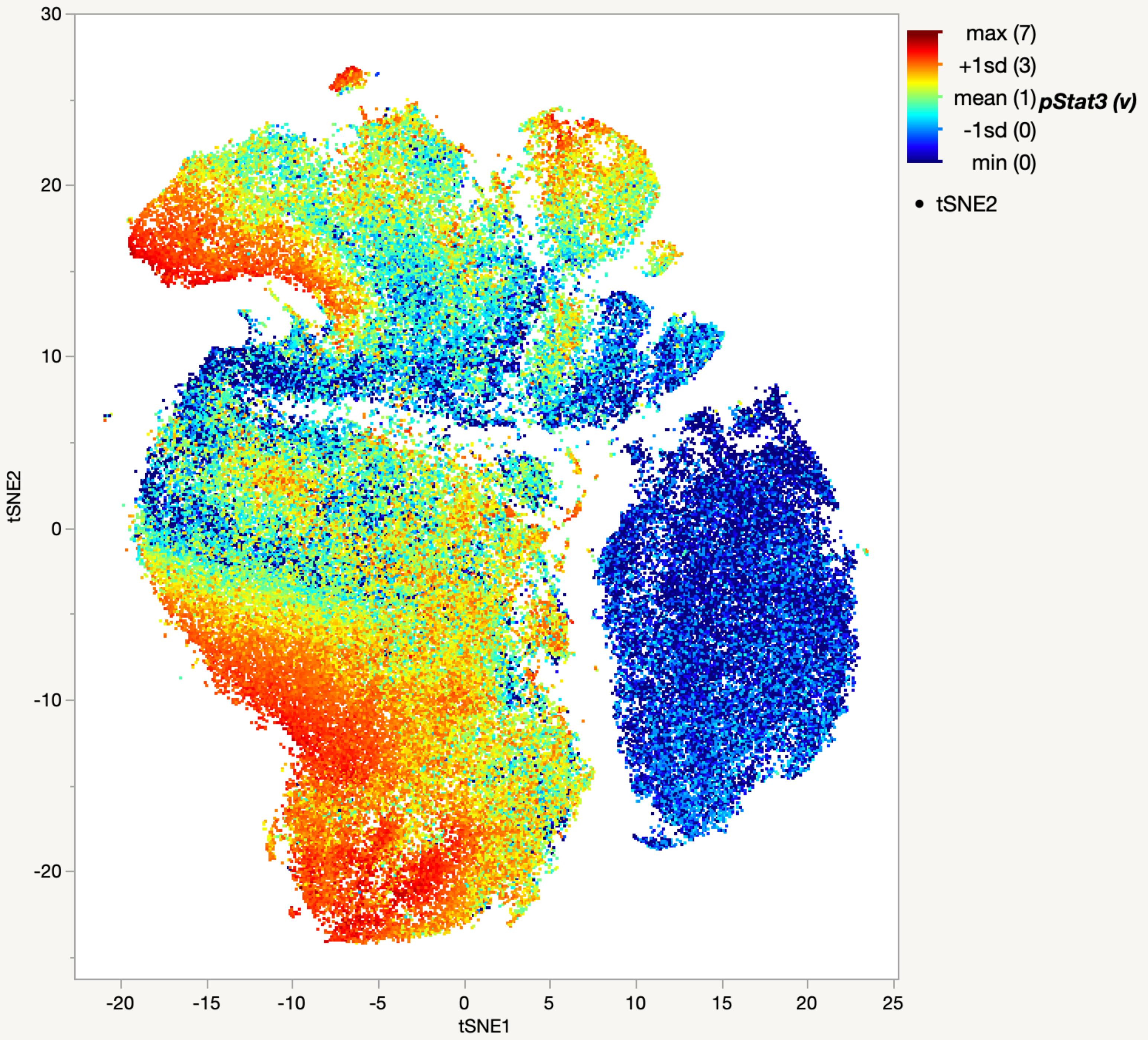

max (7)
+1sd (3)
mean (1)
-1sd (0)
min (0)
pStat3 (v)
tSNE2
30
20
10
0
-10
-20
tSNE2
-20
-15
-10
-5
0
5
10
15
20
25
tSNE1

pStat5

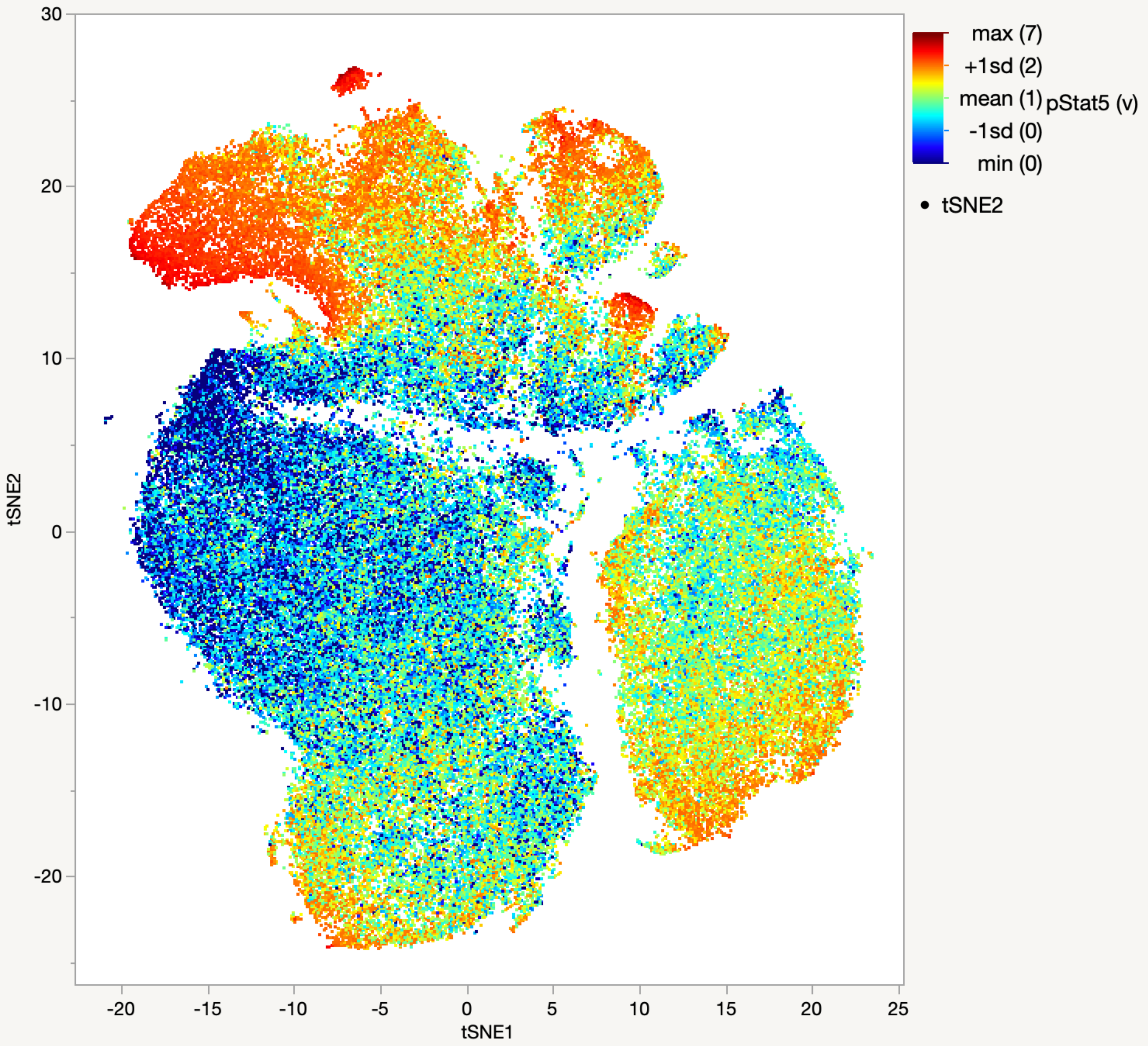
tSNE2
30
20
10
0
-10
-20
-20
-15
-10
-5
0
5
10
15
20
25
tSNE1
max (7)
+1sd (2)
mean (1)
-1sd (0)
min (0)
pStat5 (v)
tSNE2

### β-Cat

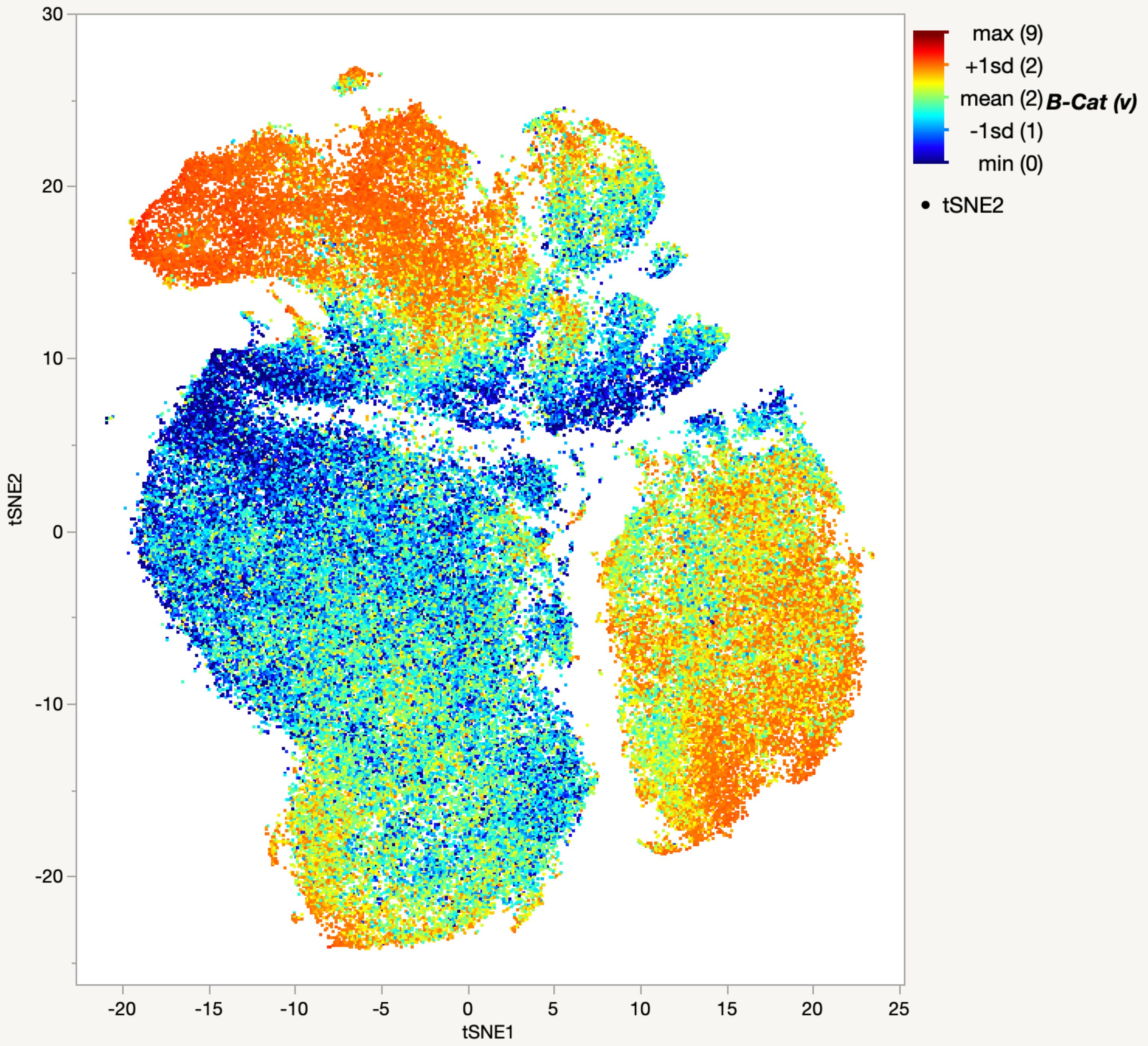

max (9)
+1sd (2)
mean (2)
B-Cat (v)
-1sd (1)
min (0)
tSNE2
tSNE1
30
20
10
0
-10
-20
-20
-15
-10
-5
0
5
10
15
20
25

### c-Myc

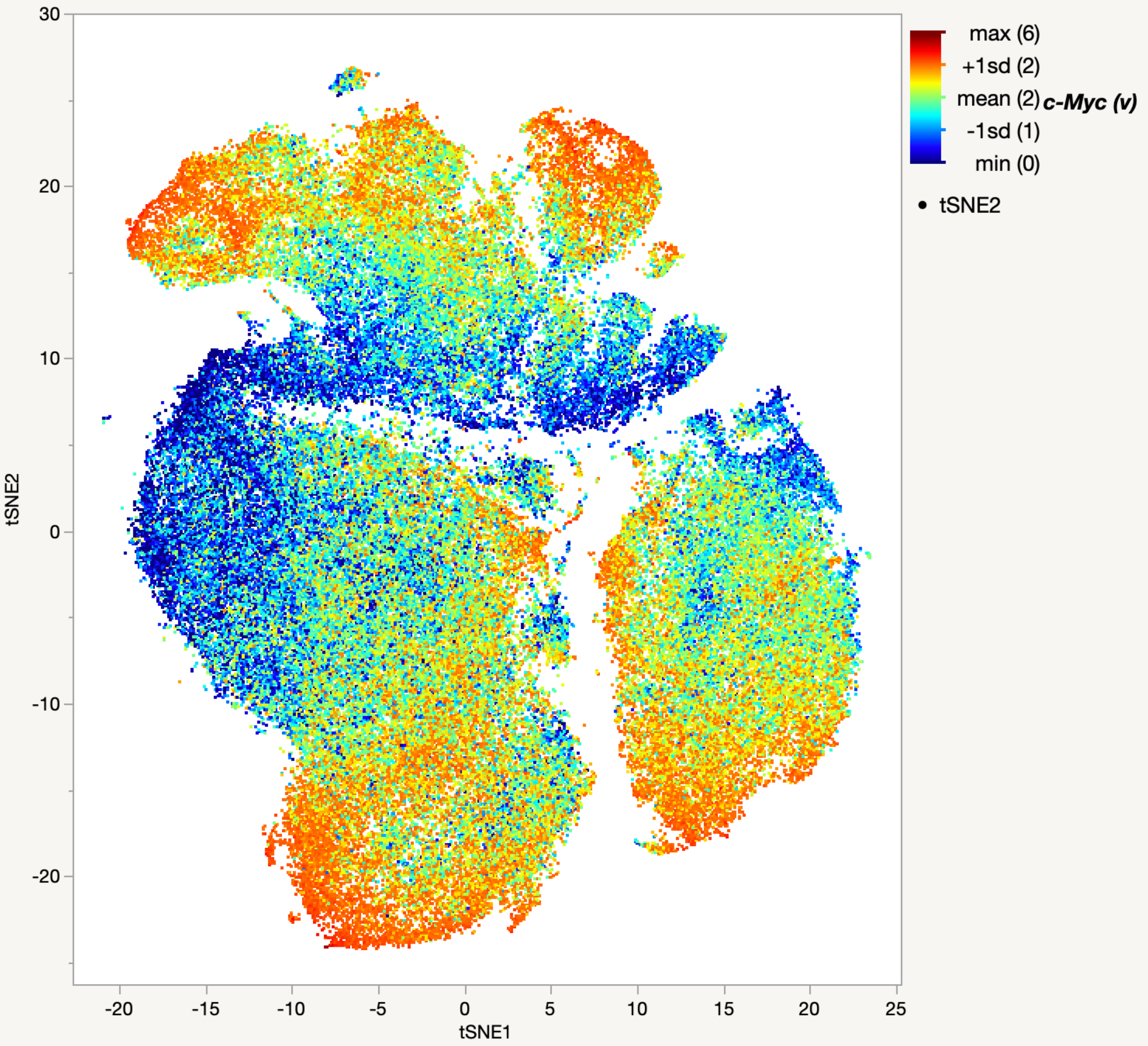

30
20
10
0
-10
-20
tSNE2
-20
-15
-10
-5
0
5
10
15
20
25
tSNE1
max (6)
+1sd (2)
mean (2)
-1sd (1)
min (0)
c-Myc (v)
tSNE2

# CD44

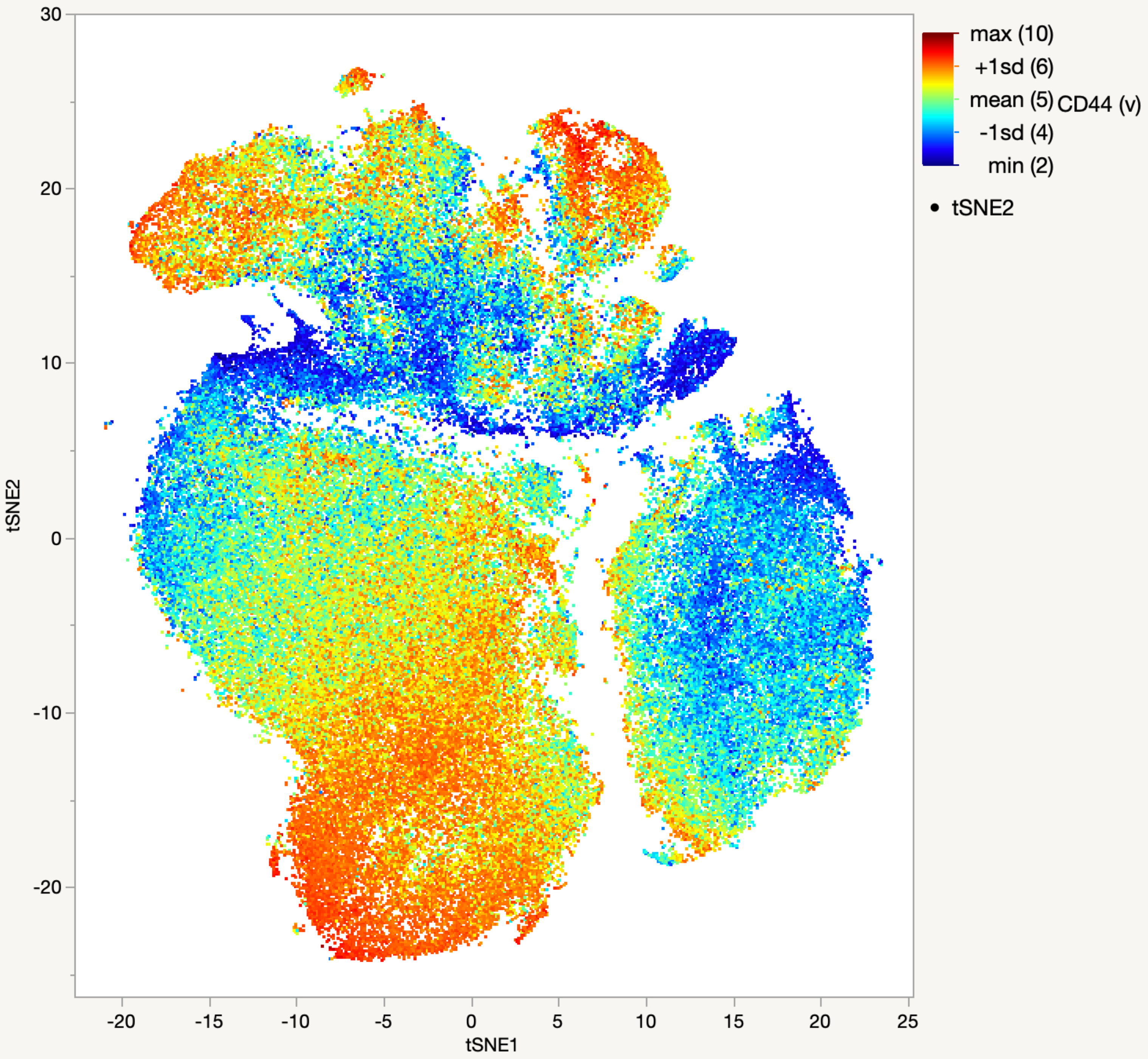
30
20
10
0
-10
-20
tSNE2
-20
-15
-10
-5
0
5
10
15
20
25
tSNE1
max (10)
+1sd (6)
mean (5)
-1sd (4)
min (2)
CD44 (v)
tSNE2

CK

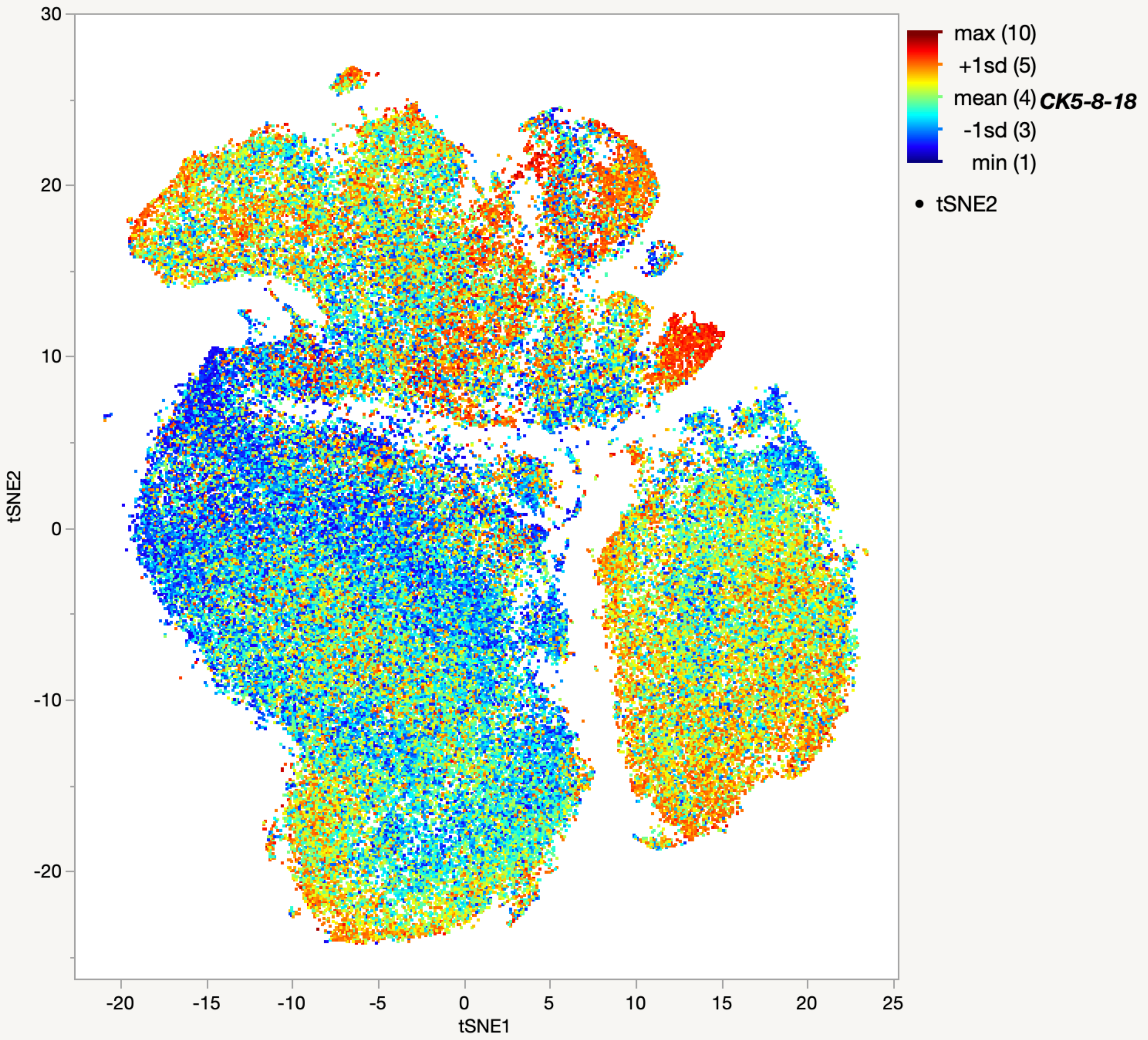

max (10)
+1sd (5)
mean (4)
CK5-8-18
-1sd (3)
min (1)
tSNE2
tSNE2
tSNE1
30
20
10
0
-10
-20
-20
-15
-10
-5
0
5
10
15
20
25

Cl-Casp3

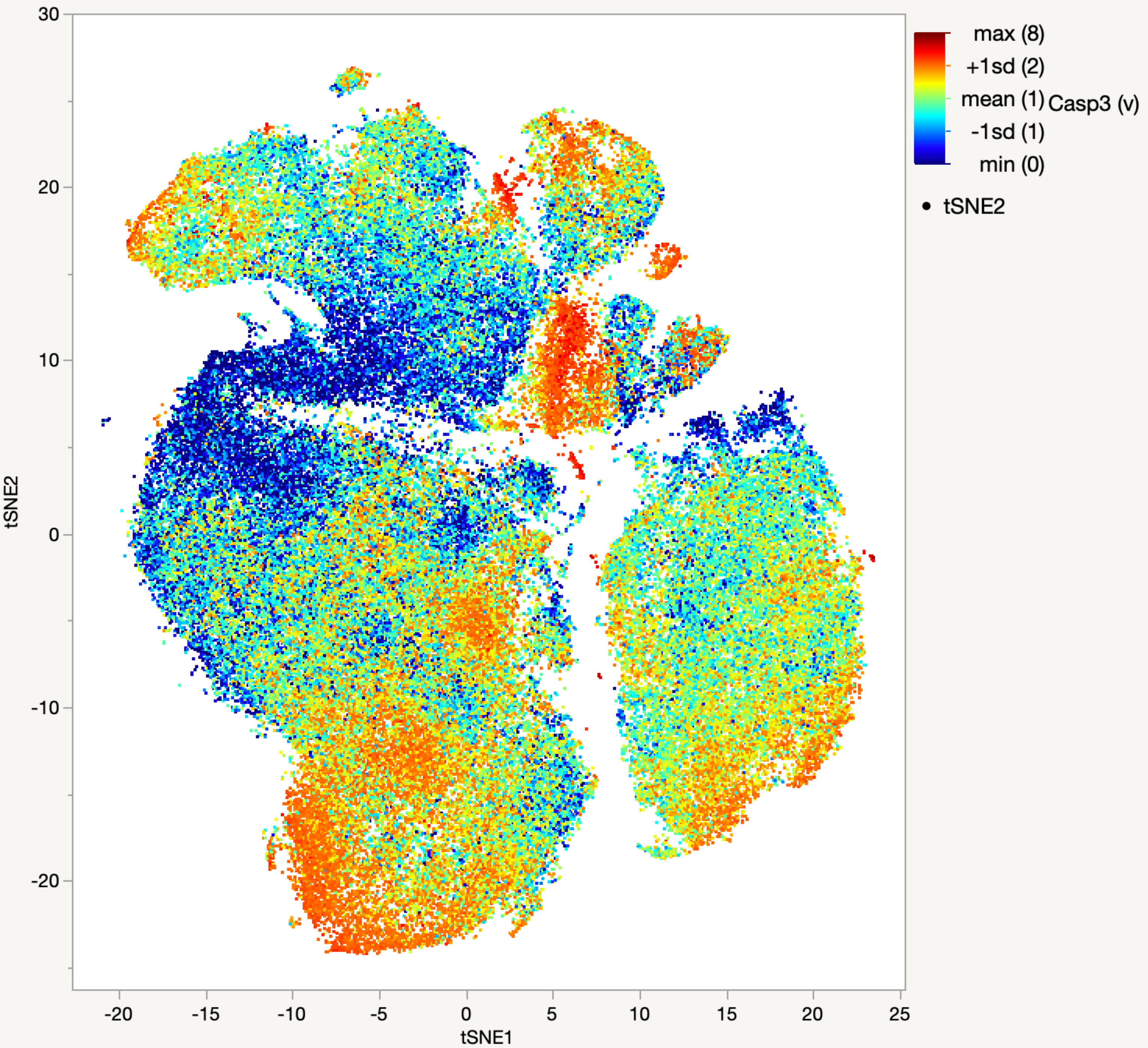

30
20
10
0
-10
-20
tSNE2
-20
-15
-10
-5
0
5
10
15
20
25
tSNE1
max (8)
+1sd (2)
mean (1)
-1sd (1)
min (0)
Casp3 (v)
tSNE2
