## Supplementary Table 1 for "Single cell proteomics of tumor compartments identifies differential kinase activities defining sensitivity to mTOR-PI3-kinase inhibition"

**SUPPLEMENTARY TABLE 1: Cell number statistics for mass cytometry datasets**

| Model | Sample Set | Mean Cell Number | Median Cell Number | Min Cell Number | Max Cell Number | Std Dev Cell Number |
|---|---|---|---|---|---|---|
| PC3 | Primary | 8998 | 2581 | 148 | 49619 | 16015.3 |
| PC3 | CTC | 89 | 44 | 15 | 314 | 100.4 |
| PC3 | Lung | 1236 | 321 | 47 | 3985 | 1544.4 |
| PC3 | Liver | 2974 | 1324 | 52 | 16806 | 5653.4 |
| PC3 | Bone | 7652 | 5134 | 141 | 17680 | 9036.5 |
| CE1-4 | Primary | 33427 | 32310 | 20304 | 50948 | 10464.7 |
| CE1-4 | CTC | 8 | 10 | 4 | 11 | 3.8 |
| CE1-4 | Lung | 141 | 165 | 12 | 247 | 119.3 |
| CE1-4 | Liver | 153 | 128 | 17 | 338 | 150.5 |
| CE1-4 | Bone | 5681 | 4998 | 20 | 12709 | 5855.5 |
