## Supplementary Table 2 for "Single cell proteomics of tumor compartments identifies differential kinase activities defining sensitivity to mTOR-PI3-kinase inhibition"

**SUPPLEMENTARY TABLE 2: Patient survival data for breast cancer dataset 2**

| | Patient | OS days | OS event | TTP days | TTP event |
|---|---|---|---|---|---|
| BR21_1_SC1_052113 | BR21 | 220 | 1 | 43 | 1 |
| BR21_1_SC2_052113 | BR21 | 220 | 1 | 43 | 1 |
| BR21_1_SC3_052113 | BR21 | 220 | 1 | 43 | 1 |
| BR29_1_012913 | BR29 | 999 | 0 | 408 | 0 |
| BR29_2_012913 | BR29 | 999 | 0 | 408 | 0 |
| BR29_3_BYL_012913 | BR29 | 998 | 0 | 407 | 0 |
| BrTr08_1_CL1_012913 | BrTr08 | 28 | 0 | 28 | 0 |
| BrTr08_1_CL2_012913 | BrTr08 | 28 | 0 | 28 | 0 |
| BrTr11_1_CL1_022513 | BrTr11 | 125 | 1 | 71 | 1 |
| BrTr11_1_CL2_022513 | BrTr11 | 125 | 1 | 71 | 1 |
| BrTr11_1_CL3_022513 | BrTr11 | 125 | 1 | 71 | 1 |
| BrTr11_1_SC2_022513 | BrTr11 | 125 | 1 | 71 | 1 |
| BrTr12_SC_022813 | BrTr12 | NA | NA | NA | NA |
| Brx03_CL1_012913 | Brx03 | NA | NA | NA | NA |
| Brx10_1_SC1_041113 | Brx10 | 621 | 1 | 29 | 1 |
| Brx10_1_SC2_041113 | Brx10 | 621 | 1 | 29 | 1 |
| Brx10_1_SC3_041113 | Brx10 | 621 | 1 | 29 | 1 |
| Brx100_1_SC3_080113 | Brx100 | 98 | 0 | 98 | 0 |
| Brx100_1_SC4_080113 | Brx100 | 98 | 0 | 98 | 0 |
| Brx100_1_SC5_080113 | Brx100 | 98 | 0 | 98 | 0 |
| Brx100_2_BYL_SC2_080213 | Brx100 | 97 | 0 | 97 | 0 |
| Brx107_1_SC2_092013 | Brx107 | 754 | 0 | 645 | 1 |
| Brx107_1_SC1_092013 | Brx107 | 754 | 0 | 645 | 1 |
| Brx109_1_SC1_121613 | Brx109 | 92 | 1 | 64 | 0 |
| Brx109_1_CL1_121613 | Brx109 | 92 | 1 | 64 | 0 |
| Brx11_CL_020413 | Brx11 | NA | NA | NA | NA |
| Brx11_SC_020413 | Brx11 | NA | NA | NA | NA |
| Brx116_1_SC1_120213 | Brx116 | 637 | 0 | 275 | 1 |
| Brx116_1_SC3_120213 | Brx116 | 637 | 0 | 275 | 1 |
| Brx116_2_SC1_120213 | Brx116 | 637 | 0 | 275 | 1 |
| Brx116_1_SC1_092414 | Brx116 | 342 | 0 | 98 | 0 |
| Brx116_2_TX_SC1_092414 | Brx116 | 342 | 0 | 98 | 0 |
| Brx117_1_SC1_120613 | Brx117 | 501 | 0 | 223 | 1 |
| Brx117_1_SC2_120613 | Brx117 | 501 | 0 | 223 | 1 |
| Brx117_2_SC1_091814 | Brx117 | 216 | 0 | 232 | 1 |
| Brx12_SC_012913 | Brx12 | NA | NA | NA | NA |
| Brx120_1_SC2_120513 | Brx120 | NA | NA | 208 | 1 |
| Brx120_1_SC3_120513 | Brx120 | NA | NA | 208 | 1 |
| Brx120_2_BYL_SC1_120613 | Brx120 | NA | NA | 207 | 1 |
| Brx121_1_SC1_120913 | Brx121 | 651 | 0 | 447 | 1 |
| Brx121_1_SC2_120913 | Brx121 | 651 | 0 | 447 | 1 |
| Brx121_1_SC3_120913 | Brx121 | 651 | 0 | 447 | 1 |
| Brx122_1_SC1_121213 | Brx122 | 636 | 0 | 165 | 1 |

| | | | | | |
|---|---|---|---|---|---|
| Brx122_1_SC2_121213 | Brx122 | 636 | 0 | 165 | 1 |
| Brx122_1_SC3_121213 | Brx122 | 636 | 0 | 165 | 1 |
| Brx126_1_SC3_021214 | Brx126 | 615 | 0 | 62 | 1 |
| Brx130_1_SC1_011714 | Brx130 | 83 | 0 | 83 | 0 |
| Brx130_1_SC2_011714 | Brx130 | 83 | 0 | 83 | 0 |
| Brx134_1_SC1_021914 | Brx134 | 593 | 0 | 53 | 1 |
| Brx134_1_SC2_021914 | Brx134 | 593 | 0 | 53 | 1 |
| Brx146_2_relapse_SC1_082114 | Brx146 | NA | NA | NA | NA |
| Brx146_2_relapse_SC2_082114 | Brx146 | NA | NA | NA | NA |
| Brx146_2_relapse_SC3_082114 | Brx146 | NA | NA | NA | NA |
| Brx163_1_SC2_090814 | Brx163 | NA | NA | 175 | 1 |
| Brx163_1_SC3_090814 | Brx163 | NA | NA | 175 | 1 |
| Brx163_1_SC4_090814 | Brx163 | NA | NA | 175 | 1 |
| Brx164_1_SC1_091814 | Brx164 | 395 | 1 | 372 | 1 |
| Brx164_1_SC2_091814 | Brx164 | 395 | 1 | 372 | 1 |
| Brx164_1_SC3_091814 | Brx164 | 395 | 1 | 372 | 1 |
| Brx164_1_SC4_091814 | Brx164 | 395 | 1 | 372 | 1 |
| Brx17_SC1_020713 | Brx17 | NA | NA | NA | NA |
| Brx17_CL2_020713 | Brx17 | NA | NA | NA | NA |
| Brx35_1_SC1_091213 | Brx35 | 567 | 1 | 56 | 1 |
| Brx35_1_SC2_110713 | Brx35 | 510 | 1 | 119 | 1 |
| Brx50_2_031113 | Brx50 | 31 | 1 | 8 | 1 |
| Brx52_CL1_020413 | Brx52 | NA | NA | NA | NA |
| Brx52_CL2_020413 | Brx52 | NA | NA | NA | NA |
| Brx52_CL3_020413 | Brx52 | NA | NA | NA | NA |
| Brx52_CL4_020413 | Brx52 | NA | NA | NA | NA |
| Brx52_CL5_020413 | Brx52 | NA | NA | NA | NA |
| Brx52_SC1_020413 | Brx52 | NA | NA | NA | NA |
| Brx52_SC2_020413 | Brx52 | NA | NA | NA | NA |
| Brx53_CL2_010313 | Brx53 | 246 | 1 | 82 | 1 |
| Brx53_SC1_010313 | Brx53 | 246 | 1 | 82 | 1 |
| Brx53_CL1_011513 | Brx53 | 219 | 1 | 55 | 1 |
| Brx53_CL2_011513 | Brx53 | 219 | 1 | 55 | 1 |
| Brx53_SC_011513 | Brx53 | 219 | 1 | 55 | 1 |
| Brx61_1_CL1_032012 | Brx61 | NA | NA | NA | NA |
| Brx61_1_CL2_032012 | Brx61 | NA | NA | NA | NA |
| Brx61_1_CL3_032012 | Brx61 | NA | NA | NA | NA |
| Brx61_1_CL4_032012 | Brx61 | NA | NA | NA | NA |
| Brx61_1_CL5_032012 | Brx61 | NA | NA | NA | NA |
| Brx61_1_CL6_032012 | Brx61 | NA | NA | NA | NA |
| Brx61_1_SC1_032012 | Brx61 | NA | NA | NA | NA |
| Brx61_1_SC2_032012 | Brx61 | NA | NA | NA | NA |
| Brx61_1_SC4_032012 | Brx61 | NA | NA | NA | NA |
| Brx66_1_012913 | Brx66 | 1007 | 0 | 1007 | 0 |
| Brx66_2_CL1_012913 | Brx66 | 1007 | 0 | 1007 | 0 |
| Brx66_2_CL2_012913 | Brx66 | 1007 | 0 | 1007 | 0 |

| | | | | | |
|---|---|---|---|---|---|
| Brx66_2_CL3_012913 | Brx66 | 1007 | 0 | 1007 | 0 |
| Brx66_2_SC1_012913 | Brx66 | 1007 | 0 | 1007 | 0 |
| Brx66_2_SC2_012913 | Brx66 | 1007 | 0 | 1007 | 0 |
| Brx70_2_BYLrelapse_SC1_092413 | Brx70 | NA | NA | 128 | 1 |
| Brx71_SC1_031413 | Brx71 | 454 | 1 | 454 | 1 |
| Brx71_SC2_031413 | Brx71 | 454 | 1 | 454 | 1 |
| Brx72_1_SC1_031813 | Brx72 | NA | NA | 67 | 1 |
| Brx72_1_SC2_031813 | Brx72 | NA | NA | 67 | 1 |
| Brx72_1_SC3_031813 | Brx72 | NA | NA | 67 | 1 |
| Brx72_1_SC4_031813 | Brx72 | NA | NA | 67 | 1 |
| Brx72_1_SC5_031813 | Brx72 | NA | NA | 67 | 1 |
| Brx72_1_SC6_031813 | Brx72 | NA | NA | 67 | 1 |
| Brx72_2_BYL_SC1_031913 | Brx72 | NA | NA | 66 | 1 |
| Brx72_2_BYL_SC2_031913 | Brx72 | NA | NA | 66 | 1 |
| Brx72_2_BYL_SC3_031913 | Brx72 | NA | NA | 66 | 1 |
| Brx73_1_SC1_032013 | Brx73 | 651 | 0 | 651 | 0 |
| Brx73_1_SC2_032013 | Brx73 | 651 | 0 | 651 | 0 |
| Brx74_1_SC2_032813 | Brx74 | 914 | 0 | 168 | 1 |
| Brx78_1_05102013 | Brx78 | 580 | 1 | 32 | 1 |
| Brx82_1_SC1_051613 | Brx82 | NA | NA | 42 | 1 |
| Brx83_1_SC1_051613 | Brx83 | 568 | 1 | 154 | 1 |
| Brx83_1_SC2_051613 | Brx83 | 568 | 1 | 154 | 1 |
| Brx83_1_SC3_051613 | Brx83 | 568 | 1 | 154 | 1 |
| Brx83_1_SC4_051613 | Brx83 | 568 | 1 | 154 | 1 |
| Brx86_1_CL1_052313 | Brx86 | 99 | 1 | 14 | 1 |
| Brx86_1_SC2_052313 | Brx86 | 99 | 1 | 14 | 1 |
| Brx86_1_SC3_052313 | Brx86 | 99 | 1 | 14 | 1 |
| Brx86_2_BYL_SC1_052413 | Brx86 | 98 | 1 | 13 | 1 |
| Brx86_2_BYL_SC2_052413 | Brx86 | 98 | 1 | 13 | 1 |
| Brx86_2_BYL_SC3_052413 | Brx86 | 98 | 1 | 13 | 1 |
| Brx87_1_SC1_062113 | Brx87 | 485 | 1 | 0 | 0 |
| Brx87_1_SC2_062113 | Brx87 | 485 | 1 | 0 | 0 |
| Brx90_1_SC1_062713 | Brx90 | 596 | 0 | 42 | 0 |
| Brx90_1_SC1_101014 | Brx90 | 126 | 0 | 116 | 1 |
| Brx90_1_SC2_101014 | Brx90 | 126 | 0 | 116 | 1 |
| Brx90_1_SC3_101014 | Brx90 | 126 | 0 | 116 | 1 |
| Brx93_1_SC1_112113 | Brx93 | NA | NA | NA | NA |
| Brx95_1_SC1_071713 | Brx95 | 835 | 0 | 835 | 0 |
| Brx95_2_BYL_CL1_071813 | Brx95 | 834 | 0 | 834 | 0 |
| Brx97_1_SC1_072313 | Brx97 | 786 | 1 | 297 | 1 |
| Brx97_1_SC2_072313 | Brx97 | 786 | 1 | 297 | 1 |
| Brx97_1_SC3_072313 | Brx97 | 786 | 1 | 297 | 1 |
| Brx97_1_SC4_072313 | Brx97 | 786 | 1 | 297 | 1 |
| Brx97_1_SC5_072313 | Brx97 | 786 | 1 | 297 | 1 |
| Brx98_1_SC3_072313 | Brx98 | 755 | 0 | 464 | 1 |
| Brx98_1_SC5_072313 | Brx98 | 755 | 0 | 464 | 1 |
