## Supplementary Table 3 for "Single cell proteomics of tumor compartments identifies differential kinase activities defining sensitivity to mTOR-PI3-kinase inhibition"

**SUPPLEMENTARY TABLE 3: Antibody signal spillover table**

| ANTIBODY | CHANNEL | | | | | | | | | | | | | | | | | | | | | | | | | | | | | | | | | |
|---|---|---|---|---|---|---|---|---|---|---|---|---|---|---|---|---|---|---|---|---|---|---|---|---|---|---|---|---|---|---|---|---|---|---|
| | Pr141 | Nd142 | Nd143 | Nd144 | Nd145 | Nd146 | Sm147 | Nd148 | Sm149 | Nd150 | Eu151 | Sm152 | Eu153 | Sm154 | Gd155 | Gd156 | Gd158 | Tb159 | Dy160 | Dy161 | Dy162 | Dy163 | Dy164 | Ho165 | Er166 | Er167 | Er168 | Tm169 | Er170 | Yb171 | Yb172 | Yb174 | Lu175 | Yb176 |
| **EpCAM** | 1 | 0.001 | 0 | 0 | 0.005 | 0.001 | 0 | 0 | 0 | 0 | 0 | 0 | 0 | 0 | 0 | 0 | 0 | 0 | 0 | 0 | 0 | 0 | 0 | 0 | 0 | 0 | 0 | 0 | 0 | 0 | 0 | 0 | 0 | 0 |
| **Cl-Casp3** | 0 | 1 | 0.004 | 0 | 0 | 0 | 0 | 0 | 0 | 0 | 0 | 0 | 0 | 0 | 0 | 0 | 0.014 | 0 | 0 | 0 | 0 | 0 | 0 | 0 | 0 | 0 | 0 | 0 | 0 | 0 | 0 | 0 | 0 | 0 |
| **ß-Cat** | 0 | 0.008 | 1 | 0.021 | 0.001 | 8E-04 | 0 | 0 | 0 | 0 | 0 | 0 | 0 | 0 | 0 | 0 | 0 | 0.012 | 0 | 0 | 0 | 0 | 0 | 0 | 0 | 0 | 0 | 0 | 0 | 0 | 0 | 0 | 0 | 0 |
| **EGFR** | 0 | 0.003 | 0.002 | 0.011 | 1 | 0.036 | 0 | 0 | 0 | 0 | 0 | 0 | 0 | 0 | 0 | 0 | 0 | 0 | 0 | 0.015 | 0 | 0 | 0 | 0 | 0 | 0 | 0 | 0 | 0 | 0 | 0 | 0 | 0 | 0 |
| **HER3** | 0 | 0 | 0 | 2E-04 | 0 | 0 | 1 | 0.023 | 0.004 | 0 | 0 | 0.002 | 0 | 0.002 | 0 | 0 | 0 | 0 | 0 | 0 | 0 | 0.003 | 0 | 0 | 0 | 0 | 0 | 0 | 0 | 0 | 0 | 0 | 0 | 0 |
| **HER2** | 0 | 0.004 | 0.001 | 0.003 | 3E-04 | 0.004 | 0 | 1 | 0 | 0.003 | 0 | 0 | 0 | 0 | 0 | 0 | 0 | 0 | 0 | 0 | 0 | 0 | 0.014 | 0 | 0 | 0 | 0 | 0 | 0 | 0 | 0 | 0 | 0 | 0 |
| **p4EBP1** | 0 | 0 | 0 | 0 | 0 | 0 | 6E-04 | 0.007 | 1 | 0.015 | 2E-04 | 0.004 | 0 | 0.003 | 0 | 0 | 0 | 0 | 0 | 0 | 0 | 0 | 0 | 0.003 | 0 | 0 | 0 | 0 | 0 | 0 | 0 | 0 | 0 | 0 |
| **pStat5** | 0 | 0.003 | 0.001 | 0.004 | 0 | 0.002 | 0 | 0.001 | 0 | 1 | 0 | 0 | 0 | 0 | 0 | 0 | 0 | 0 | 0 | 0 | 0 | 0 | 0 | 0 | 0.011 | 0 | 0 | 0 | 0 | 0 | 0 | 0 | 0 | 0 |
| **c-Met** | 0 | 0 | 0 | 0 | 0 | 0 | 0 | 0 | 0 | 0 | 1 | 5E-04 | 0.008 | 0 | 0 | 0 | 0 | 0 | 0 | 0 | 0 | 0 | 0 | 0 | 0 | 0 | 0 | 0 | 0 | 0 | 0 | 0 | 0 | 0 |
| **pAkt473** | 0 | 0 | 0 | 0 | 0 | 0 | 0 | 0 | 0 | 0 | 0 | 1 | 0 | 0.005 | 0 | 0 | 0 | 0 | 0 | 0 | 0 | 0 | 0 | 0 | 0 | 0 | 0 | 0 | 0 | 0 | 0 | 0 | 0 | 0 |
| **pStat1** | 0 | 0 | 0 | 0 | 0 | 0 | 0 | 0 | 0 | 0 | 0.006 | 0 | 1 | 1E-03 | 0 | 0 | 0 | 0 | 0 | 0 | 0 | 0 | 0 | 0 | 0 | 0 | 0 | 0 | 0 | 0 | 0 | 0 | 0 | 0 |
| **pAkt308** | 0 | 0 | 0 | 0 | 0 | 0 | 1E-04 | 0 | 1E-04 | 0 | 0 | 0.006 | 0 | 1 | 6E-04 | 0 | 0 | 0 | 0 | 0 | 0 | 0 | 0 | 0 | 0 | 0 | 0 | 0 | 0.001 | 0 | 0 | 0 | 0 | 0 |
| **pStat3** | 0 | 0 | 0 | 0 | 0 | 0 | 0 | 0 | 0 | 0 | 0 | 0 | 0 | 0 | 4E-06 | 0.001 | 1 | 0.002 | 0.006 | 0 | 0 | 0 | 0 | 0 | 0 | 0 | 0 | 0 | 0 | 0 | 0 | 0.002 | 0 | 0 |
| **c-Myc** | 0 | 0 | 0 | 0 | 0 | 0 | 0 | 0 | 0 | 0 | 0 | 0 | 0 | 0 | 0 | 0 | 0 | 1 | 0.007 | 3E-04 | 0 | 0 | 4E-06 | 0 | 0 | 0 | 0 | 0 | 0 | 0 | 0 | 0 | 0 | 0 |
| **m-CD45** | 0 | 0 | 0 | 0 | 0 | 0 | 0 | 0 | 0 | 0 | 0 | 0 | 0 | 0 | 0 | 0 | 0.005 | 0 | 1 | 0 | 0 | 0 | 0 | 0 | 0 | 0 | 0 | 0 | 0 | 0 | 0 | 0 | 0 | 0.002 |
| **pJnk** | 0 | 0 | 0 | 0 | 0 | 0 | 0 | 0 | 0 | 0 | 0 | 0 | 0 | 0 | 0 | 0 | 0 | 0 | 2E-04 | 0.015 | 1 | 0.045 | 0.007 | 0 | 0 | 0 | 0 | 0 | 0 | 0 | 0 | 0 | 0 | 0 |
| **p-p90RSK** | 0 | 0 | 0 | 0 | 0 | 0 | 0 | 0 | 0 | 0 | 0 | 0 | 0 | 0 | 0 | 0 | 0 | 0 | 0 | 3E-04 | 0.001 | 0.01 | 1 | 0.001 | 0 | 0 | 0 | 1E-04 | 5E-05 | 0 | 0 | 0 | 0 | 0 |
| **pGSK3ß** | 0 | 0 | 0 | 0 | 0 | 0 | 0 | 0 | 0 | 0 | 0 | 0 | 0 | 0 | 0 | 0 | 0 | 0 | 0 | 0 | 0 | 0 | 0 | 1 | 0.002 | 4E-05 | 0 | 0 | 0 | 0 | 0 | 0 | 0 | 0 |
| **GFP** | 0 | 0 | 0 | 0 | 0 | 0 | 0 | 0 | 0 | 0 | 0 | 0 | 0 | 0 | 0 | 0 | 0 | 0 | 0 | 0 | 0 | 0 | 0 | 0 | 0 | 0 | 2E-04 | 1 | 0.003 | 5E-05 | 0.002 | 3E-04 | 1E-04 | 0 |
| **IGF1R** | 0 | 0 | 0 | 0 | 0 | 0 | 0 | 0 | 0 | 0 | 0 | 0 | 0 | 0 | 0 | 0 | 0 | 0 | 0 | 0 | 0 | 0 | 0 | 0 | 0.004 | 0.003 | 0.006 | 4E-05 | 1 | 0.003 | 3E-04 | 0 | 4E-04 | 0 |
| **pErk** | 0 | 0 | 0 | 0 | 0 | 0 | 0 | 0 | 0 | 0 | 0 | 0 | 0 | 0 | 0 | 0 | 0 | 0 | 0 | 0 | 0 | 0 | 0 | 0 | 0 | 0 | 0 | 0 | 0.001 | 1 | 0.037 | 0.002 | 0 | 3E-04 |
| **pS6** | 0 | 0 | 0 | 0 | 0 | 0 | 0 | 0 | 0 | 0 | 0 | 0 | 0 | 0 | 0 | 0 | 0 | 0 | 0 | 0 | 0 | 0 | 0 | 0 | 0 | 0 | 0 | 0 | 0 | 0.008 | 1 | 0.01 | 0 | 6E-04 |
| **CK** | 0 | 0 | 0 | 0 | 0 | 0 | 0 | 0 | 0 | 0 | 0 | 0 | 0 | 0 | 0 | 0 | 0 | 0 | 0 | 0 | 0 | 0 | 0 | 0 | 0 | 0 | 0 | 0 | 0 | 0 | 0 | 6E-04 | 0 | 1 |
